# The Population Genetic Signature of Polygenic Local Adaptation

**DOI:** 10.1101/000026

**Authors:** Jeremy J. Berg, Graham Coop

## Abstract

Adaptation in response to selection on polygenic phenotypes occurs via subtle allele frequencies shifts at many loci. Current population genomic techniques are not well posed to identify such signals. In the past decade, detailed knowledge about the specific loci underlying polygenic traits has begun to emerge from genome-wide association studies (GWAS). Here we combine this knowledge from GWAS with robust population genetic modeling to identify traits that have undergone local adaptation. Using GWAS data, we estimate the mean additive genetic value for a give phenotype across many populations as simple weighted sums of allele frequencies. We model the expected differentiation of GWAS loci among populations under neutrality to develop simple tests of selection across an arbitrary number of populations with arbitrary population structure. To find support for the role of specific environmental variables in local adaptation we test for correlations with the estimated genetic values. We also develop a general test of local adaptation to identify overdispersion of the estimated genetic values values among populations. This test is a natural generalization of QST /FST comparisons based on GWAS predictions. Finally we lay out a framework to identify the individual populations or groups of populations that contribute to the signal of overdispersion. These tests have considerably greater power than their single locus equivalents due to the fact that they look for positive covariance between like effect alleles. We apply our tests to the human genome diversity panel dataset using GWAS data for six different traits. This analysis uncovers a number of putative signals of local adaptation, and we discuss the biological interpretation and caveats of these results.

## 1 Introduction

From the earliest days, the study of the genetic basis of evolutionary change has generated a great deal of controversy over the nature of the variation that contributes to evolution and adaptation (Pearson, 1897; Bateson, 1902; Weldon, 1902, 1903; Pearson, 1903; Darbishire, 1905). The earliest debates pitted Mendelians against biometricians, and often centered on the question of whether evolution proceeds by continuous or discontinuous changes (see Provine, 2001, for a full account). The resolution of this conflict (Yule, 1902; Weinberg, 1909; Fisher, 1918) was central to the development of early population and quantitative genetics, with both fields seeded in large part by Fisher’s (1918) demonstration that the inheritance and evolution of quantitative characters could be explained by small contributions from many independent Mendelian loci.

While still theoretically aligned (Turelli and Barton, 1990), these two fields have often been divergent in empirical practice. Evolutionary quantitative geneticists have historically focused either on mapping the genetic basis of relatively simple traits (Slate, 2004), or in the absence of any such knowledge, on understanding the evolutionary dynamics of phenotypes in response to selection over relatively short time-scales (Kingsolver *et al.*, 2001). Population geneticists, on the other hand, have focused on understanding the subtle signals left in genetic data by selection over longer time scales (Hudson *et al.*, 1987; McDonald and Kreitman, 1991; Begun and Aquadro, 1992), usually at the expense of a clear relationship between these patterns of genetic diversity and evolution at the phenotypic level. Given this still imperfect understanding, it should come as no surprise that much debate remains over the nature of the genetic variation that matters for evolution and adaptation (e.g. Orr and Coyne, 1992; Rockman, 2012).

Enabled by technological advancements allowing for the generation of vast quantities of DNA polymorphism or sequence data, population genomic techniques have revolutionized our ability to identify recent individual selective events. These approaches are typically based on identifying unusually large allele frequency changes either by observed differentiation among populations; by identifying the diversity reducing signatures of selective sweeps (Maynard Smith and Haigh, 1974; Kaplan *et al.*, 1989; Sabeti *et al.*, 2002, 2007; Bersaglieri *et al.*, 2004; Nielsen *et al.*, 2005; Wang *et al.*, 2006; Voight *et al.*, 2006; Sabeti *et al.*, 2006; Nielsen *et al.*, 2007; Fujimoto *et al.*, 2008; Pickrell *et al.*, 2009; Akey, 2009; Chen *et al.*, 2010) or by identifying large allele frequency changes along geographic or environmental gradients (Coop *et al.*, 2010; Hancock *et al.*, 2010, 2011). While the interpretation of these tests remains difficult (e.g. Coop *et al.*, 2009; Hernandez *et al.*, 2011), they provide the first insights into the mechanistic and ecological basis of adaptive evolutionary change at a genome-wide scale.

Such approaches are nonetheless limited because they rely on identifying individual loci that look unusual, and thus are only capable of identifying selection on traits where an individual locus has a large effect on fitness. When selection acts weakly on a phenotype that is underwritten by a large number of loci, the response at any given locus is expected to be modest (Pritchard *et al.*, 2010), and the signal instead manifests as a coordinated shift in allele frequency across many loci, with the phenotype increasing (decreasing) alleles all on average shifting in the same direction (Latta, 1998, 2003; Le Corre and Kremer, 2003, 2012; Kremer and Le Corre, 2011). Because this signal is so weak at the level of the individual locus, it is impossible to identify against the genome-wide background without a very specific annotation of which sites are the target of selection on a given trait.

The advent of well-powered genome wide association studies with large sample sizes (Risch and Merikangas, 1996) has allowed for just this sort of annotation, enabling the mapping of many small effect alleles associated with phenotypic variation down to the scale of linkage disequilibrium in the population. The development and application of these methods in human populations has identified thousands of loci associated with a wide array of traits, largely confirming the polygenic view of phenotypic variation (Visscher *et al.*, 2012).

Although the field of human medical genetics has been the largest and most rapid to adopt such approaches, evolutionary geneticists studying non-human model organisms have also carried out GWAS for a wide array of fitness-associated traits, and the development of further resources is ongoing (Atwell *et al.*, 2010; Fournier-Level *et al.*, 2011; Mackay *et al.*, 2012). In human populations, the cumulative contribution of these loci to the additive variance so far only explain a fraction of the narrow sense heritability for a given trait (usually less than 15%), a phenomenon known as the missing heritability problem (Manolio *et al.*, 2009; Bloom *et al.*, 2013). Nonetheless, these GWAS hits represent a rich source of information about the loci underlying phenotypic variation.

Many researchers have begun to test whether the loci uncovered by these studies tend to be enriched for signals of selection, in the hopes of learning more about how adaptation has shaped phenotypic diversity and disease risk (e.g. Myles *et al.*, 2008; Casto and Feldman, 2011; Jostins *et al.*, 2012). The tests applied are generally still predicated on the idea of identifying individual loci that look unusual, such that a positive signal of selection is only observed if some subset of the GWAS loci have experienced strong enough selection to make them individually distinguishable from the genomic background. As noted above, it is unlikely that such a signature will exist if adaptation is truly polygenic, and thus many selective events will not be identified by this approach.

Here we develop and implement a general method based on simple quantitative and population genetic principals, using allele frequency data at GWAS loci to test for a signal of selection on the phenotypes they underwrite while accounting for the hierarchical structure among populations induced by shared history and genetic drift. Our work is most closely related to the recent work of Turchin *et al.* (2012); ? and Corona *et al.* (2013), who look for co-ordinated shifts in allele frequencies of GWAS alleles for particular traits. Our approach constitutes an improvement over the methods implemented in these studies as it provides a high powered and theoretically grounded approach to investigate selection in an arbitrary number of populations with an arbitrary relatedness structure.

Using the set of GWAS effect size estimates and genome wide allele frequency data, we estimate the mean genetic value (Fisher, 1930; Falconer, 1960) for the trait of interest in a diverse array of human populations. These genetic values may in some cases be poor predictors of the actual phenotypes for reasons we address below and in the Discussion. As such, we place little weight on their prediction ability, and focus instead on population genetic modeling of their joint distribution across populations, which provides a strong and robust way of indirectly investigating the effects of selection on the genetic basis of complex phenotypes.

We develop a framework to describe how genetic values covary across populations based on a flexible model of genetic drift and population history. Using this null model, we implement simple test statistics based on transformations of the genetic values that remove this covariance among populations. We judge the significance of the departure from neutrality by comparing to a null distribution of test statistics constructed from well matched sets of control SNPs. Specifically, we test for local adaptation by asking whether the transformed genetic values show excessive correlations with environmental or geographic variables. We also develop and implement a less powerful but more general test, which asks whether the genetic values are over-dispersed among population compared to our null model of drift. We show that this overdispersion test, which is closely related to *Q_ST_* (the phenotypic analog of *F_ST_*; Prout and Barker, 1993; Spitze, 1993), as well as Lewontin and Krakauer’s (1973) test and subsequent extensions (Bonhomme *et al.*, 2010; Günther and Coop, 2012), gains considerable power to detect selection over single locus tests by looking for unexpected covariance among loci in the deviation they take from neutral expectations. Lastly, we develop an extension of our model that allows us to identify individual populations or groups of populations whose genetic values deviate from their neutral expectations given the values observed for related populations, and thus have likely been impacted by selection.

## 2 Results

### Estimating Genetic Values with GWAS Data

Consider a trait of interest where *L* loci (SNPs) have been identified from a genome-wide association study. We arbitrarily label the alleles *A*_1_ and *A*_2_ at each locus. These loci have additive effect size estimates *α*_1_,…*α_L_*, where *α*_ℓ_ is the average effect on an individual’s phenotype of replacing an *A*_2_ allele with an *A*_1_ allele at locus *ℓ*. We have allele frequency data for *M* populations at our *L* SNPs, and denote by *p_mℓ_* the observed sample frequency of allele *A*_1_ at the *ℓ^th^* locus in the *m^th^* population. From these, we estimate the mean genetic value in the *m^th^* population as

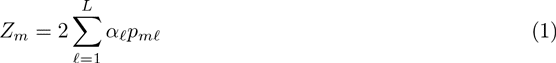

and we take 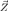 to be the vector containing the mean genetic values for all *M* populations.

The relationship between our estimated genetic values and true phenotypic values is imperfect, and thus it is worth taking a moment to cover these differences:

1. We assume that the *L* loci act strictly in an additive manner, i.e. no dominance or epistasis. This is likely to be a reasonable assumption for small effect loci, as typically found by GWASs (Hill *et al.*, 2008; Crow, 2010). However, if this assumption is violated, differences in allele frequencies across populations will result in differences in the additive effect size, and may lead to inconsistencies between estimated genetic values and observed phenotypes.
2. For the majority of GWAS associations we do not know the causal variant(s) at a locus, but rather a SNP that is in linkage disequilibrium (LD) with the causal variant(s). As the structure of LD may vary between populations, our estimates will be better in some populations than in others.
3. We assume that the effect sizes are perfectly estimated, when they may in fact by biased upward or downward due to the winner’s curse (Zollner and Pritchard, 2007) or incomplete LD (Spencer *et al.*, 2011) respectively.
4. The genetic values strictly apply only to the environment of the population in which the association study was done. They will only be representative of the phenotypes of the populations in their “natural” environment to the extent that environmental differences between populations are unimportant.
5. We do not have all of the causal variants in the population that the GWAS was performed in, and we may be missing other loci that contribute to variation in other populations but not the one in which the GWAS was done.

These limitations may result in inconsistencies between our estimated genetic values and the true genetic values, as well as the actual observed phenotypic values. However, because the methods we develop below rely only on the rejection of a neutral null model, none of them can lead to us infer a false positive signal of selection if the mapped loci evolve neutrally. They do, however, result in a loss of statistical power, and significantly complicate the biological interpretation of a positive signal. We note that one can obtain a false positive, however, due to unaccounted for population structure in the original GWAS ascertainment panel. We return to both of these issues in greater depth in the Empirical Applications and Discussion sections.

### A Model of Genetic Value Drift

We are chiefly interested in developing a framework for testing the hypothesis that the joint distribution of 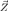 is driven by neutral processes alone, with rejection of this hypothesis implying the action of selection. We first describe a general model for the expected joint distribution of estimated genetic values 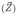 across populations under neutrality, accounting for genetic drift and shared population history.

A simple approximation to a model of genetic drift is that the current frequency of an allele in a population is normally distributed around some ancestral frequency (*ϵ*). Under a Wright-Fisher model of genetic drift, the variance of this distribution is approximately *fε*(1 − *ε*), where *f* is a property of the population shared by all loci, reflecting the compounded effect of many generations of binomially sampling (see Nicholson *et al.*, 2002, for more justification). Note also that for small values, *f* is approximately equal to the inbreeding coefficient of the present day population relative to the defined ancestral population, and thus has an interpretation as the correlation between two randomly chosen alleles relative to the ancestral population (Nicholson *et al.*, 2002).

We can expand this framework to describe the distribution of allele frequencies in an arbitrary number of populations for an arbitrary demographic history by assuming that the vector of allele frequencies in *M* populations follows a multivariate normal distribution

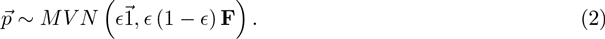

where **F** is an *M* by *M* positive definite matrix describing the correlation structure of allele frequencies across populations relative to the mean/ancestral frequency. Note again that for small values it is also approximately the matrix of inbreeding coefficients (on the diagonal) and kinship coefficients (on the off-diagonals) describing the relatedness among populations (WEIR and HILL, 2002; Bonhomme *et al.*, 2010). This flexible model was introduced, to our knowledge, by Cavalli-Sforza *et al.* (1964) (see Felsenstein, 1982, for a review), and has subsequently been used as a computationally tractable model for population history inference (e.g. Nicholson *et al.*, 2002; **?**), and as a null model for signals of selection (e.g. Coop *et al.*, 2010; Bonhomme *et al.*, 2010; Günther and Coop, 2012; Fariello *et al.*, 2013). So long as the multivariate normal assumption of drift holds reasonably well, this framework can summarize arbitrary population histories, including tree-like structures with substantial gene flow between populations (**?**), or even those which lack any coherent tree-like component (e.g. isolation by distance models: Guillot, 2012; Bradburd *et al.*, 2013).

Recall that our estimated genetic values 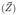 are merely a sum of sample allele frequencies weighted by effect size. If the underlying allele frequencies are well described by the multivariate normal model described above, then the distribution of 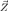 is a weighted sum of multivariate normal distributions, such that this distribution is itself multivariate normal:

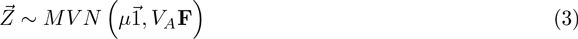

where 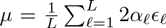 and 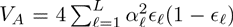 are respectively the expected phenotypic value and additive genetic variance of the ancestral (global) population. The covariance matrix describing the distribution of 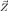 therefore differs from that describing the distribution of frequencies at individual loci only by a scaling factor that can be interpreted as the contribution of the associated loci to the additive genetic variance present in a hypothetical population with allele frequencies equal to those of the entire metapopulation.

The assumption that the drift of allele frequencies around their shared mean is normally distributed (2) may be problematic if there is substantial drift. However, even if that is the case, the estimated genetic values may still be assumed to follow a multivariate normal distribution by appealing to the central limit theorem, as each estimated genetic value is a sum over many loci. We show in the Results that this assumption often holds in practice.

Note that in light of the caveats discussed above, the elements of 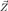 are likely to be imperfect predictions both of the true genetic values, and the observed phenotypic values. However these caveats do not invalidate our null model, as it describes only the expected allele frequency change under neutrality at the associated loci, which should be independent from their identification in a GWAS (provided that ascertainment schemes are not too complicated) and from the effect sizes estimated. As such, if the GWAS loci truly are evolving neutrally, then their evolution should be well described be our model regardless of which of the assumptions outlined above are violated.

### Fitting the Model and Standardizing the Estimated Genetic Values

As described above, we obtain the vector 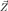 by summing allele frequencies across loci while weighting by effect size. We do not get to observe the ancestral genetic value of the sample (*µ*), so we assume that this is simply equal to the mean genetic value across populations 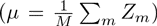. This assumption costs us a degree of freedom, and so we must work with a vector 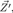 which is the vector of estimated genetic values for the first *M* − 1 populations, centered at the mean of the *M* (see Methods for details). Note that this procedure will be the norm for the rest of this paper, and thus we will always work with vectors of length *M* − 1 that are obtained by subtracting the mean of the *M* vector and dropping the last component.

To estimate the null covariance structure of the *M* − 1 populations we sample a large number K random unlinked SNPs. In our procedure, the *K* SNPs are sampled so as to match certain properties of the *L* GWAS SNPs (the specific matching procedure is described in more depth below and in the Methods section). Setting *ϵ_k_* to be the mean sample allele frequency across populations at the *k^th^* SNP, we standardize the sample allele frequency in population *m* as (*p_mk_ − ε_k_*)*/* (*ε_k_* (1 − *ε_k_*)). We then calculate the sample covariance matrix (**F**) of these standardized frequencies, accounting for the *M* − 1 rank of the matrix (see Methods). We estimate the scaling factor of this matrix **F** as

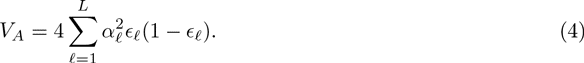

We now have an estimated genetic value for each population, and a simple null model describing their expected covariance due to shared population history. Under this multivariate normal framework, we can transform the vector of mean centered genetic values 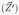 so as to remove this covariance. First, we note that the Cholesky decomposition of the **F** matrix is

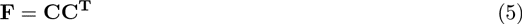

where **C** is a lower triangular matrix, and **C^T^** is its transpose. Informally, this can be thought of as taking the square root of **F**, and so **C** can be thought of as analogous to the standard deviation matrix.

Using this matrix **C** we can transform our estimated genetic values as:

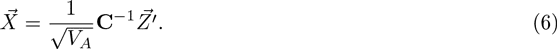

If 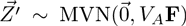 then 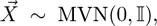 where 𝕀 is the identity matrix. Therefore, under the assumptions of our model, these standardized genetic values should be independent and identically distributed ∼*N* (0, 1) random variates.

It is worth spending a moment to consider what this transformation has done to the allele frequencies at the loci underlying the estimated genetic values. As our original genetic values are written as a weighted sum of allele frequencies, our transformed genetic values can be written as a weighted sum of transformed allele frequencies (which have passed through the same transform). We can write

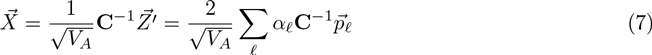

and so we can simply define the vector of transformed allele frequencies at locus *ℓ* to be

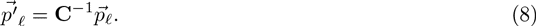

This set of transformed frequencies exist within a set of transformed populations, which by definition have zero covariance with one another under the null, and are related by a star-like population tree with branches of equal length.

As such, we can proceed with simple, straightforward and familiar statistical approaches to test for the impact of spatially varying selection on the estimated genetic values. Below we describe three simple methods for identifying the signature of polygenic adaptation, which arise naturally from this observation.

### Environmental correlations

Assume we have a vector 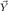, containing measurements of a specific environmental variable of interest in each of the *M* populations. We mean-center this vector and put it through a transform identical to that which we applied to the estimated genetic values in (7). This gives us a vector 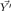, which is in the same frame of reference as the transformed genetic values.

There are many possible models to describe the relationship between a trait of interest and a particular environmental variable that may act as a selective agent. We first consider a simple linear model, where we model the distribution of transformed genetic values 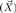 as a linear effect of the transformed environmental variables 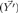

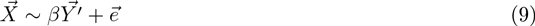

where 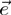 under our null is a set of normal, independent and identically distributed random variates (i.e. residuals), and we can estimate *β* by finding the value that minimizes the variance of 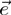 (i.e. ordinary least squares regression). We can also calculate the associated squared Pearson correlation coefficient (*r*^2^) as a measure of the fraction of variance explained by our variable of choice, as well as the non-parametric Spearman’s rank correlation 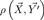, which is robust to outliers that can mislead the linear model. In the Methods we show that the linear environmental model applied to our transformed genetic values has a natural interpretation in terms of the underlying individual loci. Therefore, exploring the environmental correlations of estimated genetic values nicely summarizes information in a sensible way at the underlying loci identified by the GWAS.

In order to assess the significance of these measures, we implement an empirical null hypothesis testing framework, using *β*, *r*^2^, and *ρ* as test statistics. We sample many sets of *L* SNPs randomly from the genome, again applying a matching procedure discussed below and in the Methods. With each set of *L* SNPs we construct a vector 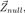, which represents a single draw from the genome-wide null distribution for a trait with the given ascertainment profile. We then perform an identical set of transformations and analyses on each 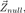, thus obtaining an empirical genome-wide null distribution for all test statistics.

### Excess Variance Test

As an alternative to testing the hypothesis of an effect by a specific environmental variable, one might simply test whether the estimated genetic values exhibit more variance among populations than expected due to drift. Here we develop a simple test of this hypothesis.

As 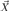 is composed of *M* − 1 independent, identically distributed standard normal random variables, a natural choice of test statistic is given by

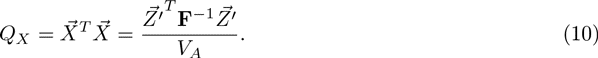

This *Q_X_* statistic represents a standardized measure of the among population variance in estimated genetic values that is not explained by drift and shared history. It is also worth noting that by comparing the rightmost form in (10) to the multivariate normal likelihood function, we find that *Q_X_* is proportional to the negative log likelihood of the estimated genetic values under the model, and is thus the natural measurement of the model’s ability to explain their distribution. Values of this statistic that are in the upper tail correspond to an excess of variance among populations (i.e. local adaptation), while values in the lower tail correspond a paucity of variance, and thus potentially to stabilizing selection. Multivariate normal theory predicts that this statistic should follow a *χ*^2^ distribution with *M* − 1 degrees of freedom under the null hypothesis. Nonetheless, we use a similar approach to that described for the linear model, generating the empirical null distribution by resampling SNPs genome-wide. As discussed below, we find that in practice the empirical null distribution tends to be very closely matched by the theoretically predicted 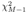 distribution.

#### The Relationship of *Q_X_* to Previous Tests

Our *Q_X_* statistic is closely related to *Q_ST_*, the phenotypic analog of *F_ST_*, which measures the fraction of the genetic variance that is among populations relative to the total genetic variance (Wright, 1951; Spitze, 1993; Prout and Barker, 1993). *Q_ST_* is typically estimated in traditional local adaptation studies via careful measurement of phenotypes from related individuals in multiple populations in a common garden setting. If the loci underlying the trait act in a purely additive manner and are experiencing only neutral genetic drift, then 𝔼[*Q_ST_*] = 𝔼[*F_ST_*] (Lande, 1992; Whitlock, 1999).

If both quantities are well estimated, and we also assume that there is no hierarchical structure among the populations, then 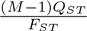 is known to have a 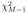 distribution under a wide range of models (Rogers and Harpending, 1983; Whitlock, 2008; Whitlock and Guillaume, 2009). Thus the statistic 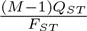 is a natural phenotypic extension of the Lewontin and Krakauer (1973) *F_ST_* based-test (LK test).

To see the close correspondence between *Q_X_* and *Q_ST_*, consider the case of a starlike population tree with branches of equal length (i.e. *f_mm_* = *F_ST_* and *f_m≠n_* = 0). Under this demographic model, we have

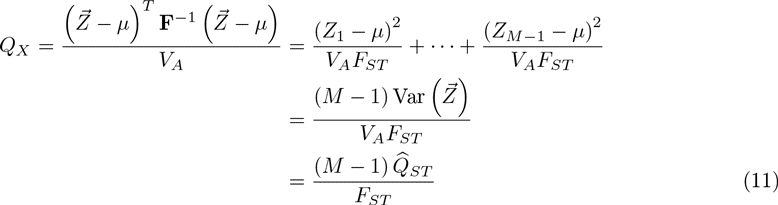

where 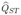 is an estimated value for *Q_ST_* obtained from our estimated genetic values. This relationship between *Q_X_* and *Q_ST_* breaks down when some pairs of populations do not have zero covariance in allele frequencies under the null, in which case the *χ*^2^ distribution of the LK test also breaks down (Robertson, 1975; Nei and Maruyama, 1975). Bonhomme *et al.* (2010) recently proposed an extension to the LK test that accounts for a population tree, thereby recovering the *χ*^2^ distribution (see also Günther and Coop, 2012, who relax the tree-like assumption), and our *Q_X_* statistic is the natural phenotypic extension of this enhanced statistic.

Given that our estimated genetic values are simple linear sums of allele frequencies, it is natural to ask how *Q_X_* can be written in terms of these frequencies. Again, restricting ourselves to the case where **F** is diagonal, (i.e. *f_mm_* = *F_ST_* and *f_m≠n_* = 0), we can express *Q_X_* as

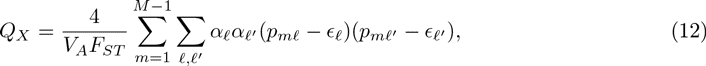

which can be rewritten as

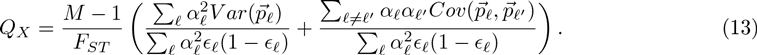

The numerator of the first term inside the parentheses is the weighted sum of the variance among populations over all GWAS loci, scaled by the contribution of those loci to the additive genetic variance in the total population. As such this first term is similar to *F_ST_* calculated for our GWAS loci, but instead of just averaging the among population and total variances equally across loci in the numerator and denominator, these quantities are weighted by the squared effect size at each locus. This weighting nicely captures the relative importance of different loci to the trait of interest.

The second term in (13) is less familiar; the numerator is the weighted sum of the covariance of allele frequencies between all pairs of GWAS loci, and the denominator is again the contribution of those loci to the additive genetic variance in the total population. This term is thus a measure of the correlation across loci in the deviation they take from the ancestral value, or the across population component of linkage disequilibrium (for a more in depth discussion of this relationship in the context of *Q_ST_*, see Latta, 1998, 2003).

Under the neutral null hypothesis, the expectation of this second term is equal to zero, as drifting loci are equally likely to covary in either direction. With differential selection among populations, however, we expect loci underlying a trait not only to vary more than we would expect under a neutral model, but also to covary in a consistent way, e.g. we expect the alleles with a positive effect on the trait to increase or decrease in concert with one another. Models of local adaptation predict that it is this covariance among alleles that is primarily responsible for differentiation at the phenotypic level (Latta, 1998, 2003; Le Corre and Kremer, 2003, 2012; Kremer and Le Corre, 2011), and as such, we expect this test statistic to offer considerably higher power than simply averaging *F_ST_* over loci, or inspecting *F_ST_* at individual GWAS loci, a point which we investigate with simulations below.

As noted above (8), when **F** is non-diagonal, our transformed genetic values can be written as a weighted sum of transformed allele frequencies. Consequently, we can obtain a similar expression to (13) when population structure exists, but now expressed in terms of the covariance of a set of allele frequencies in transformed populations that have no covariance with each other under the null hypothesis. Specifically, when the covariance is non-diagonal we can write:

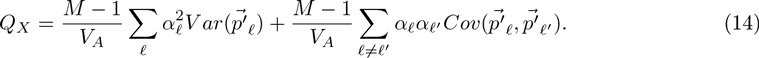

We refer to the first term in this sum as the standardized *F_ST_*-like component and the second term as the standardized LD-like component. We use this decomposition below to demonstrate for empirical examples that significant *Q_X_* statistics are due to unexpected covariance between like effect alleles, rather than unexpectedly large variance at individual loci.

### Identifying Outlier Populations

Having detected a putative signal of selection for a given trait, one may wish to identify individual regions and populations which contribute to the signal. Here we rely on our multivariate normal model of relatedness among populations, along with well understood methods for generating conditional multivariate normal distributions, in order to investigate specific hypotheses about individual populations or groups of populations.

Briefly, using standard results from multivariate normal theory, we can generate the expected joint conditional distribution of genetic values for an arbitrary set of populations given the observed genetic values in some other set of populations. These conditional distributions allow for a convenient way to ask whether the estimated genetic values observed in certain populations or groups of populations differ significantly from the values we would expect them to take given the values observed in related populations.

Specifically, we exclude a set of populations and then calculate the expected mean and variance of genetic values in these excluded populations given the remaining populations. Using this mean and variance we calculate a Z-score to describe how well fit the estimated genetic values of the excluded populations are by our model of drift, conditional on the values in the remaining populations. The details of our procedure are given in the Methods section, and we pursue implementations of this approach at two levels of population structure in our Empirical Applications.

### Datasets

We conducted power simulations and an empirical application of our methods based on the Human Genome Diversity Panel (HGDP) population genomic dataset (Li *et al.*, 2008), and a number of GWAS SNP sets. To ensure that we made the fullest possible use of the information in the HGDP data, we took advantage of a genome wide allele frequency dataset of ∼3 million SNPs imputed from the Phase II HapMap into the 52 populations of the HGDP. These SNPs were imputed as part of the HGDP phasing procedure in Pickrell *et al.* (2009); see our Methods section for a recap of the details. We applied our method to test for signals of selection in six human GWAS datasets identifying SNPs associated with height, skin pigmentation, body mass index (BMI), type 2 diabetes (T2D), Crohn’s Disease (CD) and Ulcerative Colitis (UC).

#### Choosing null SNPs

Various components of our procedure involve sampling random sets of SNPs from across the genome. While we control for biases in our test statistics introduced by population structure through our **F** matrix, we are also concerned that subtle ascertainment effects of the GWAS process could lead to biased test statistics, even under neutral conditions. We control for this possibility by sampling null SNPs so as to match the joint distribution of certain properties of the ascertained GWAS SNPs. Specifically, we chose our random SNPs to match the GWAS SNPs in each study in terms of their minor allele frequency (MAF) in the ascertainment population and the imputation status of the allele in our population genomic dataset. In addition, we were concerned that GWAS SNPs might be preferentially found close to genes and in low recombination regions, the latter due to better tagging, and as such may be subject to a high rate of drift due to background selection, leading to higher levels of differentiation at these sites (Charlesworth *et al.*, 1997). Therefore, in addition to MAF and imputation status, we also matched our random SNPs to an estimate of the background selection environment experienced by the GWAS SNPs, as measured by B value (McVicker *et al.*, 2009), which is a function of both the density of functional sites and recombination rate calibrated to match the reduction in genetic diversity due to background selection. We detail the specifics of the binning scheme for matching the discretized distributions of GWAS and random SNPs in the Methods.

### Power Simulations

To assess the power of our methods in comparison to other possible approaches, we conducted a series of power simulations. As statistical power is necessarily a function of the data, we sample 1000 null sets of SNPs for each phenotype according to the same matching procedure described above and in the Methods. We then gradually increase the effect of selection on these underlying loci, and measure the corresponding increase in power.

For each set of matched SNPs, we compared two of our statistics, (*r*^2^ and *Q_X_*) against their naive counterparts, which are not adjusted for population structure (*nr*^2^ and *Q_ST_*). For *Q_X_* and *Q_ST_*, we count a given simulation as having detected the signal of selection if it lies in the upper 5% tail of the null distribution, whereas for the environmental correlation statistics (*r*^2^ and *nr*^2^) we use a two-tailed 5% test. We also compared our tests to two single locus enrichment tests, the first where we counted the number of loci with individual SNP environmental correlations in the 5% tail using the single locus *r*^2^ (*slr*^2^) and in the second we counted the number of loci in the upper 5% tail of *F_ST_* (*slF_ST_*). We then considered these enrichment tests to have detected the signal of selection if the number of individual loci in the 5% tail was itself in the 5% tail using a binomial test. We do not include our alternative linear model statistics *β* and *ρ* in these plots for the sake of figure legibility, but they generally had very similar power to that of *r*^2^. While slightly more powerful versions of the single locus tests that account for sampling noise are available (e.g. WEIR and HILL, 2002; Coop *et al.*, 2010), note that our tests could be extended similarly as well, so the comparison is fair.

We base our power simulations on empirical data altered to have an increasing effect of selection along a latitudinal gradient. In order to mimic the effect of selection, we generate a new set of allele frequencies (*p_s,mℓ_*) by taking the original frequency (*p_mℓ_*) and adding a small shift according to

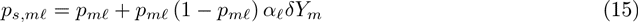

where *α*_ℓ_ is the effect size assigned assigned to locus *ℓ*, and *Y_m_* is the mean centered absolute latitude of the population. We used the 1000 simulations at *δ* = 0 to form null distribution for each of our test statistics, and from this established the 5% significance level. We then increment *δ* and give the power of each statistic as the fraction of simulations whose test statistic falls beyond this cutoff. While this approach to simulating selection is obviously naive to the way selection actually operates, it captures most of the important effects on the loci underlying a given trait. Namely, loci will have greater shifts if they experience rather extreme environments, have large effects on the phenotype, or are at intermediate frequencies. Because we add these shifts to allele frequencies sampled from real, putatively neutral loci, the effect of drift on their joint distribution is already present, and thus does not need to be simulated. Figure 1 shows simulations matched to the set of *p_ℓ_* and *α_ℓ_* of the Type 2 Diabetes dataset (see Figures S1–S5 for all other traits).

**Figure 1:**
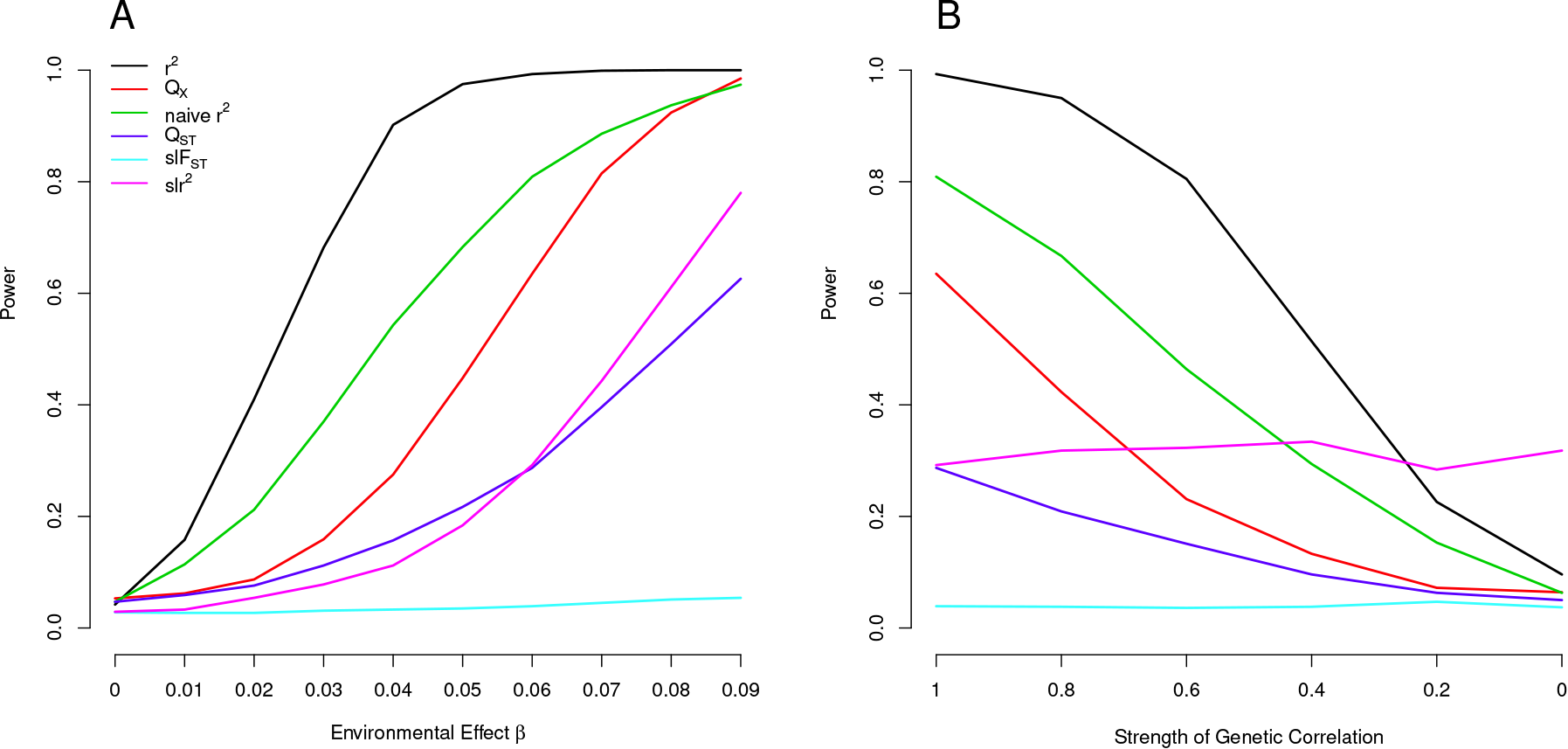
Power of our statistics as compared to alternative approaches (A) Across a range of effect sizes (*β*) of latitude, and (B) when we hold *β* constant at 0.06 and decrease *ϕ*, the genetic correlation between the trait of interest on the selected trait.

Our population structure adjusted statistics clearly outperform tests that do not account for structure, as well as single locus outlier based tests. Notably, tests based on an excess of *F_ST_* outliers or average *F_ST_* (not shown here) have essentially zero power for cases in which our tests have nearly 100% power. This nicely illustrates the fact that for highly polygenic traits, nearly all of the differentiation at the trait level arises as a consequence of across population linkage disequilibrium among the underlying loci, and not as a result of substantial differentiation at the loci themselves. While our environment-genetic value correlation tests outperform *Q_X_*, this is somewhat artificial as it assumes that we know the environmental variable responsible for our allele frequency shift. In reality we may at best hope to guess a correlate of the selection pressure, so *Q_X_* may outperform the environmental correlations.

In many cases, it may be that selection has not acted directly on the phenotype in question, but rather on one that is genetically correlated. Therefore, we are interested in the power of our tests to detect selection on traits genetically correlated with those for which we have GWAS data. The question of how to properly simulate this relationship is largely an empirical one, as the correlation structure underlying any two traits will depend on their shared genetic architecture. We therefore chose a simple and general approach to capture a flavor of this situation. We simulate the effect of selection as above (15), but using effect sizes 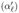 for an unobserved phenotype which are only partially correlated with our observed effect sizes. For simplicity we assume that *α_ℓ_* and 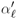 have a bivariate normal distribution around zero with equal variance and correlation parameter *ϕ*, and then simulate 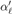 from its conditional distribution given *α_ℓ_*(i.e. 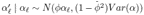). For each SNP *ℓ* in (15) we replaced *α* by its effect 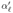 on the unobserved phenotype. Here *ϕ* can be thought of as the genetic correlation between our phenotypes if this simple multivariate form held true for all of the loci contributing to our two traits.

In Figure 1 we hold the value of *β* constant at 0.06 and vary this genetic correlation *φ* from one down to zero. Predictably, our GWAS prediction-based statistics lose power as the two traits become more and more genetically independent, but do retain reasonable power out to quite low genetic correlations. In contrast, counting the number of SNPs that are significantly correlated with a given environmental variable remains equally powerful across all effect sizes, as these loci are still altered from neutral expectations along the environmental gradient, and the loci’s effect sizes and directionality do not enter into our calculation.

In some ways this may be considered a desirable property of the environmental outliers enrichment approach, it can still detect the effect of selection on these loci, even if the genetic effect sizes of the two traits are uncorrelated. On the other hand, this is also problematic, as it implies that without detailed knowledge of the various phenotypic associations at each locus, such tests may often be detecting selection on pleiotropically related phenotypes. Our approaches, however, are more suited to determining whether the genetic basis of a trait of interest has been affected by selection on a trait that is at least somewhat genetically correlated.

### Empirical Applications

We estimated genetic values for each of six traits from the subset of GWAS SNPs that were present in the HGDP dataset, as described above. We discuss the analysis of each dataset in detail below, and address general points first. For each dataset, we constructed the covariance matrix from a sample of approximately 20, 000 appropriately matched SNPs, and the null distributions of our test statistics from a sample of 10, 000 sets of null genetic values, which were also constructed according to a similar matching procedure (as described in the Methods).

To test for an effect of major climate variables on the genetic basis of a given trait, we used the set of six climate variables (measured in both summer and winter seasons) considered by Hancock *et al.* (2008) in their analysis of adaptation to climate at the level of individual SNPs. We followed their general procedure by running principal components (PC) analysis for both seasons on a matrix containing the six climate variables, as well as latitude and longitude (following Hancock *et al.*’s (2008) rationale that these two geographic variables may capture certain elements of the long term climatic environment experienced by human populations). The percent of the variance explained by these PCs and their weighting (eigenvectors) of the different environmental variables are given in Table 1.

**Table 1:** The contribution of each geo-climatic variable to each of our four principal components, scaled such that the absolute value of the entries in each column sum to one (up to rounding error). We also show for each principal component the percent of the total variance across all eight variables that is explained by the PC.

| Geo-Climatic Variable | SUMPC1 | SUMPC2 | WINPC1 | WINPC2 |
| --- | --- | --- | --- | --- |
| Latitude | -0.16 | -0.10 | -0.17 | -0.01 |
| Longitude | 0.02 | 0.12 | 0.04 | 0.05 |
| Maximum Temp | 0.24 | -0.08 | 0.17 | -0.03 |
| Minimum Temp | 0.24 | 0.07 | 0.17 | 0.08 |
| Mean Temp | 0.25 | -0.03 | 0.17 | 0.03 |
| Precipitation Rate | -0.01 | 0.16 | 0.07 | 0.32 |
| Relative Humidity | -0.06 | 0.21 | -0.06 | 0.34 |
| Short Wave Radiation Flux | -0.03 | -0.22 | 0.15 | -0.13 |
| Percent of Variance Explained | 38% | 35% | 58% | 20% |

We also applied our *Q_X_* test to identify traits whose underlying loci showed consistent patterns of unusual differentiation across populations. In Figure 2 we show for each GWAS set the observed value of *Q_X_* and the empirical null distribution of *Q_X_* using SNPs matched to the GWAS loci as described above. We also plot the expected null distribution of the *Q_X_* statistic 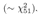 The expected null distribution closely matches the empirical distribution, suggesting that our multivariate normal framework provides a good null model for the data (although we will use the empirical null distribution to obtain measures of statistical significance).

**Figure 2:**
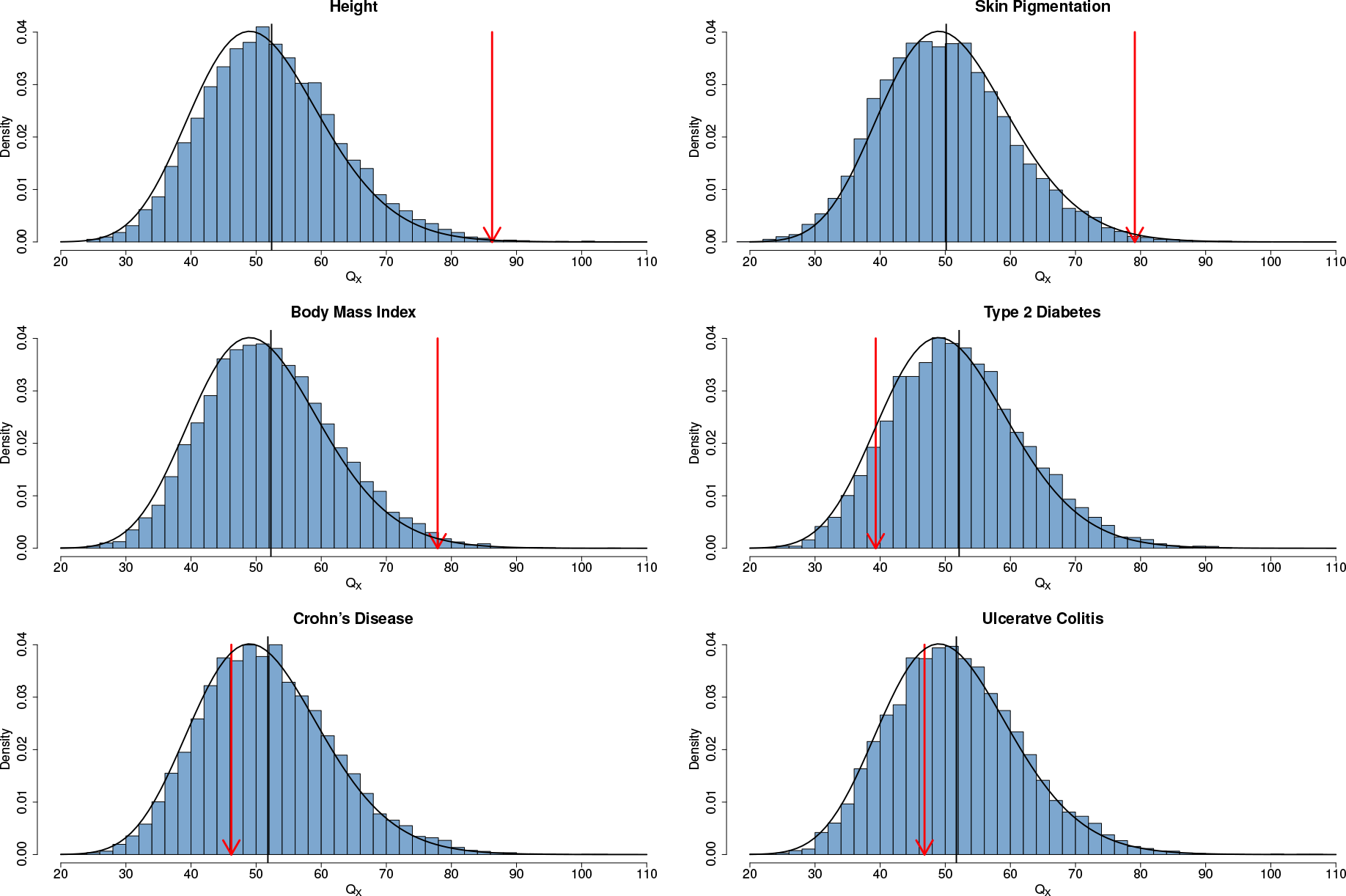
Histogram of the empirical null distribution of *Q_X_* for each trait, obtained from genome-wide resampling of well matched SNPs. The mean of each distribution is marked with a vertical black bar and the observed value is marked by a red arrow. The expected 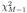 density is shown as a black curve.

For each GWAS SNP set we can also separate our *Q_X_* statistic into its two components (an *F_ST_*-like component and an LD-like term), as described in (14). In Figure 3 we plot the null distributions of these two components for the height dataset as histograms, with the observed value marked by red arrows. As with all of the significant *Q_X_* statistics we have examined, the signal of unusual differentiation comes largely from the standardized LD-like component (see Figures S6–S10 for all other traits at the global level). This highlights the fact that the information about departures from neutrality comes largely from a subtle signal of covariance across loci of like effect alleles.

**Figure 3:**
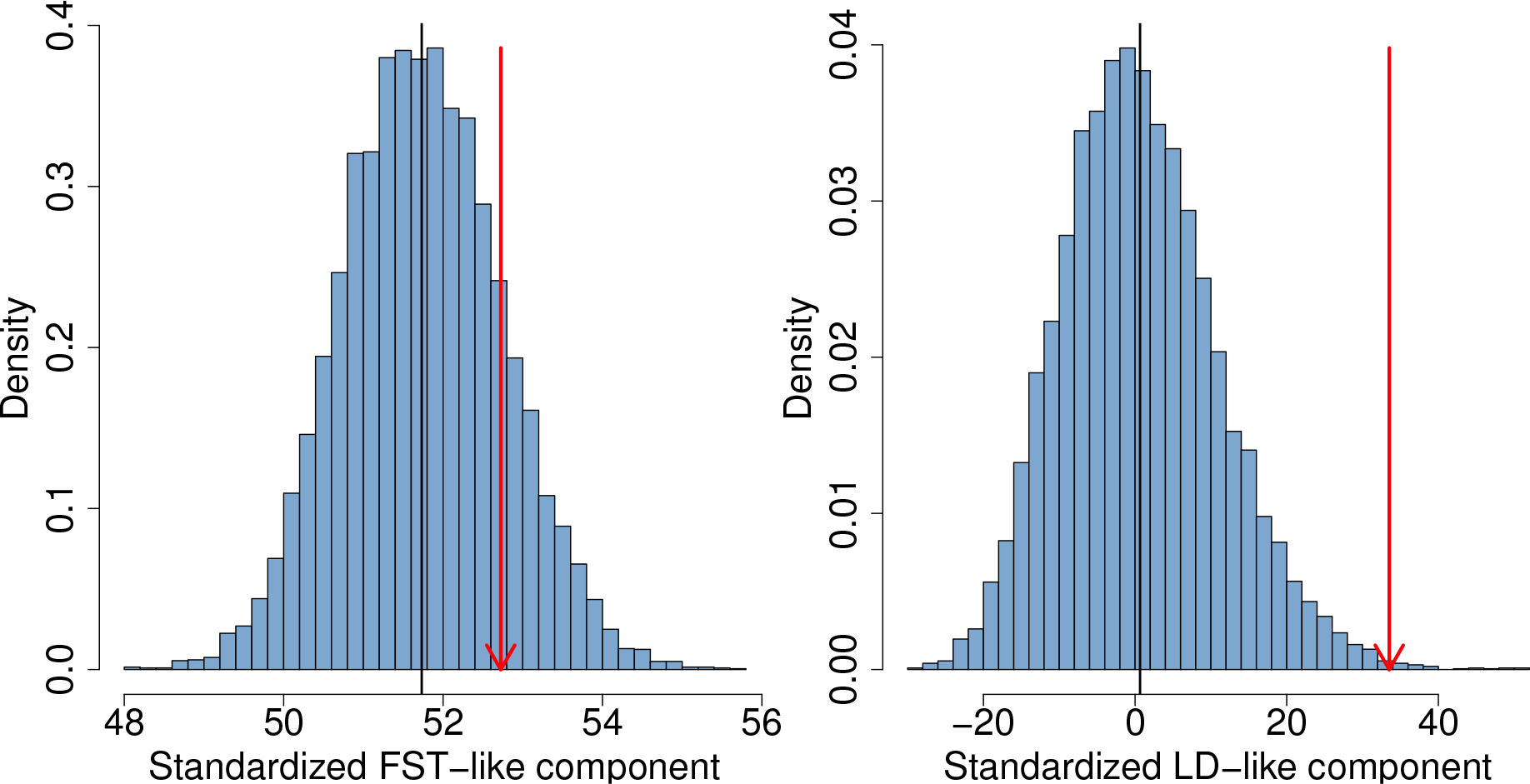
The two components of *Q_X_* for the height dataset, as described by the left and right terms in (14). The null distribution of each statistic is shown as a histogram. The mean value is shown as a black bar, and the observed value as a red arrow.

As discussed above and in the Methods, we can use our multivariate normal framework to identify specific regions or populations that are poorly fit by the neutral model when predicted from other populations. We do this for each of our GWAS SNP sets at two different levels of population structure: seven broad geographic regions (as identified by Rosenberg *et al.*, 2002) as well as individual populations. We first calculate the average of the estimated genetic values within one of the seven broad regions. We then calculate the expected joint conditional distribution for the populations in that region, given the estimated genetic values observed in each of the other six regions. This results in a vector of expectations for each population, and a conditional covariance matrix relating them, which we can convert into to the expected conditional mean and variance of our regional average. We use this expected conditional mean and variance and the observed regional average to calculate a Z score as a measure of how differentiated that region is from the rest of the world. In Figures 5 and 6, we present the conditional expectation for each region as a light grey dashed line, and the observed mean as a solid bar, with darkness and thickness proportional to Z score.

We also calculate the conditional expectation and variance for each population in turn, given the values observed for all other populations in the dataset, allowing us to calculate population specific Z scores. Whereas our regional level Z scores are designed to detect selection events occurring over broader geographical regions (and potentially deeper time scales), our populations specific Z scores are designed to identify individual populations that are not well predicted by the model, and thus may have experienced more recent selection. In Figures 5 and 6, we present each population’s estimated genetic value as a colored circle, with its size proportional to its population specific Z score. Full characterizations of these results are available in Tables S3–S14.

#### Height

We first analyzed the set of 180 height associated loci identified by Lango Allen *et al.* (2010), which explain approximately 14% of the additive genetic variance for height (Zaitlen *et al.*, 2013). This dataset is an ideal first test for our methods because it contains the largest number of associations identified for a single phenotype to date, maximizing our power gain over single locus methods (Figure S1). In addition, Turchin *et al.* (2012) have already identified a signal of pervasive weak selection at these same loci in European populations, and thus we should expect our methods to replicate this observation.

Of the 180 loci identified by Lango Allen *et al.* (2010), 161 were present in our HGDP dataset. We used these 161 loci in conjunction with the allele frequency data from the HGDP dataset to estimate genetic values for height in the 52 HGDP populations. Although these genetic values are correlated with the observed heights of these populations, as expected they are far from perfect predictions (see Figure S11 and Table S1, which compares our estimated genetic values to observed sex-average heights for the subset of HGDP populations with a close proxy in the dataset of Gustafsson and Lindenfors, 2009). We find a signal of excessive correlation with winter PC2 (Figure 4 and Table 2), but find no effect of any other climatic variables. Our *Q_X_* test also strongly rejects the neutral hypothesis, suggesting that our estimated genetic values are overly dispersed compared to the null model of neutral genetic drift and shared population history (Figure 2 and Table 2),

**Figure 4:**
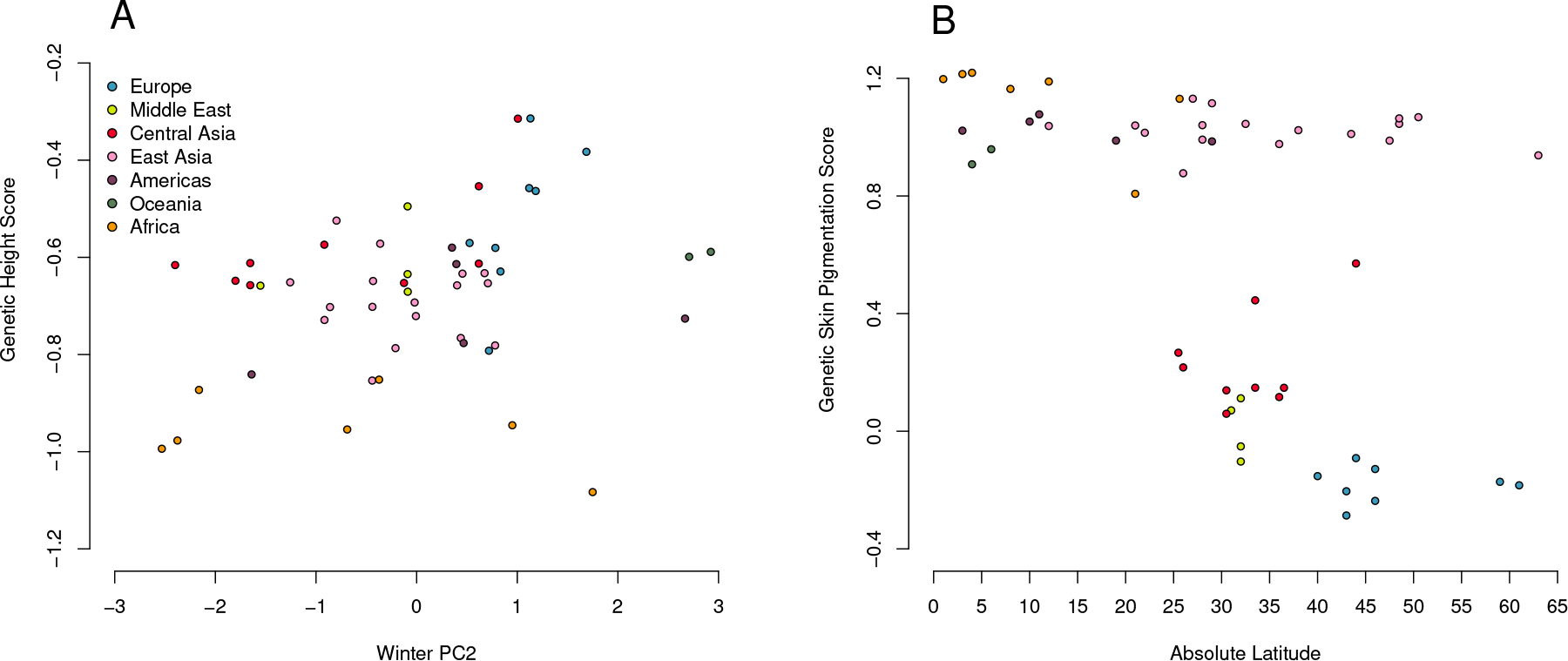
Estimated genetic height (A) and skin pigmentation score (B) plotted against winter PC2 and absolute latitude respectively. Both correlations are significant against the genome wide background after controlling for population structure (Table 2).

**Table 2:**
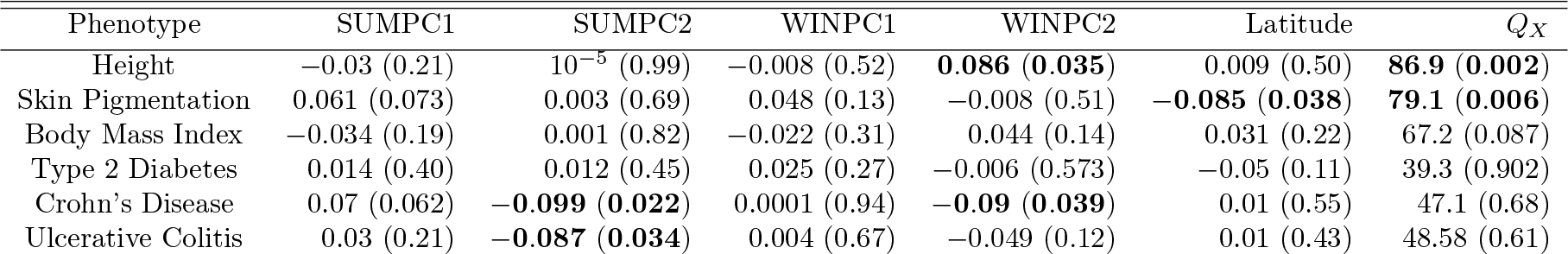
Climate Correlations and *Q_X_* statistics for all six phenotypes in the global analysis. We report *sign*(*β*)*r*^2^, for the correlation statistics, such that they have an interpretation as the fraction of variance explained by the environmental variable, after removing that which is explained by the relatedness structure, with sign indicating the direction of the correlation. P-values are two–tailed for *r*^2^ and upper tail for *Q_X_*. Values for *β* and *ρ* are reported in Tables S15 and S16.

We followed up on this signal of differentiation by conducting regional level analyses, which indicate that the worldwide signal of excess variance arises from both inter– and intra–regional processes. We find that the mean genetic height in Africa is lower than expected given the values observed in the rest of the world (*Z* = −2.0, *p* = 0.045; see also Figure 5 and Table S3). We also find evidence for differential selection within Europe, as suggested by Turchin *et al.* (2012), and to a somewhat lesser extent in the Middle East and Central Asia (Figure 5 and Table 3).

**Figure 5:**
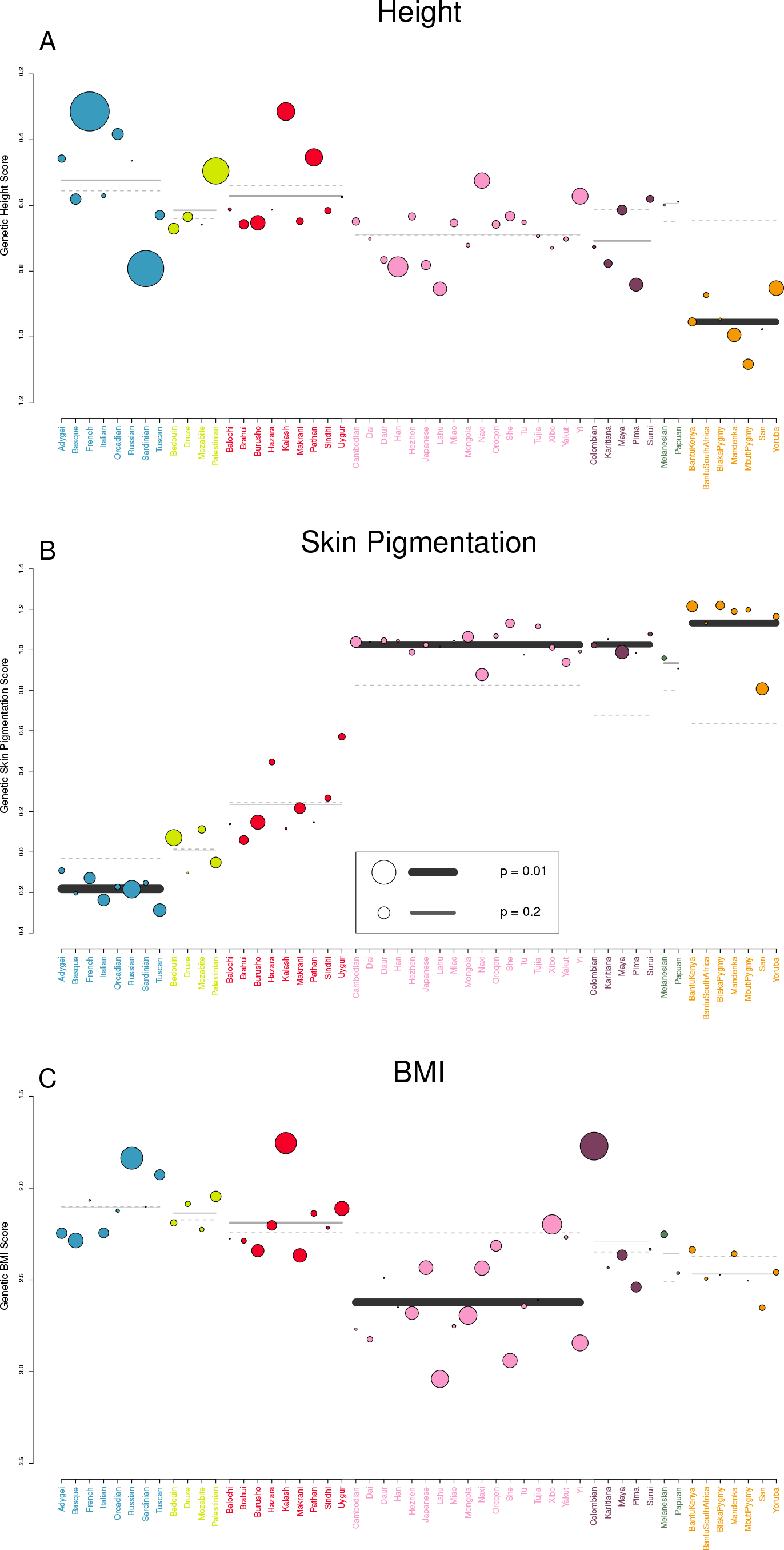
Visual representation of outlier analysis at the regional and individual population level for (A) Height, (B) Skin Pigmentation, and (C) Body Mass Index. For each geographic region we plot the expectation of the regional average, given the observed values in the rest of the dataset as a grey dashed line. The true regional average is plotted as a solid bar, with darkness and thickness proportional to the regional Z score. For each population we plot the observed value as a colored circle, with circle size proportional to the population specific Z score. For example, in (A), one can see that estimated genetic height is systematically lower than expected across Africa. Similarly, estimated genetic height is significantly higher (lower) in the French (Sardinian) population than expected, given the values observed for all other populations in the dataset.

**Table 3:**
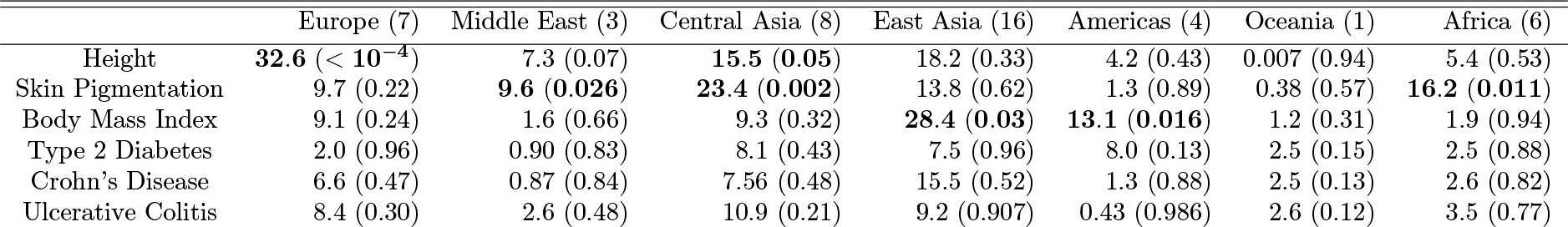
*Q_X_* statistics and their empirical p-values for each of our six traits in each of the seven geographic regions delimited by Rosenberg *et al.* (2002). The theoretical expected value of the statistic under neutrality for each region is equal to *M* − 1, where *M* is the number of populations in the region. We report the value of *M* − 1 next to each region for reference.

|  | Europe (7) | Middle East (3) | Central Asia (8) | East Asia (16) | Americas (4) | Oceania (1) | Africa (6) |
| --- | --- | --- | --- | --- | --- | --- | --- |
| Height | **32.6** ($< 10^{-4}$) | 7.3 (0.07) | **15.5** (**0.05**) | 18.2 (0.33) | 4.2 (0.43) | 0.007 (0.94) | 5.4 (0.53) |
| Skin Pigmentation | 9.7 (0.22) | **9.6** (**0.026**) | **23.4** (**0.002**) | 13.8 (0.62) | 1.3 (0.89) | 0.38 (0.57) | **16.2** (**0.011**) |
| Body Mass Index | 9.1 (0.24) | 1.6 (0.66) | 9.3 (0.32) | **28.4** (**0.03**) | **13.1** (**0.016**) | 1.2 (0.31) | 1.9 (0.94) |
| Type 2 Diabetes | 2.0 (0.96) | 0.90 (0.83) | 8.1 (0.43) | 7.5 (0.96) | 8.0 (0.13) | 2.5 (0.15) | 2.5 (0.88) |
| Crohn's Disease | 6.6 (0.47) | 0.87 (0.84) | 7.56 (0.48) | 15.5 (0.52) | 1.3 (0.88) | 2.5 (0.13) | 2.6 (0.82) |
| Ulcerative Colitis | 8.4 (0.30) | 2.6 (0.48) | 10.9 (0.21) | 9.2 (0.907) | 0.43 (0.986) | 2.6 (0.12) | 3.5 (0.77) |

#### Skin Pigmentation

We next analyzed data from a recent GWAS for skin pigmentation in an African-European admixed population of Cape Verdeans (Beleza *et al.*, 2013), which identified four loci of major effect that explain approximately 35% of the variance in skin pigmentation in that population after controlling for admixture proportion. Beleza *et al.* (2013) report effect sizes in units of modified melanin (MM) index, which is calculated as 100 × log(1*/*%melanin reflectance at 650 nM), i.e. a higher MM index corresponds to darker skin, and a lower value to lighter skin.

We used these four loci to calculate a genetic skin pigmentation score in each of the HGDP populations. As expected, we identified a strong signal of excess variance among populations, as well as a strong correlation with latitude (Figure 4 and Table 2). Note, however, that this signal was driven entirely by the fact that populations of western Eurasian descent have a lower genetic skin pigmentation score than populations of African descent. Using only the markers from Beleza *et al.* (2013), light skinned populations in East Asian and the Americas have a genetic skin pigmentation score that is almost as high as that of most African populations, an effect that is clearly visible when we plot the measured skin pigmentation and skin reflectance of HGDP populations (Jablonski and Chaplin, 2000; Lao *et al.*, 2007) against their genetic values (see Figures S12 and S13).

The correlation with latitude is thus weaker than one might expect, given the known phenotypic distribution of skin pigmentation in human populations (Jablonski and Chaplin, 2000, 2010). To illustrate this point further, we re-ran the analysis on a subsample of the HGDP consisting of populations from Europe, the Middle East, Central Asia, and Africa. In this subsample, the correlation with latitude, and signal of excess variance, was notably stronger (*r*^2^ = 0.2, *p* = 0.019; *Q_X_* = 60.1, *p* = 8 × 10^−4^).

This poor fit to observed skin pigmentation is due to the fact that we have failed to capture all of the loci that contribute to variation in skin pigmentation across the range of populations sampled, likely due to the partial convergent evolution of light skin pigmentation in Western and Eastern Eurasian populations (Norton *et al.*, 2006). Including other loci putatively involved in skin pigmentation (Miller *et al.*, 2007; Edwards *et al.*, 2010) decreases the estimated genetic pigmentation score of the other Eurasian populations (Figures S12 and S13), but we do not include these in our main analyses as they differ in ascertainment (and the role of KITLG in pigmentation variation has been contested by Beleza *et al.*, 2013).

Within Africa, the San population has a decidedly lower genetic skin pigmentation score than any other HGDP African population. This is in accordance with the observation by Jablonski and Chaplin (2000) that the San are more lightly pigmented than other African populations represented by the HGDP and the observation that other putative light skin pigmentation alleles have higher frequency in the San than other African populations (Norton *et al.*, 2006). A regional analysis of the six African populations alone identified a marginally significant correlation with latitude (*r*^2^ = 0.62, *p* = 0.0612), and a signal of excess variance among populations (*Q_X_* = 16.19, *p* = 0.01), suggesting a possible role for selection in shaping pigmentation variation within Africa.

#### Body Mass Index

We next investigate two traits related to metabolic phenotypes (BMI and Type 2 diabetes), as there is a long history of adaptive hypotheses put forward to explain phenotypic variation among populations, with little conclusive evidence emerging thus far. We first focus on the set of 32 BMI associated SNPs identified by Speliotes *et al.* (2010) in their Table 1, which explain approximately 2-4% of the genetic variance for BMI. Of these 32 associated SNPs, 28 were present in the HGDP dataset, which we used to calculate a genetic BMI score for each HGDP population. We identified no significant signal of selection at the global level (Table 2).

Our regional level analysis indicated that the mean genetic BMI score is significantly lower that expected in East Asia (*Z* = −2.48, *p* = 0.01; see also Figure 5 and Table S7), while marginal *Q_X_* statistics identify excess intraregional variation within East Asia and the Americas (Table 3).

#### Type 2 Diabetes

We next investigated the 65 loci reported by Morris *et al.* (2012) as associated with T2D in their supplementary Table 2. Of these 65 SNPs, 61 were present in the HGDP dataset. We used effect sizes from stage 1 meta-analysis, and where Morris *et al.* (2012) reported a range of allele frequencies (due to differing sample frequencies among cohorts), we simply used the average. Where multiple SNPs were reported per locus we used the lead SNP from the combined meta-analysis. Also note that Morris *et al.* (2012) report effects in terms odds ratios (OR), which can be converted into additive effects by taking the logarithm (the same is true of the IBD data from Jostins *et al.*, 2012, analyzed below).

The distribution of genetic T2D risk scores showed no significant correlations with any of the five eco-geographic axes we tested, and was in fact fairly underdispersed worldwide relative to the null expectation due to population structure (Table 2).

Our regional level analysis revealed that while T2D genetic risk is well explained by drift in Africa, Central and Eastern Asia, Oceania, and the Americas, European populations have far lower T2D genetic risk than expected (*Z* = −2.79, *p* = 0.005) and Middle Eastern populations a higher genetic risk than expected (*Z* = 2.37, *p* = 0.018). It’s not necessarily clear, however, that these observations should be interpreted as evidence for selection either in Europe or the Middle East. While the dichotomous regional labels “Europe” vs. “Middle East” and genetic T2D risk explains the majority of the variance not accounted for by population structure (*r*^2^ = 0.77, *p* = 5 × 10^−4^), this is essentially the same signal detected by the regional Z scores, and our *Q_X_* statistic finds no signal of excess variance (*Q_X_* = 10.9, *p* = 0.48) among the twelve HGDP populations in these two regions. Expanding to the next most closely related region, we tested for a signal of excess differentiation between Central Asia and either Europe or Middle East, but find no convincing signal in either case (*r*^2^ = 0.13, *p* = 0.21; *Q_X_* = 12.0, *p* = 0.75 and *r*^2^ = 0.15, *p* = 0.19; *Q_X_* = 9.8, *p* = 0.63 respectively).

To the extent that our results are consistent with an impact of selection on the genetic basis of T2D risk, they appear to be consistent only with a scenario in which selection has pushed the frequency of alleles that increase T2D risk up in Middle Eastern populations and down in European populations. This is an extremely subtle signal, which arises only after deep probing of the data, and as such we are skeptical that our results represent a meaningful signal of selection.

#### IBD

Finally, we analyzed the set of associations reported by Jostins *et al.* (2012) for Crohn’s Disease (CD) and Ulcerative Colitis (UC). Because CD and UC are closely connected phenotypes that share much of their genetic etiology, Jostins *et al.* (2012) used a likelihood ratio test of four different models (CD only, UC only, both CD and UC with equal effects on each, both CD and UC with independent effects) to distinguish which SNPs where associated with either or both phenotypes, and to assign effect sizes to SNPs (see their supplementary methods section 1d). We take these classifications at face value, resulting in two partially overlapping lists of 140 and 135 SNPs associated with CD and UC, of which 95 and 89 respectively were present in our HGDP dataset.

We used these sets of SNPs to calculate genetic risk scores for CD and UC across the 52 HGDP populations. Both CD and UC showed strong negative correlations with summer PC2 (Figure 7), while CD also showed a significant correlation with winter PC1, and a marginally significant correlation with summer PC1 (Table 2).

**Figure 6:**
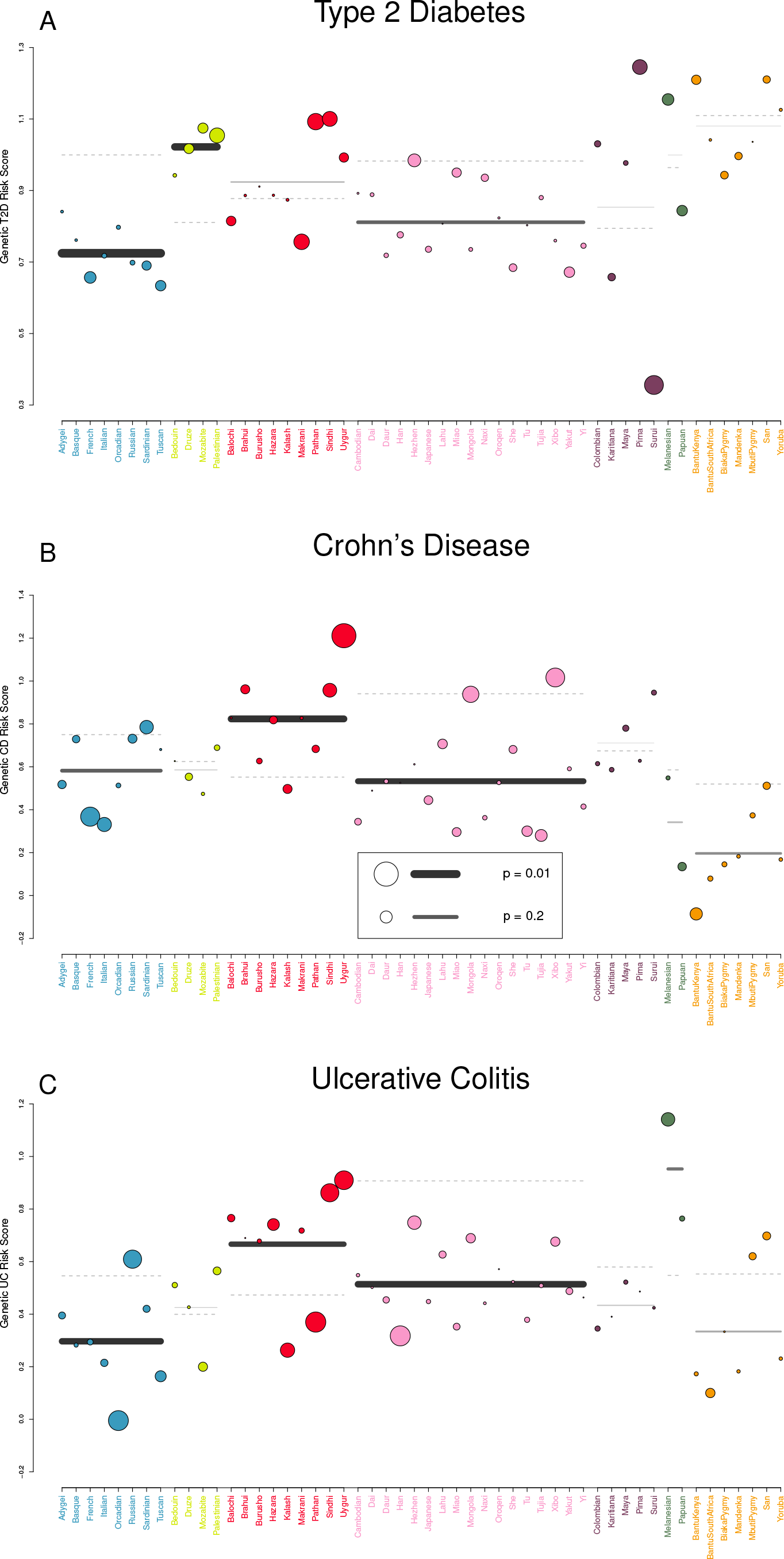
Visual representation of outlier analysis at the regional and individual population level for (A) Type 2 Diabetes, (B) Crohn’s Disease, and (C) Ulcerative Colitis. See Figure 5 for explanation.

**Figure 7:**
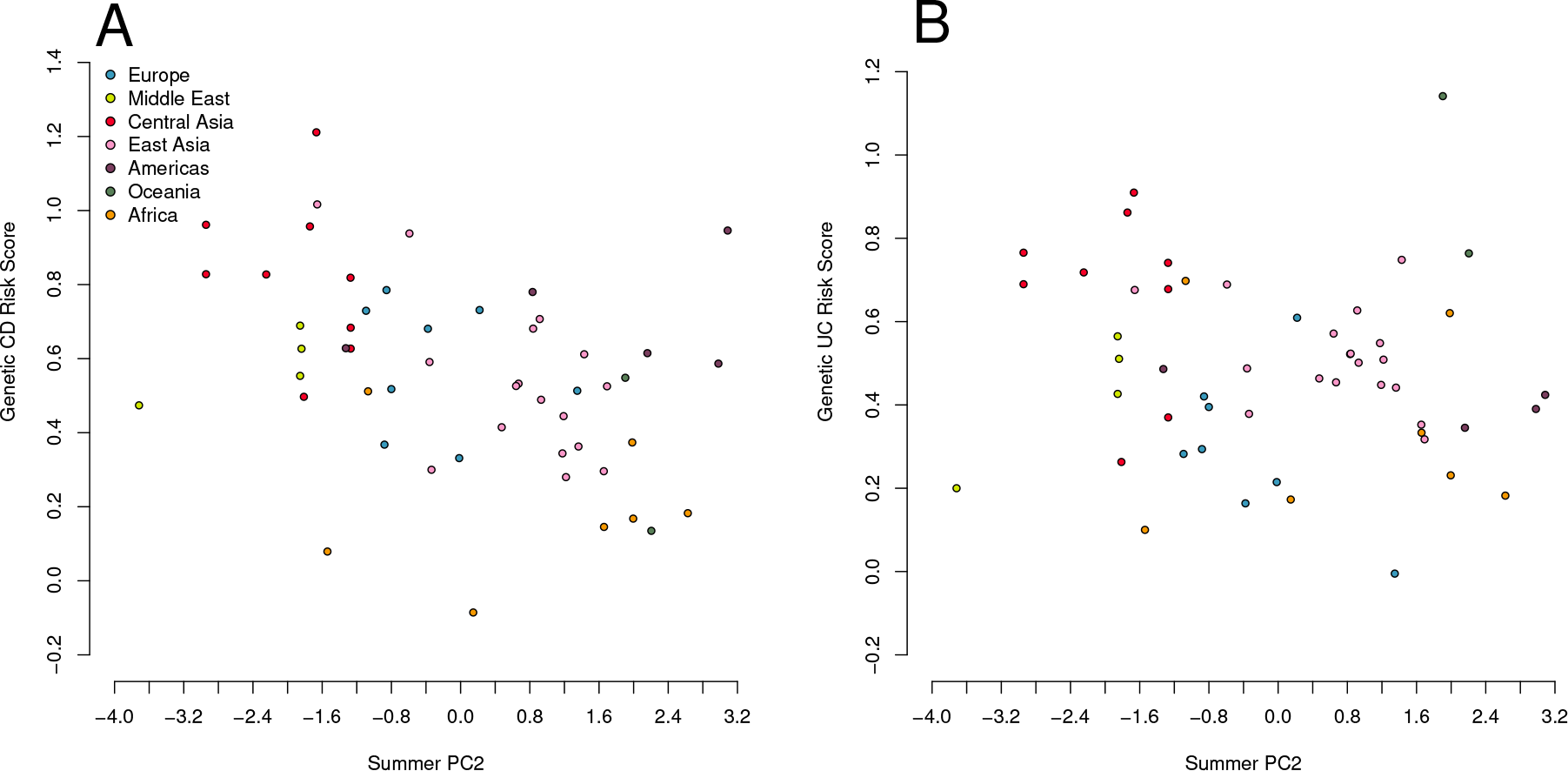
Estaimted genetic risk score for Crohn’s Disease (A) and Ulcerative Colitis (B) risk plotted against summer PC2. Both correlations are significant against the genome wide background after controlling for population structure (Table 2). Since a large proportion of SNPs underlying these traits are shared, we note that these results are not independent.

We did not observe any significant *Q_X_* statistics for either trait, either at the global or the regional level, suggesting that our environmental correlation signals most likely arise from subtle differences between regions, as opposed to divergence among closely related populations. Indeed, we find moderate signals of regional level divergence in Europe (UC: *Z* = −2.08, *p* = 0.04), Central Asia (CD: *Z* = 2.21, *p* = 0.03), and East Asia (CD: *Z* = −1.90, *p* = 0.06 and UC: *Z* = −2.12, *p* = 0.03; see also Figure 6 and Tables S13 and S14).

## 3 Discussion

In this paper we have developed a powerful framework for identifying the influence of local adaptation on the genetic loci underlying variation in polygenic phenotypes. Below we discuss two major issues related to the application of such methods, namely the effect of the GWAS ascertainment scheme on our inference, and the interpretation of positive results.

### Ascertainment and Population Structure

Among the most significant potential pitfalls of our analysis (and the most likely cause of a false positive) is the fact that the loci used to test for the effect of selection on a given phenotype have been obtained through a GWAS ascertainment procedure, which can introduce false signals of selection if potential confounds are not properly controlled. We condition on simple features of the ascertainment process via our allele matching procedure, but deeper issues may arise from artifactual associations that result from the effects of population structure in the GWAS ascertainment panel. Given the importance of addressing this issue to the broader GWAS community, a range of well developed methods exist for doing GWAS in structured populations, and we refer the reader to the existing literature for a full discussion (Freedman *et al.*, 2004; Campbell *et al.*, 2005; Price *et al.*, 2006; Kang *et al.*, 2008; Price *et al.*, 2010; Diao and Chen, 2012; Liu *et al.*, 2013). Here, we focus on two related issues. First, the propensity of population structure in the GWAS ascertainment panel to generate false positives in our selection analysis, and second, the difficulties introduced by the sophisticated statistical approaches employed to deal with this issue when GWAS are done in strongly structured populations.

The problem of population structure arises generally when there is a correlation in the ascertainment panel between phenotype and ancestry (due to either genetic or environmental effects), such that SNPs that are ancestry informative will appear to be associated with the trait, even when no causal relationship exists (e.g. Campbell *et al.*, 2005). To make matters worse, multiple false positive associations will often line up with same axis of population structure. If the populations being tested with our methods lie at least partially along the same axis of structure present in the GWAS ascertainment panel, then the ascertainment process will serve to generate the very signal of positive covariance among like effect alleles that our methods rely on to detect the signal of selection.

The primary takeaway from this observation is that the more diverse the array of individuals sampled for a given GWAS are with respect to ancestry, the greater the possibility that failing to control for population structure will generate false associations and hence false positives for our method.

What bearing do these complications have on our empirical results? The GWAS datasets we used can be divided into those conducted within populations of European descent and the skin pigmentation dataset (which used an admixed population). We will first discuss our analysis of the former.

The European GWAS loci we used were found in relatively homogeneous populations, in studies with rigorous standards for replication and control for population structure. Therefore, we are reasonably confident that these loci are true positives. Couple this with the fact that they were ascertained in (European) populations that are fairly homogenous relative to the global scale of our analyses, it is unlikely that population structure in the ascertainment panels is driving our positive signals. One might worry that we could still generate false signals by including European populations in our analysis, however many of the signals we see are driven by patterns outside of Europe (where the influence of structure within Europe should be much lessened). For height, where we do see a strong signal from within Europe, we use a set of loci that Turchin *et al.* (2012) have independently verified using a family based design that is immune to the effects of population structure.

We further note that for a number of our GWAS datasets, studies of non-European populations have replicated many of these loci (Cho *et al.*, 2009, 2011; Voight *et al.*, 2010; Kooner *et al.*, 2011; N’Diaye *et al.*, 2011; Carty *et al.*, 2012; **?**). More broadly, a recent study by Marigorta and Navarro (2013) found that replication rates for disease GWAS in European populations were quite high in East Asian populations, consistent with simple power considerations, an unlikely result if associations are driven primarily by population structure. This suggests that, at least for GWAS done in relatively homogenous human populations, structure is unlikely to be a major confounding factor.

The issue of population structure may be more profound for our style of approach when GWAS are conducted using individuals from more strongly structured populations. In some cases it is desirable to conduct GWAS in such populations as locally adaptive alleles will be present at intermediate frequencies in these broader samples. A range of methods have been developed to adjust for population structure in these setting (Kang *et al.*, 2010; Zhou and Stephens, 2012; Liu *et al.*, 2013). While generally effective in their goal, these methods present their own issues for our selection analysis. Consider the extreme case, such as that of Atwell *et al.* (2010), who carried out a GWAS in *Arabidopsis thaliana* for 107 phenotypes across an array of 183 inbred lines of diverse geographical and ecological origin. Atwell *et al.* (2010) used the genome-wide mixed model program EMMA (Kang *et al.*, 2008, 2010; Zhou and Stephens, 2012) to control for the complex structure present in their ascertainment panel. This practice helps ensure that many of the identified associations are likely to be real, but also means that the loci found are likely to have unusual frequencies patterns across the species range. This follows from the fact that the loci identified as associated with the trait must stand out as being correlated with the trait in a way not predicted by the individual kinship matrix (as used by EMMA and other mixed model approaches). Our approach is predicated on the fact that we can use genome-wide patterns of kinship to adjust for population structure, but this correction is exactly the null model that loci significantly associated with phenotypes by mixed models have overcome. For this reason, both the theoretical *χ*^2^ distribution of the *Q_X_* statistic, as well as the empirical null distributions we construct from resampling, are likely to be inappropriate.

The Cape Verde skin pigmentation data we used could qualify as this second type of study. The Cape Verde population is an admixed population of African/European descent, and has substantial inter-individual variation in admixture proportion. Due to its admixed nature, the population segregates alleles which would not be at intermediate frequency in either parental population, and the four loci mapped in the study explain approximately 35% of the variance in skin pigmentation within the population (Beleza *et al.*, 2013).

Despite the considerable population structure, the fact that interbreeding continues to mix genotypes in this population means that much of the LD due to the African/European population structure has been broken up (and the remaining LD is well predicted by an individual’s genome-wide admixture coefficient). Population structure seems to have been well controlled for in this study, and a number of the loci have been replicated in independent admixed populations. While it is thus unlikely that the four loci we use are false associations, they could in principle suffer from the structured ascertainment issues described above, so it is unclear that the null distributions we use are strictly appropriate. That said, provided that Beleza *et al.* (2013) have indeed appropriately controlled for population structure, under neutrality there would be no reason to expect that the correlation among the loci should be strongly positive with respect to the sign of their effect on the phenotype, and thus the pattern observed is at least consistent with a history of selection, especially in light of the multiple alternative lines of evidence for adaptation on the basis of skin pigmentation (Jablonski and Chaplin, 2000; Sabeti *et al.*, 2007; Lao *et al.*, 2007; Williamson *et al.*, 2007; Sturm, 2009; Jablonski and Chaplin, 2010).

Further work is needed to determine how best to modify the tests proposed herein to deal with GWAS performed in structured populations.

### Complications of Intepretation

Our understanding of the genetic basis of most complex traits remains very incomplete, and as such the results of these analyses must be interpreted with care. That said, because our methods are based simply on the rejection of a neutral null model, an incomplete knowledge of the genetic basis of a given trait should only lead to a loss of statistical power, and not to a high false positive rate.

For all traits analyzed here except for skin pigmentation, the within population variance for genetic value is considerably larger than the variance between populations. This suggests that much of what we find is relatively subtle adaptation even on the level of the phenotype, and emphasizes the fact that for most genetic and phenotypic variation in humans, the majority of the variance is within populations rather than between populations (see Figures S14–S19). The influence of the environment likely plays a stronger role in the differences between populations for true phenotypes than the subtle differences we find here (as demonstrated by the rapid change in type 2 diabetes incidence with changing diet, e.g. Franco *et al.*, 2013). That said, an understanding of how adaptation has shaped the genetic basis of a wide variety of phenotypes is clearly of interest, even if environmental differences dominate as the cause of present day population differences, as it informs our understanding of the biology and evolutionary history of these traits.

The larger conceptual issues relate to the interpretation of our positive findings, which we detail below. A number of these issues are inherent to the conceptual interpretation of evidence for local adaptation (Kawecki and Ebert, 2004).

#### Environmental Correlations

Significant correlation with environmental variables must be interpreted with caution. It is unlikely that the environmental variables included in our PC set will include the agent of selection and rather may simply be a correlate of the actual selection pressure. Further, given the large geographic and temporal scales considered here, it seems unlikely that many of our positive findings are the result of any single selection pressure, but rather may result from a combination of many different pressures, with perhaps different pressures acting in different geographic regions.

Consider also that we investigate only a limited set of phenotypes, which will each covary genetically with many other phenotypes. As such, any evidence for selection acting on the genetic basis of one of our phenotypes may represent selection on some correlated phenotype(s) not included in our study.

As an example, consider the fact that both UC and CD show a negative correlation with summer PC2, which include humidity and rainfall as primary components. However, it seems very unlikely that UC or CD have been the direct target of selection. It has been proposed that UC and CD might be understood as pleiotropic side effects of pathogen response (Hansen *et al.*, 2010; Vatn, 2012; Cagliani *et al.*, 2013), in which case the signal we see might correspond to local adaptation to variation in pathogen load/diversity (with humidity and/or rainfall as a possible predictor of these differences). Our results are likely to be compatible with many different hypotheses, however, and thus we consider this only one possible explanation.

#### Gene-by-Environment Interactions

Gene by environment interactions may also complicate considerably the interpretation of a positive result. For example, we find evidence for adaptation in height, particularly in western Eurasia (consistent with the results of Turchin *et al.*, 2012), along with a worldwide correlation with winter PC2. However, our effect estimates are specific to a modern, northern European environment (although we note that the loci thus far identified in non European populations tend to overlap those discovered in Europe: Cho *et al.*, 2009; N’Diaye *et al.*, 2011; Carty *et al.*, 2012). It is possible that these alleles have experienced selection due to different effects in different environments, and so evidence for differentiation at the genetic level may not align with the observed phenotypic distribution. In an extreme case, different sets of alleles could be locally selected to maintain a constant phenotype across populations due to gene-by-environment interactions. Such a scenario could lead to a signal of local adaptation on a genetic level but no change in the phenotype across populations, a phenomenon known as countergradient variation (Conover and Schultz, 1995).

It is also the case that GWAS loci as yet only explain a fraction of the total genetic variation for any given trait in any given environment. Because statistical power to detect the effect of a given locus is inversely related to its effect size, they are also likely those with the largest effect on the additive variance of a phenotype, and thus may also exhibit a great deal of pleiotropy. In this scenario, selection on a correlated trait could be responsible for the signals we observe, while loci not included in the GWAS set may evolve in a compensatory manner to maintain the same phenotypic optimum.

#### Missing variants

The missing variant problem may also arise simply due to the interaction between demographic history and adaptation. A particularly dramatic example of this is offered by our analysis of skin pigmentation associated loci, where the alleles found by a GWAS in the Cape Verde population completely fail to predict the skin pigmentation of East Asians and Native Americans. This reflects the fact that the alleles responsible for light skin pigmentation in those populations are not variable in Cape Verde due to the independent and convergent adaptive evolution of light skin pigmentation in the ancestral East Asian population (Norton *et al.*, 2006). It is unclear how prevalent convergent evolution by geographically separated populations is, but there a number of examples (Ffrench-Constant *et al.*, 2000; Arendt and Reznick, 2008; Pearce *et al.*, 2009; Novembre and Di Rienzo, 2009), and theory suggests that it might be surprisingly easy for traits with a more moderate number of loci contributing to variation (Ralph and Coop, 2010).

While such cases should not create a false signal of selection if only drift is involved, they do complicate the interpretation of positive signals. As an illustration, when we take a subset of all the Eurasian HGDP populations we see a significant correlation between genetic skin pigmentation score and longitude (*r*^2^ = 0.15, *p* = 0.015), despite the fact that no such phenotypic correlation exists. While the wrong interpretation is easy to avoid here because we have a good understanding of the true phenotypic distribution, for the majority of GWAS studies such complications will be subtler and so care will have to be taken in their interpretation.

#### Loss of constraint and mutational pressure

One further complication in the interpretation of our results is in how loss of constraint may play a role in driving apparent signals of local adaptation. Traits evolving under uniform stabilizing selection across all populations should be less variable than predicted by our covariance model of drift, due to negative covariances among loci, and so should be underrepresented in the extreme tails of our environmental correlation statistics and the upper tail of *Q_X_*. As such, loss of constraint (i.e. weaker stabilizing selection in some populations than others), should not on its own create a signal of local adaptation. While the loci underpinning the phenotype can be subject to more drift in those populations, there is no systemic bias in the direction of this drift, and thus it should be well accommodated by our model. Loss of constraint, therefore, will not tend to create significant environmental correlations or systematic covariance between alleles of like effect.

An issue may arise, however, when loss of constraint is paired with biased mutational input (i.e. new mutations are more likely to push the phenotype in one direction than another; Zhang and HILL, 2008) or if it is asymmetric (i.e. stabilizing selection is relaxed on one tail of the phenotypic distribution, but not the other). Under these two scenarios, mutations that push the phenotype in one particular direction would tend to drift up in frequency in populations with loss of constraint, creating systematic trends and positive covariance among like effect alleles at different loci, and resulting in a positive signal under our framework. While one would be mistaken to assume that the signal was necessarily that of recent positive directional selection, these scenarios do still imply that selection pressures on the genetic basis of the phenotype vary across space. Positive tests under our methods are thus fairly robust in being signals of differential selection among populations, but are themselves agnostic about the specific processes involved. Further work is needed to establish whether these scenarios can be distinguished from recent directional selection based on only allele frequencies and effect sizes, and as always, claims of recent adaptation should be supported by multiple lines of evidence beyond those provided by population genomics alone.

#### Limited evidence of adaptation on the genetic basis of variation in type 2 diabetes suspectibility

Given the strong interest and history of investigation into whether selection has shaped phenotypic variation and disease risk, even the cases where we find limited evidence of local adaptation are potentially interesting. For example, a number of investigators have claimed that individual GWAS loci for Type 2 Diabetes show signals of selection (Helgason *et al.*, 2007; Hancock *et al.*, 2008, 2010; Klimentidis *et al.*, 2010), a fact that is seen as support for the idea that genetic variation for type 2 diabetes risk has been shaped by local adaptation, potentially consistent with the thrifty genotype hypothesis (Neel, 1962, and variants on this hypothesis). However, we find no evidence of environmental correlations and only a very subtle signal of significant differentiation for type 2 diabetes genetic risk in the Europe/Middle East comparison.

This suggests that local adaptation has not had a large role in shaping the present day world-wide distribution of type 2 diabetes susceptibility alleles (as mapped to date in Europe). One explanation of this discrepancy is that it seems biologically unrealistic that the phenotype of type 2 diabetes susceptibility would exhibit strong adaptive differentiation. Rather, local adaptation may be shaping some pleiotropically related phenotype (which shares only some of the loci involved). However, as seen in Figure 1, our methods have better power than single locus statistics, even when the genetic correlation between the GWAS phenotype and the phenotype under selection is weak. As such our results are more consistent with the idea that local adaptation has had little influence on the genetic basis of type 2 diabetes or a closely correlated phenotype, based on our current (admittedly poor) understanding of the genetic basis of this phenotype. This is not to say that some of the patterns seen at individual type 2 diabetes loci are not due to selection, but rather that is seem unlikely that they have been selected for due to their role in variation in a phenotype closely related to type 2 diabetes.

#### Future directions

In this article we have focused on methods development and so have only just scratched the surface of the wide-range of populations and phenotypes that these types of methods could be applied to. Of particular interest is the possibility of applying these methods to GWAS performed in other species where the ecological determinants of local adaptation are better understood (Atwell *et al.*, 2010; Fournier-Level *et al.*, 2011).

One substantial difficulty with our approach, particularly in its application to other organisms, is that genome-wide association studies of highly polygenic phenotypes require very large sample sizes to map even a fraction of the total genetic variance. One promising way to partially sidestep this issue is by applying methods recently developed in animal breeding. In these genomic prediction/selection approaches, one does not attempt to map individual markers, but instead concentrates on predicting an individual’s genetic value for a given phenotype using all markers simultaneously (Meuwissen *et al.*, 2001; Hayes *et al.*, 2009; Meuwissen *et al.*, 2013). This is accomplished by fitting simple linear models to genome-wide genotyping data, in principle allowing common SNPs to tag the majority of causal sites throughout the genome without attempting to explicitly identify them (see Zhou *et al.*, 2013, for an overview of approaches). These methods have been applied to a range of species, including humans (Yang *et al.*, 2010; Davies *et al.*, 2011; Yang *et al.*, 2011; Lee *et al.*, 2012; de los Campos *et al.*, 2010, 2012, 2013), demonstrating that these predictions can potentially explain a relatively high fraction of the additive genetic variance within a population (and hence much of the total genetic variance). As these predictions are linear functions of genotypes, and hence allele frequencies, we might be able to predict the genetic values of sets of closely related populations for phenotypes of interest and apply very similar methods to those developed here. Such an approach may allow us to assess signals of local adaptation acting on the genetic basis of a large fraction of the total genetic variance for traits across a range of species. However, we note that if the only goal is to establish evidence for local adaptation in a given phenotype, then common garden experiments and traditional *Q_ST_*-based analyses (or intermediate approaches which use genome wide data to estimate relationships among populations but still focus on phenotypic measurements as opposed to explicitly genetic estimates, e.g. Ovaskainen *et al.*, 2011) may offer a simpler approach in species where such comparisons are possible.

As discussed in various places above, it is unlikely that all of the loci underpinning the genetic basis of a trait will have been subject to the same selection pressures, due to their differing roles in the trait and their pleiotropic effects. One potential avenue of future investigation is whether, given a large set of loci involved in a trait we can identify sets of loci in particular pathways or with a particular range of effect sizes that drive the signal of selection on the additive genetic basis of a trait.

Another promising extension of our approach is to deal explicitly with multiple correlated phenotypes. With the increasing number of GWAS efforts both empirical and methodological work are beginning to focus on understanding the shared genetic basis of various phenotypes. This raises the possibility that we may be able to disentangle the genetic basis of which phenotypes are more direct targets of selection, and which are responding to correlated selection on these direct targets (see Kremer *et al.*, 1997; Blows, 2007; Chenoweth and Blows, 2008, for progress along these lines using *Q_ST_*). Such tools may also offer a way of incorporating GxE interactions, as multiple GWAS for the same trait in different environments could be treated as correlated traits.

As association data for a greater variety of populations, species, and traits becomes available, we view the methods described out here, along with future extensions, as a productive way forward in developing a quantitative framework to explore the genetic and phenotypic basis of local adaptation.

## 4 Acknowledgements

We would like to thank Gideon Bradburd, Yaniv Brandvain, Luke Jostins, Chuck Langley, Joe Pickrell, Jonathan Pritchard, Peter Ralph, Jeff Ross-Ibarra, Alisa Sedghifar, Michael Turelli and Michael Whitlock for helpful discussion and/or comments on earlier versions of the manuscript.

## 5 Methods

### Mean Centering and Covariance Matrix Estimation

Written in matrix notation, the procedure of mean centering the estimated genetic values and dropping one population from the analysis can be expressed as

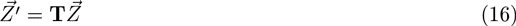

where **T** is an *M* − 1 by *M* matrix with 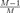 on the main diagonal, and 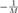 elsewhere.

In order to calculate the corresponding expected neutral covariance structure about this mean, we use the following procedure. Let **G** be an *M* by *K* matrix, where each column is a vector of allele frequencies across the *M* populations at a particular SNP, randomly sampled from the genome according to the matching procedure described below. Let *ε_k_* be the mean allele frequency in column *k* of **G**, and let **S** be a diagonal matrix such that 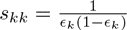 and *s*_*k*≠*i*_ = 0. With this data, we can estimate **F** as

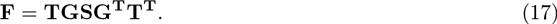

This transformation performs the operation of centering the matrix at the mean value, and rooting the analysis with one population by dropping it from the covariance matrix (the same one we dropped from the vector of estimated genetic values), resulting in a covariance matrix describing the relationship of the remaining *M* − 1 populations. This procedure thus escapes the singularity introduced by centering the matrix at the observed mean of the sample.

Note that as we do not get to observe the population allele frequencies, the entries of **G** are the sample frequencies at the randomly chosen loci, and thus the covariance matrix **F** also includes the effect of finite sample size. Because the noise introduced by the sampling of individuals is uncorrelated across populations (in contrast to that introduced by drift and shared history), the primary effect is to inflate the diagonal entries of the matrix (see the supplementary material of **?**, for discussion).

#### Standardized environmental variable

Given a vector of environmental variable measurements for each population, we apply both the T and Cholesky tranformation as for the estimated genetic values

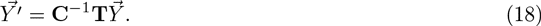

This provides us with a set of *M* − 1 adjusted observations for the environmental variable which can be compared to the transformed genetic values for inference. This step is necessary as we have rotated the frame of reference of the estimated genetic values, and so we must do the same for the environmental variables to keep them both in a consistent reference frame.

### Identifying Outliers with Conditional MVN Distributions

As described in the Results, we can use our multivariate normal model of relatedness to obtain the expected distribution of genetic values for an arbitrary set of populations, conditional on the observed values in some other arbitrary set.

We first partition our populations into two groups, those for which we want to obtain the expected distribution of genetic values (group 1), and those on which we condition in order to obtain this distribution (group 2). We then re–estimate the covariance matrix such that it is centered on the mean of group 2. This is a necessary step, as the amount of divergence between the populations in group 1 and the mean of group 2 will always be greater than the amount of divergence from the global mean, even under the neutral model, and our covariance matrix needs to reflect this fact in order to make accurate predictions. We can obtain this re-parameterized **F** matrix as follows. If *M* is the total number of populations in the sample, then let *q* be the number of populations in group one, and let *M − q* be the number of populations in group 2. We then define a new **T_R_** matrix such that the *q* columns corresponding the populations in group one have 1 on the diagonal, and 0 elsewhere, while the *M − q* columns corresponding to group two have 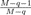 on the diagonal, and 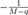 elsewhere. We can then re–estimate a covariance matrix that is centered at the mean of the *M − q* populations in group 2. Recalling our matrices **G** and **S** from (17), this matrix is calculated as

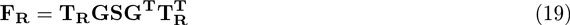

where we write **F_R_** to indicate that it is a covariance matrix that has been re-centered on the mean of group two.

Once we have calculated this re–centered covariance matrix, we can use well known results from multivariate normal theory to obtain the expected joint distribution of the genetic values for group one, conditional on the values observed in group two.

We partition our vector of genetic values and the re–centered covariance matrix such that

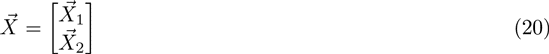

and

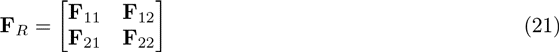

where 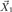 and 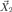 are vectors of genetic values in group 1 and 2 respectively, and **F**_11_, **F**_22_ and 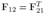 are the marginal covariance matrices of populations within group 1, within group 2, and across the two groups, respectively. Letting 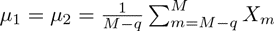 (i.e. the sum of the elements of 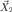), we wish

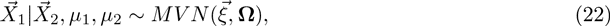

where 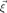 and **Ω** give the expected means and covariance structure of the populations in group 1, conditional on the values observed in group 2. These can be calculated as

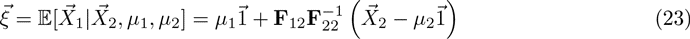

and

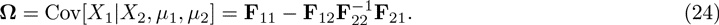

where the one vectors in line (23) are of length *q* and *M − q* respectively.

This distribution is itself multivariate normal, and as such this framework is extremely flexible, as it allows us to obtain the expected joint distribution for arbitrary sets of populations (e.g. geographic regions or continents), or for each individual population. Further,

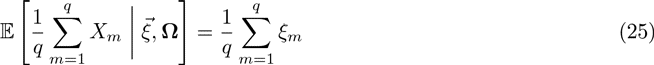

and

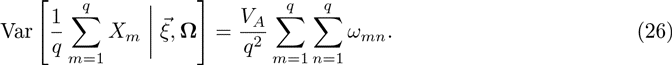

where *ω_nm_* denotes the elements of **Ω**. In words, the conditional expectation of the mean estimated genetic value across group 1 is equal to the mean of the conditional expectations, and its variance is equal to the mean value of the elements of the conditional covariance matrix. As such we can easily calculate a Z score (and corresponding p value) for group one as a whole as

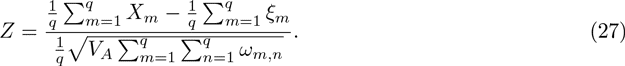

This Z score is a normal random variable with mean zero, variance one under the null hypothesis, and thus measures the divergence of the genetic values between the two populations relative to the null expectation under drift. Note that the observation of a significant Z score in a given population or region cannot necessarily be taken as evidence that selection has acted in that population or region, as selection in the some of the populations on which we condition (especially the closely related ones) could be responsible for such a signal. As such, caution is warranted when interpreting the output of these sort of analyses, and is best done in the context of demographic history information (e.g. from TreeMix, **?**, which uses the same basic model to draw inference about population history as we use to control for it and to generate conditional distributions).

### The Linear Model at the Individual Locus Level

It is natural to ask about the relationship between our environmental correlation tests performed at the genetic value level, and tests that could be performed on individual SNPS and then aggregated.

As noted in (8), we can obtain for each underlying locus a set of transformed allele frequencies, which have passed through the same transformation as the estimated genetic values. We assume that each locus *ℓ* has a regression coefficient

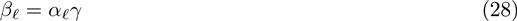

where *γ* is shared across all loci

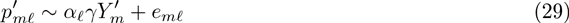

where the *e_mℓ_* are independent and identically distributed residuals. We can find the maximum likelihood estimate 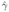 by treating 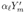 as the linear predictor, and taking the regression of the combined vector 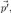, across all populations and loci, on the combined vector 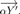. As such

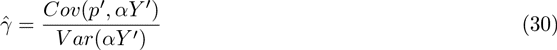

we can decompose this into a sum across loci

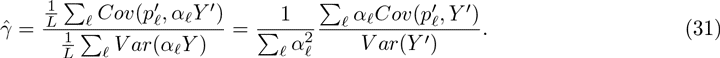

As noted in (8), our transformed genetic values can be written as

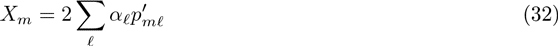

and so the estimated slope 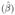 of our regression (*X* = *βY′*) is

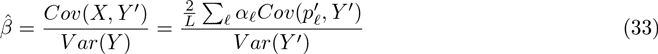

Comparing these equations, the mean regression coefficient at the individual loci (31) and the regression coefficient of the estimated genetic values (33) are proportional to each other via a constant that is given by the sum over effect sizes 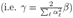, and thus the two approaches are equivalent.

### HGDP data and imputation

We used imputed allele frequency data in the HGDP, where the imputation was performed as part of the phasing procedure of Pickrell *et al.* (2009), as per the recommendations of Conrad *et al.* (2006). We briefly recap their procedure here.

Phasing and imputation were done using fastPHASE (Scheet and Stephens, 2006), with the settings that allow variation in the switch rate between subpopulations. The populations were grouped into subpopulations corresponding to the clusters identified in Rosenberg *et al.* (2002). Haplotypes from the HapMap YRI and CEU populations were included as known, as they were phased in trios and are highly accurate. HapMap JPT and CHB genotypes were also included to help with the phasing.

### Choosing null SNPs

Various components of our procedure involve sampling random sets of SNPs from across the genome. While we control for biases in our test statistics introduced by population structure through our **F** matrix, we are also concerned that subtle ascertainment effects of the GWAS process could lead to biased test statistics, even under neutral conditions. We control for this possibility by sampling null SNPs so as to match the joint distribution of certain properties of the ascertained GWAS SNPs. Specifically, we were concerned that the minor allele frequency (MAF) in the ascertainment population, the imputation status of the allele in the HGDP datasets, and the background selection environment experienced at a given locus, as measured by B value (McVicker *et al.*, 2009) might influence the distribution of allele frequencies across populations in ways that we could not predict.

We partitioned SNPs into a three way contingency table, with 25 bins for MAF (i.e. a bin size of 0.02), 2 bins for imputation (either imputed or not), and 10 bins for B value (B values range from 0 to 1, and thus our bin size was 0.1). As such, for each set of null genetic values, we sampled one null SNP from the same cell in the contingency table as each of the GWAS SNPs, and assigned this null SNP the effect size associated with the GWAS SNP it was sampled to match. While we do not assign effect sizes to sampled SNPs used to estimate the covariance matrix **F** (instead simply scaling **F** by a weighted sum of squared effect sizes, which is mathematically equivalent under our assumption that all SNPs have the same covariance matrix), we follow the same sampling procedure to ensure that **F** describes the expected covariance structure of the GWAS SNPs.

For the skin pigmentation GWAS (Beleza *et al.*, 2013) we do not have a good proxy present in the HGDP population, as the Cape Verdeans are an admixed population. Cape Verdeans are admixed with ∼59.53% African ancestry, and 41.47% European ancestry in the sample obtained by Beleza *et al.* (2013) (Beleza, pers. comm., April 8, 2013). As such, we estimated genome wide allele frequencies in Cape Verde by taking a weighted mean of the frequencies in the French and Yoruban populations of the HGDP, such that *p_CV_* = 0.5953*p_Y_* + 0.4147*p_F_*. We then used these estimated frequencies to assign SNPs to frequency bins.

Beleza *et al.* (2013) also used an admixture mapping strategy to map the genetic basis of skin pigmentation. However, if they had only mapped these loci in an admixture mapping setting we would have to condition our null model on having strong enough allele frequency differentiation between Africans and Europeans at the functional loci for admixture mapping to have power (Reich and Patterson, 2005). The fact that Beleza *et al.* (2013) mapped these loci in a GWAS framework allows us to simply reproduce the strategy, and we ignore the results of the admixture mapping study (although we note that the loci and effect sizes estimated were similar). This highlights the need for a reasonably well defined ascertainment population for our approach, a point which we comment further on in the discussion.

## 6 Supplementary materials

**Figure S1:**
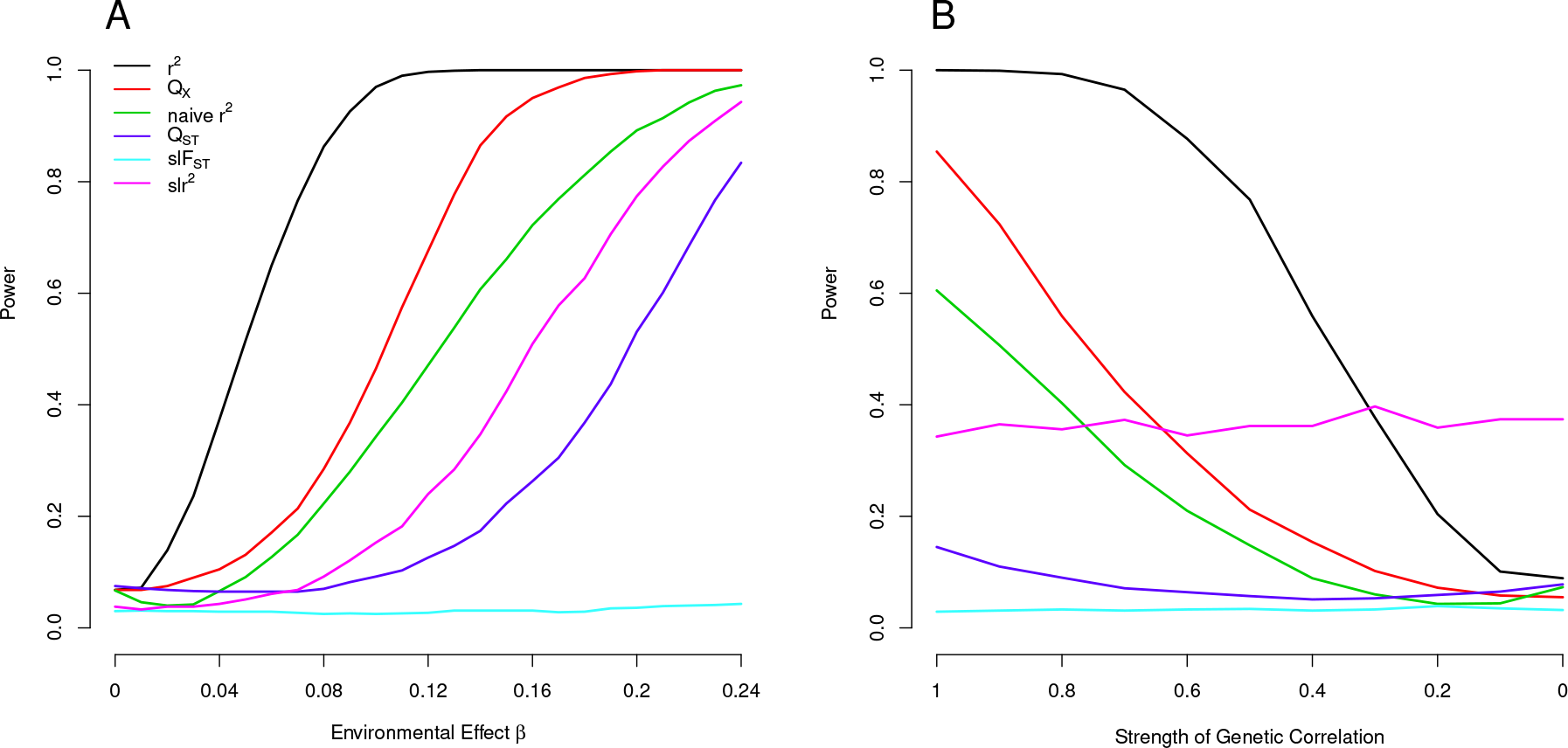
Power of tests described in the main text to detect a signal of selection on the mapped genetic basis of height (Lango Allen *et al.*, 2010) as an increasing function of the strength of selection (A), and a decreasing function of the genetic correlation between height and the selected trait with the effect of selection held constant at *β* = 0.14 (B).

**Figure S2:**
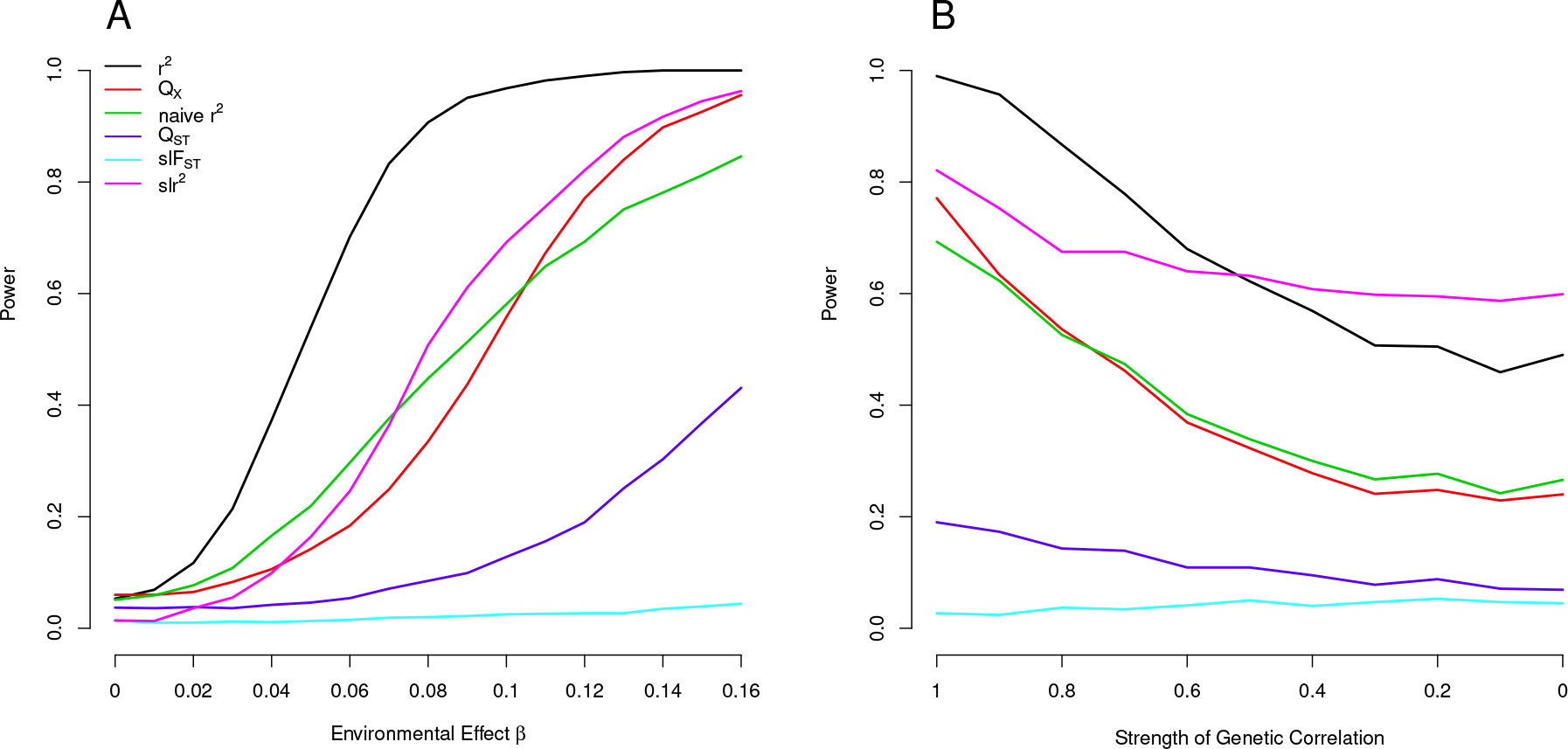
Power of tests described in the main text to detect a signal of selection on the mapped genetic basis of skin pigmentation (Beleza *et al.*, 2013) as an increasing function of the strength of selection (A), and a decreasing function of the genetic correlation between skin pigmentation and the selected trait with the effect of selection held constant at *β* = 0.12 (B).

**Figure S3:**
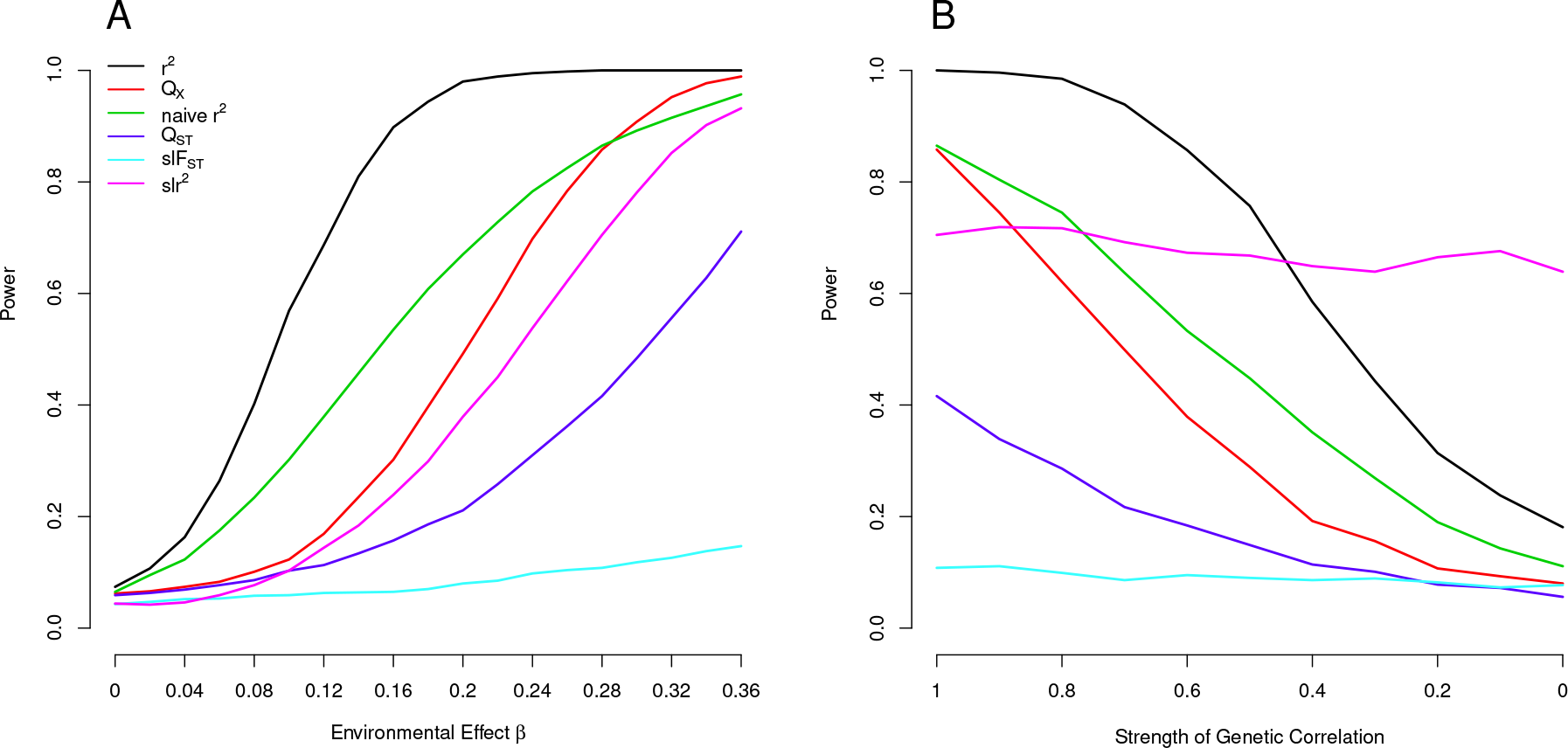
Power of tests described in the main text to detect a signal of selection on the mapped genetic basis of BMI (Speliotes *et al.*, 2010) as an increasing function of the strength of selection (A), and a decreasing function of the genetic correlation between BMI and the selected trait with the effect of selection held constant at *β* = 0.28 (B).

**Figure S4:**
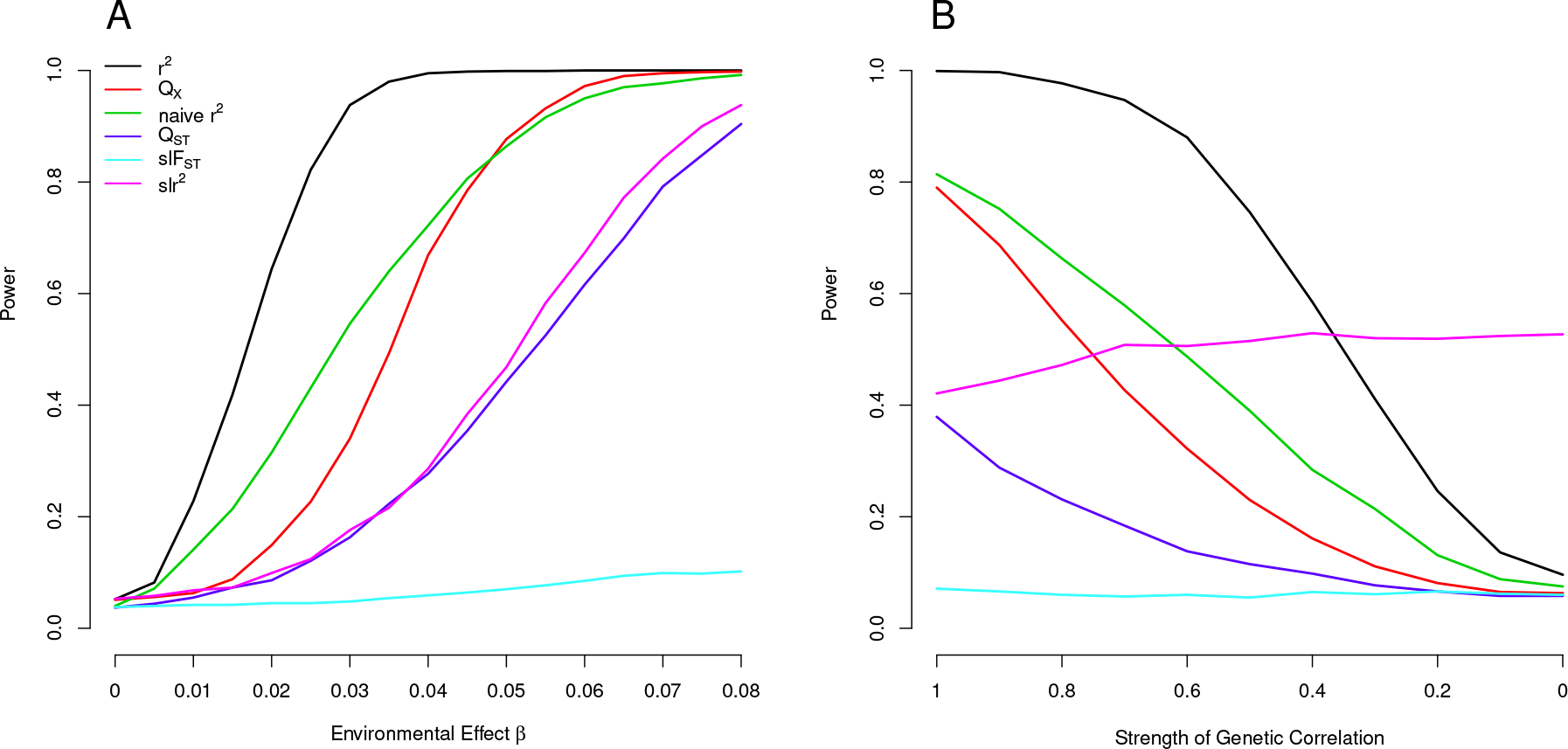
Power of tests described in the main text to detect a signal of selection on the mapped genetic basis of CD (Jostins *et al.*, 2012) as an increasing function of the strength of selection (A), and a decreasing function of the genetic correlation between CD and the selected trait with the effect of selection held constant at *β* = 0.045 (B).

**Figure S5:**
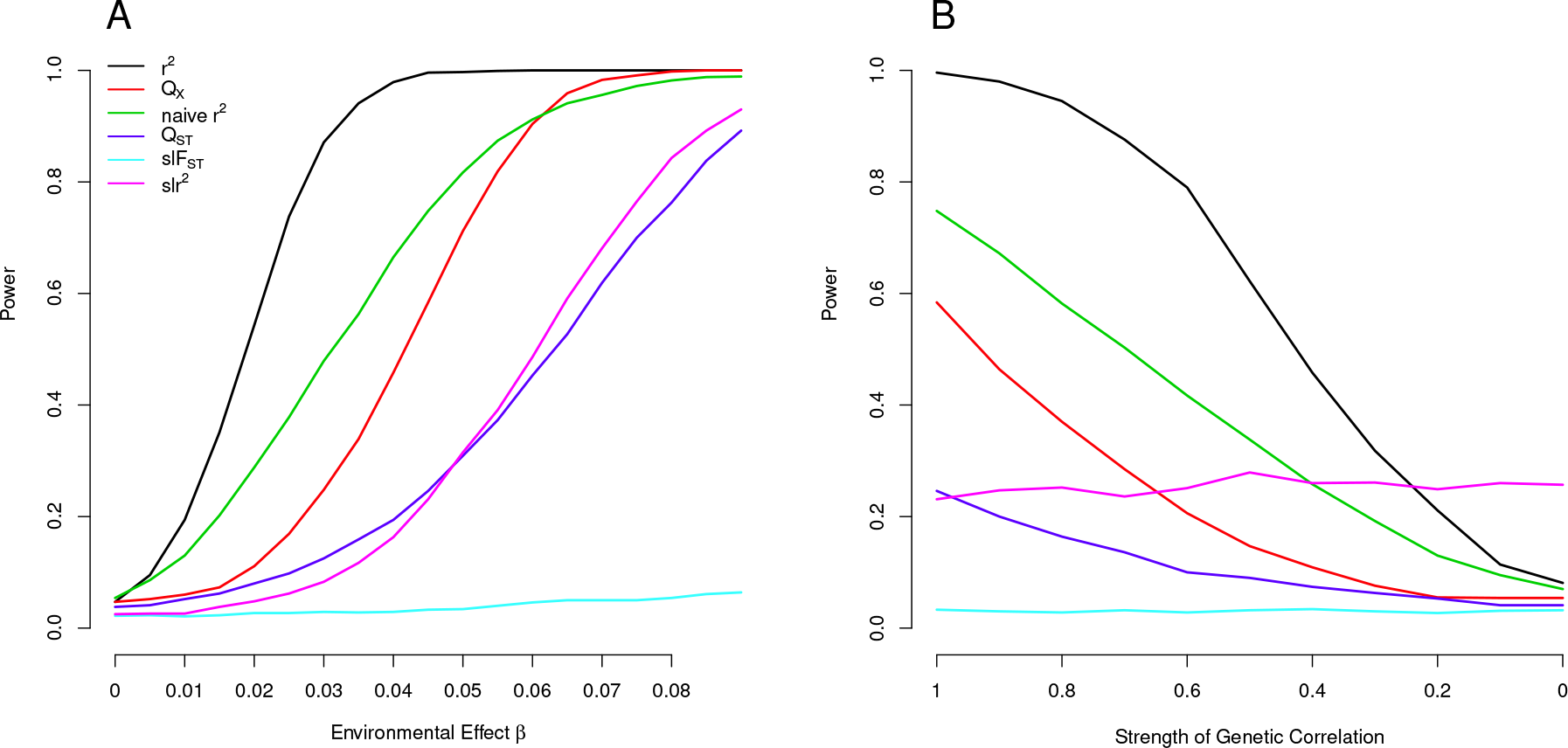
Power of tests described in the main text to detect a signal of selection on the mapped genetic basis of UC (Jostins *et al.*, 2012) as an increasing function of the strength of selection (A), and a decreasing function of the genetic correlation between UC and the selected trait with the effect of selection held constant at *β* = 0.045 (B).

**Figure S6:**
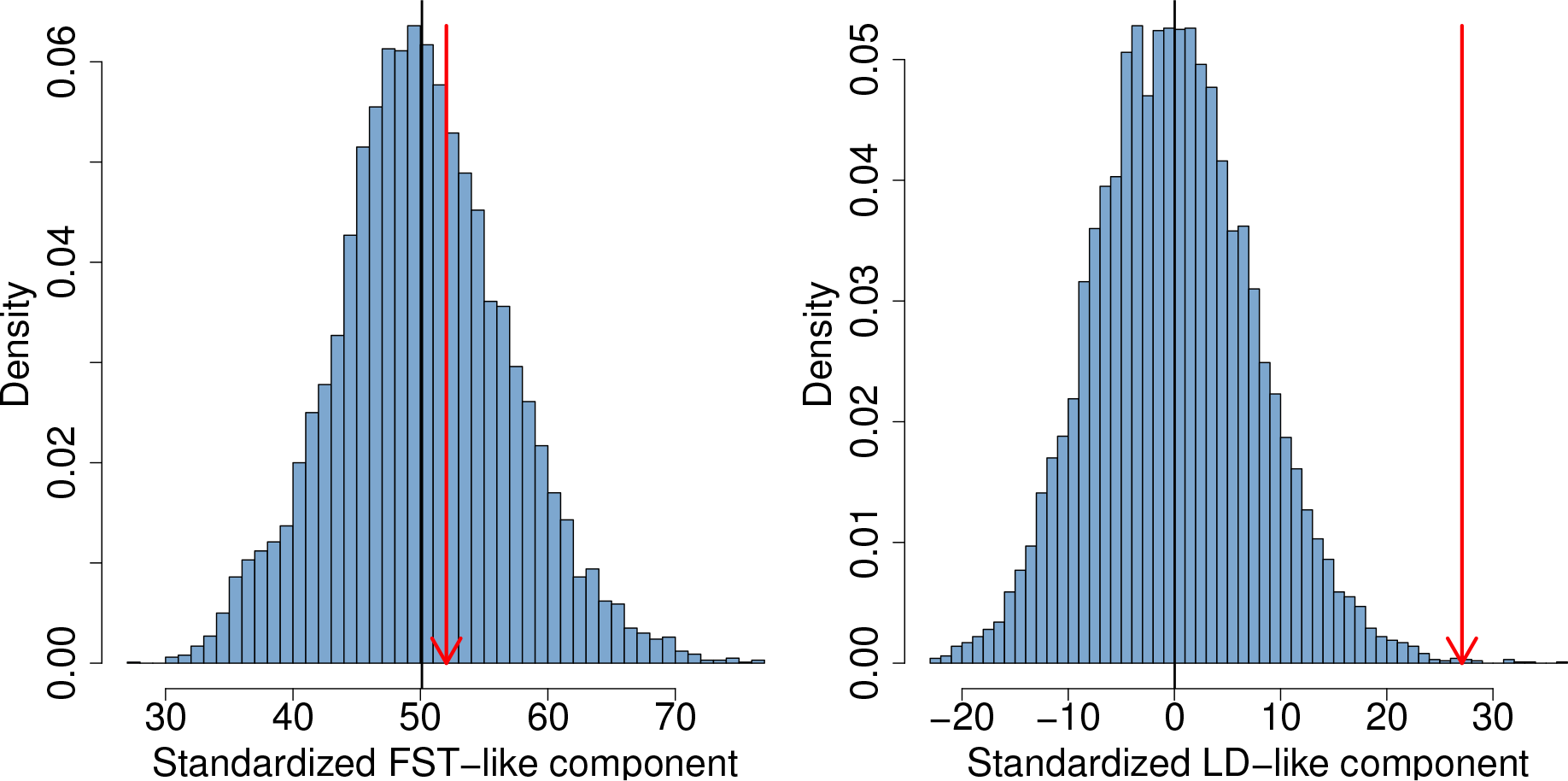
The two components of *Q_X_* for the skin pigmentation dataset, as described by the left and right terms in (14). The null distribution of each component is shows as a histogram. The expected value is shown as a black bar, and the observed value as a red arrow.

**Figure S7:**
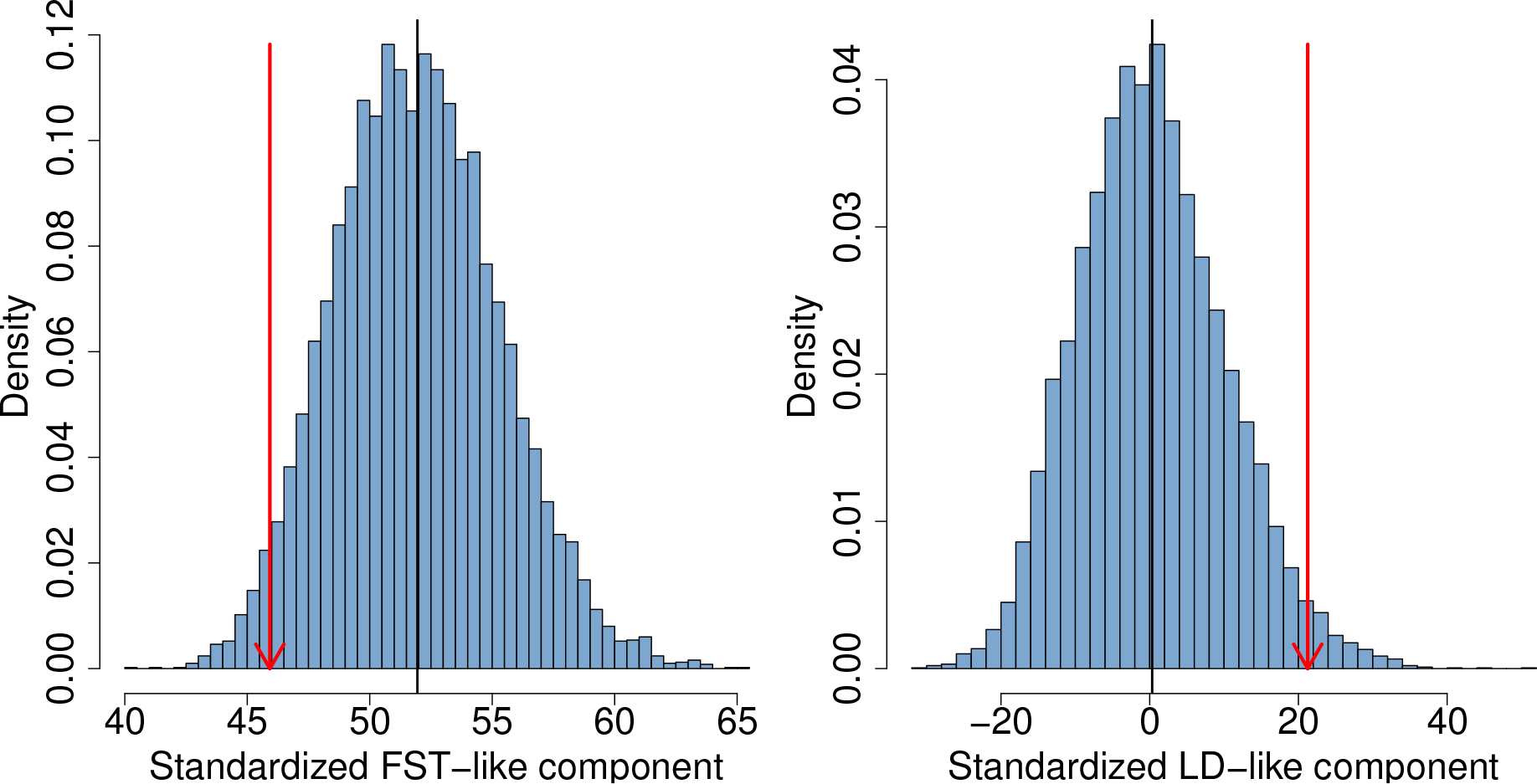
The two components of *Q_X_* for the BMI dataset, as described by the left and right terms in (14). The null distribution of each component is shows as a histogram. The expected value is shown as a black bar, and the observed value as a red arrow.

**Figure S8:**
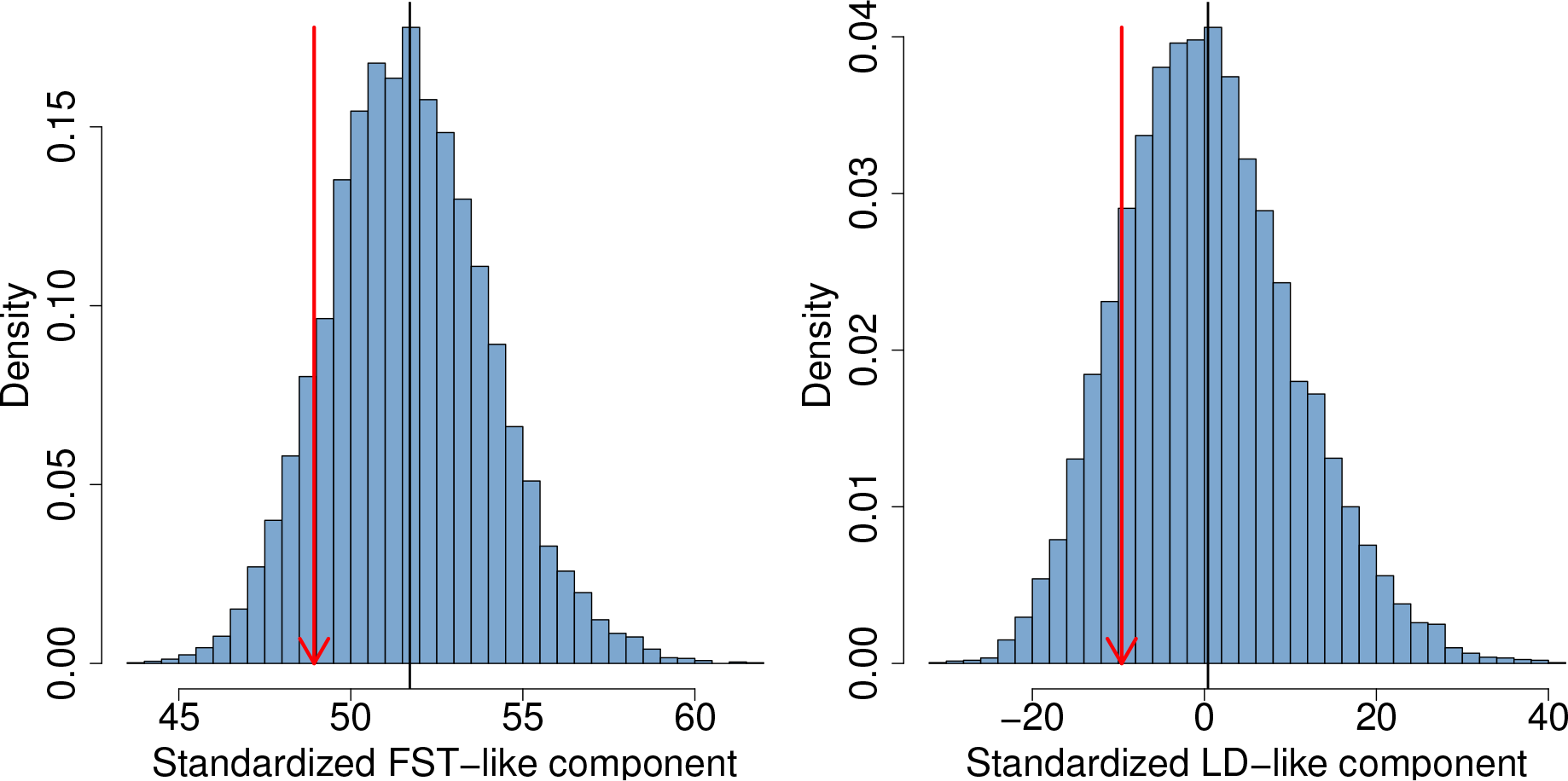
The two components of *Q_X_* for the T2D dataset, as described by the left and right terms in (14). The null distribution of each component is shows as a histogram. The expected value is shown as a black bar, and the observed value as a red arrow.

**Figure S9:**
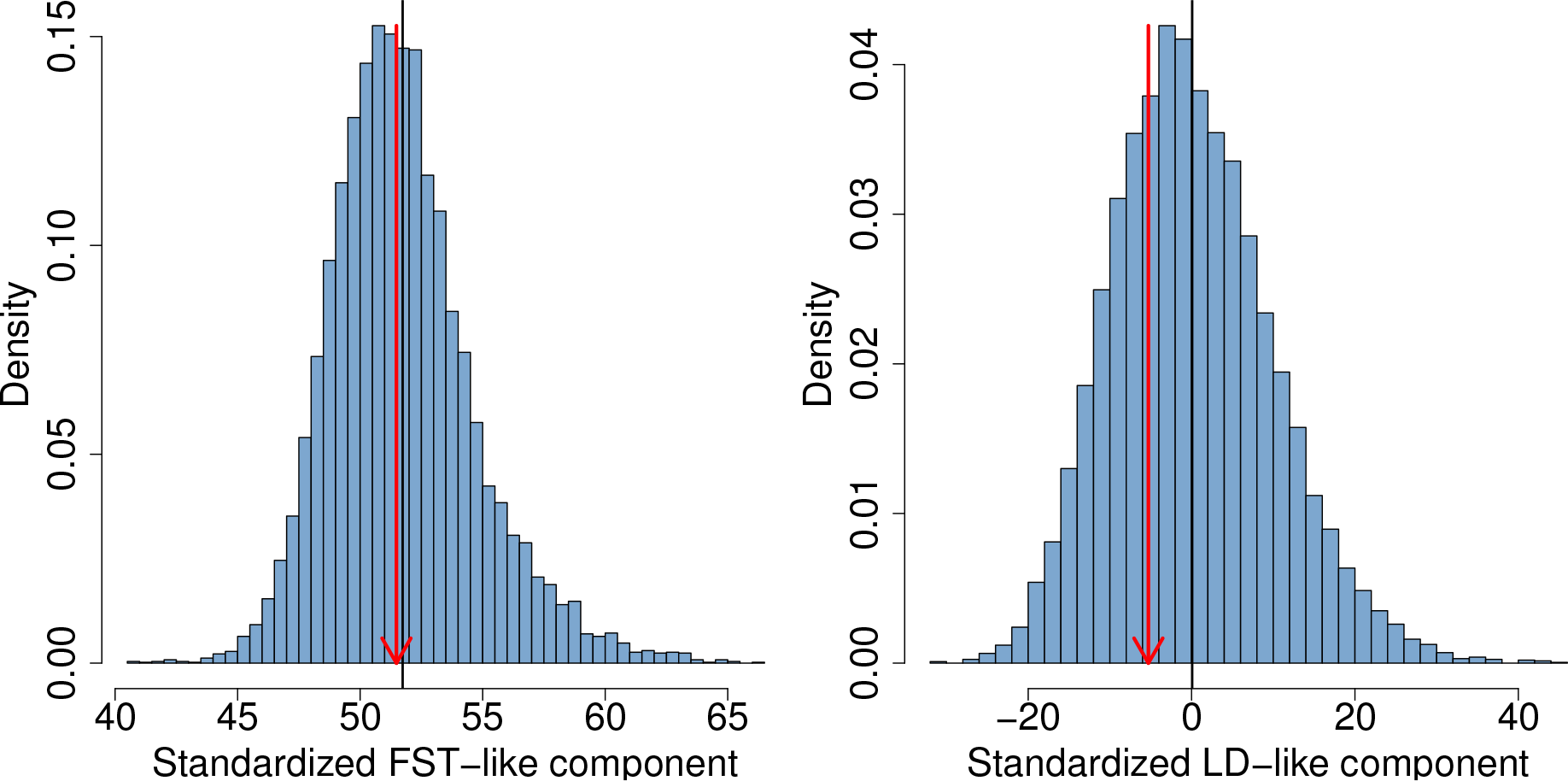
The two components of *Q_X_* for the CD dataset, as described by the left and right terms in (14). The null distribution of each component is shows as a histogram. The expected value is shown as a black bar, and the observed value as a red arrow.

**Figure S10:**
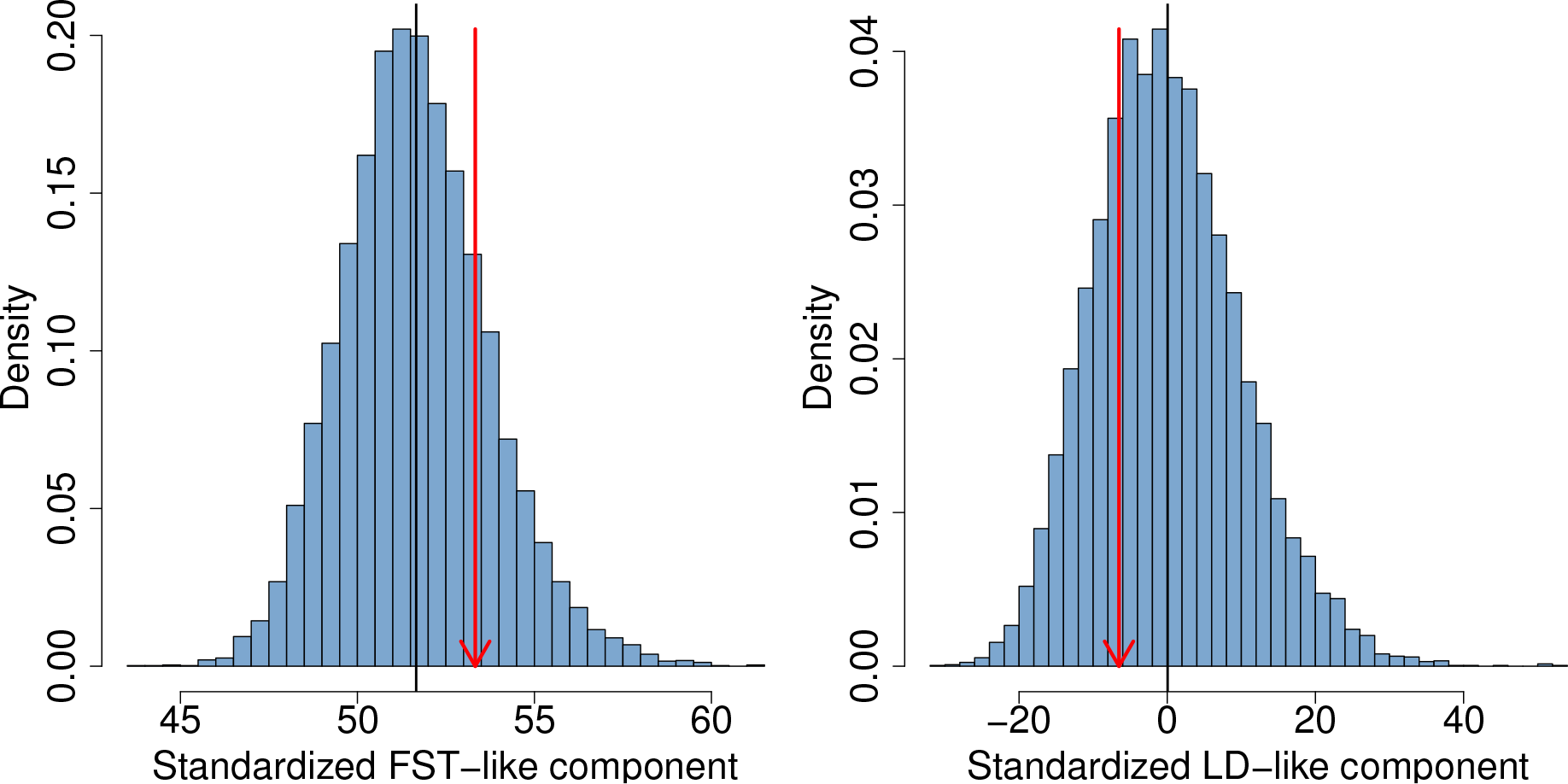
The two components of *Q_X_* for the UC dataset, as described by the left and right terms in (14). The null distribution of each component is shows as a histogram. The expected value is shown as a black bar, and the observed value as a red arrow.

**Figure S11:**
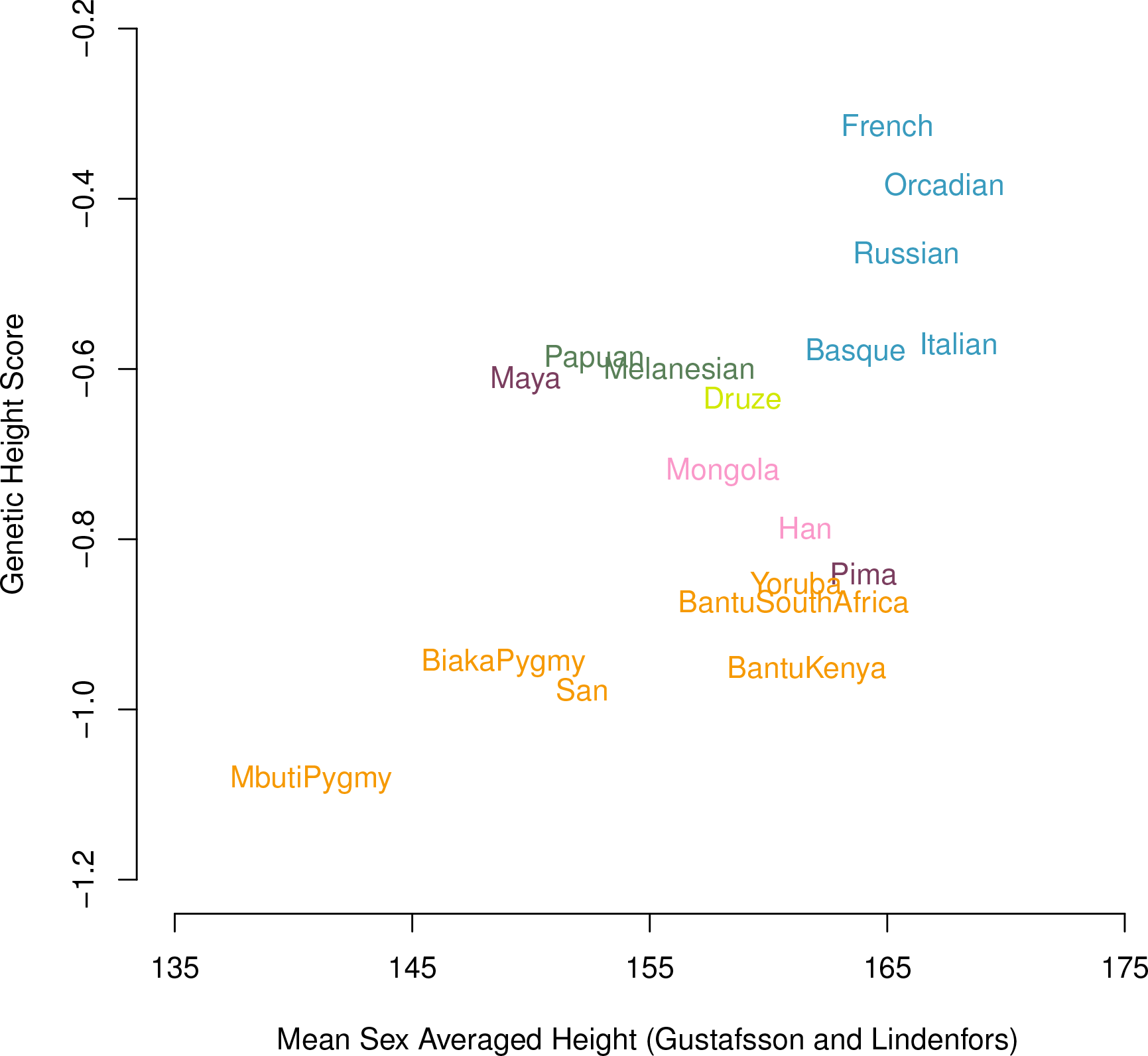
The genetic values for height in each HGDP population plotted against the measured sex averaged height taken from Gustafsson and Lindenfors (2004). Only the subset of populations with an appropriately close match in the named population in Gustafsson and Lindenfors’s (2004) Appendix I are shown, values used are given in Supplementary table **??**

**Figure S12:**
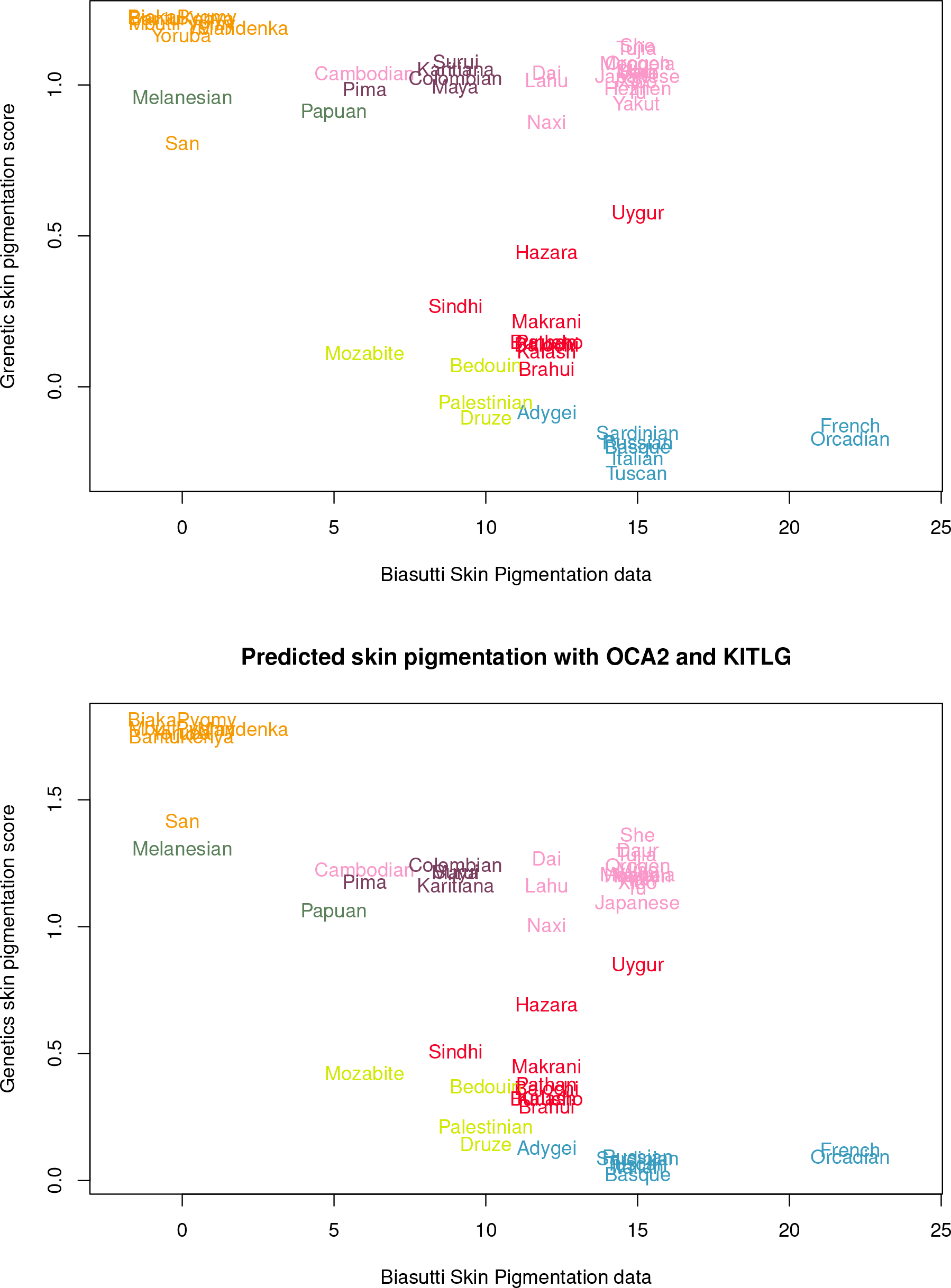
The genetic skin pigmentation score for a each HGDP population plotted against the HGDP populations values on the skin pigmentation index map of Biasutti 1959. Data obtained from Supplementary table of Lao *et al.* (2007). Note that Biasutti map is interpolated, and so values are known to be imperfect. Values used are given in Supplementary table S2

**Figure S13:**
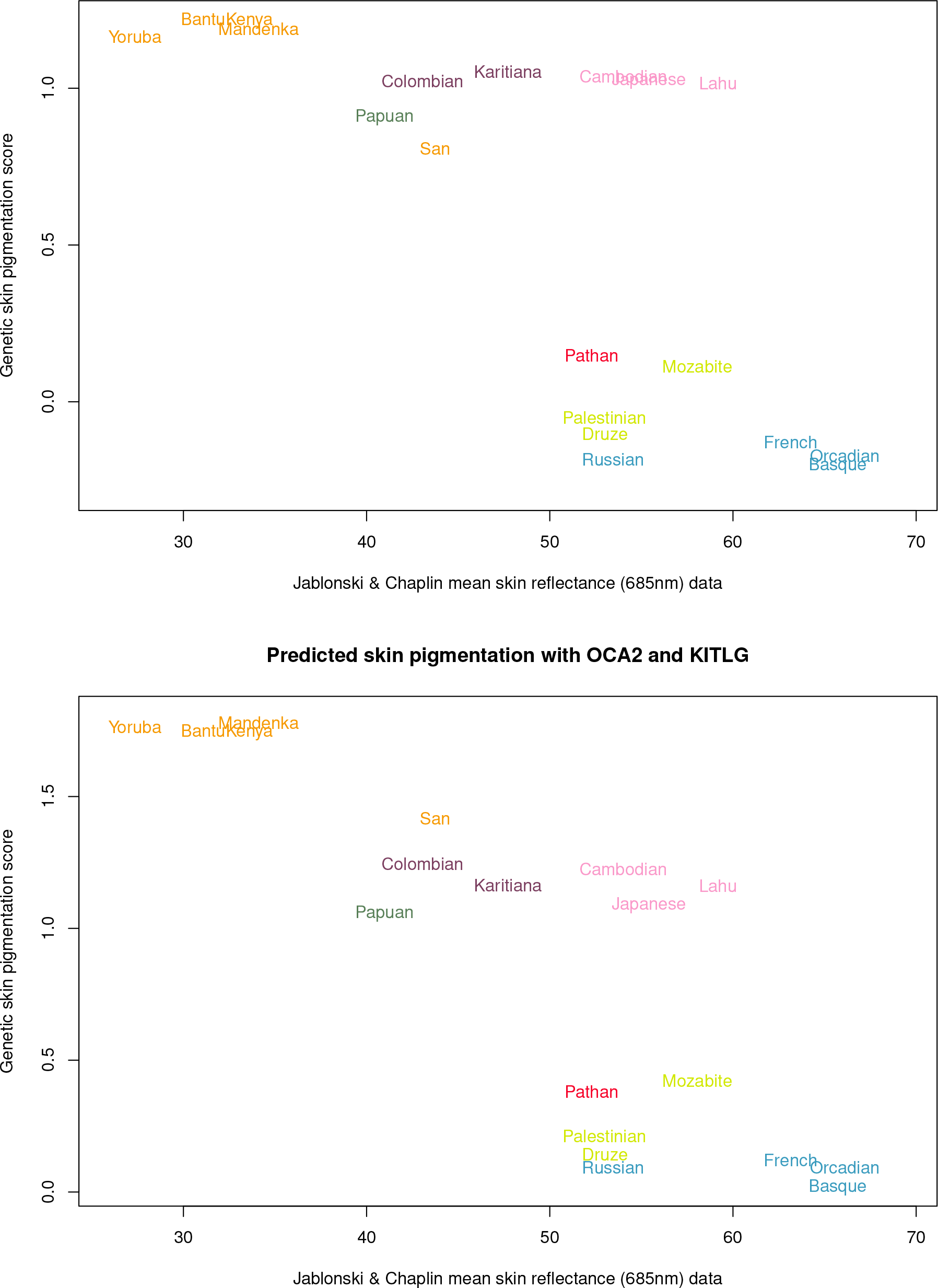
The genetic skin pigmentation score for a each HGDP population plotted against the HGDP populations values from the Jablonski and Chaplin (2000) mean skin reflectance (685 nm) data (their Table 6). Only the subset of populations with an appropriately close match were used as in the Supplementary table of Lao *et al.* (2007). Values and populations used are given in Table S2

**Figure S14:**
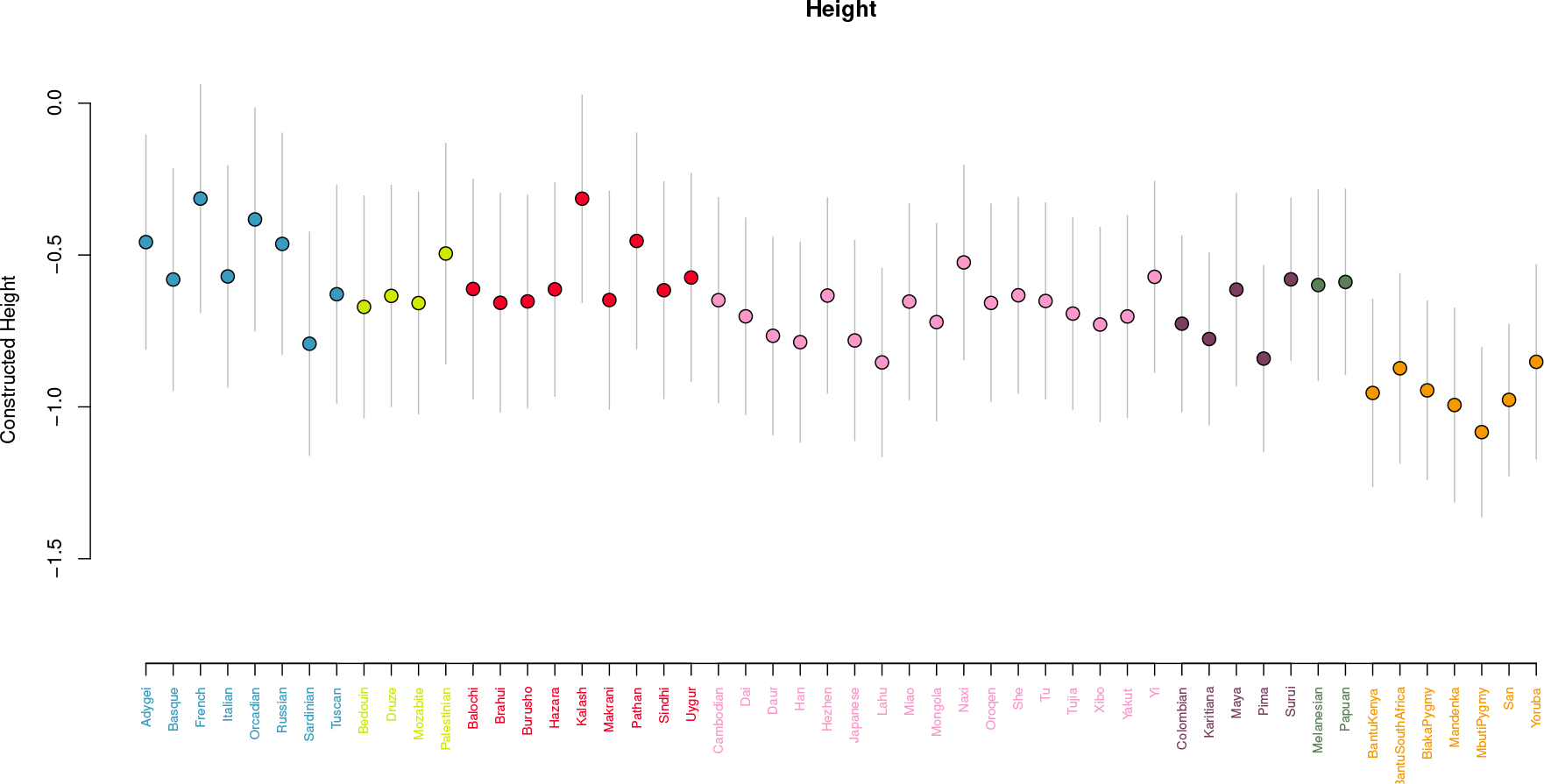
The distribution of genetic height score across all 52 HGDP populations. Grey bars represent the 95% confidence interval for the genetic height score of an individual randomly chosen from that population under Hardy-Weinberg assumptions

**Figure S15:**
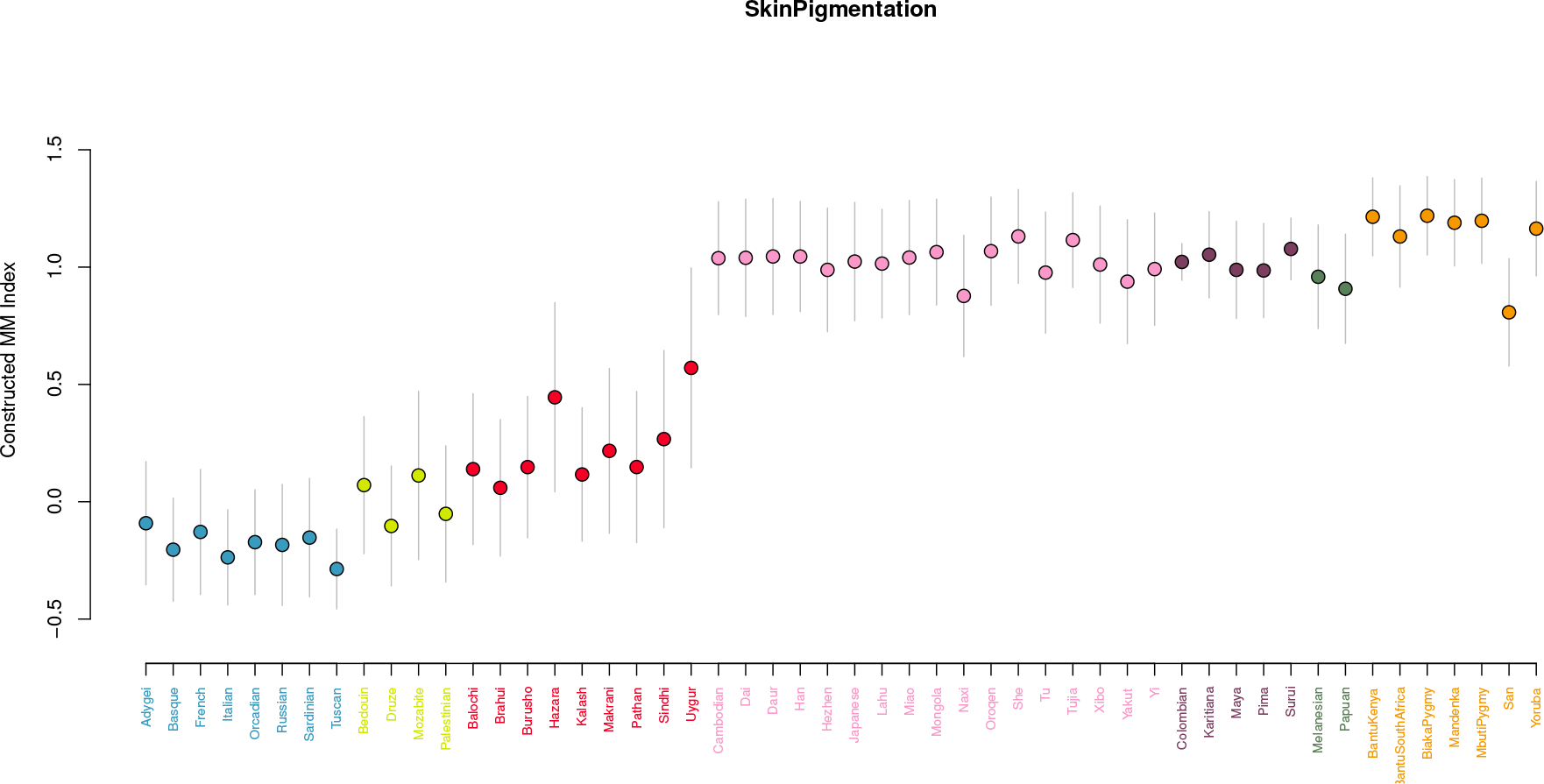
The distribution of genetic skin pigmentation score across all 52 HGDP populations. Grey bars represent the 95% confidence interval for the genetic skin pigmentation score of an individual randomly chosen from that population under Hardy-Weinberg assumptions

**Figure S16:**
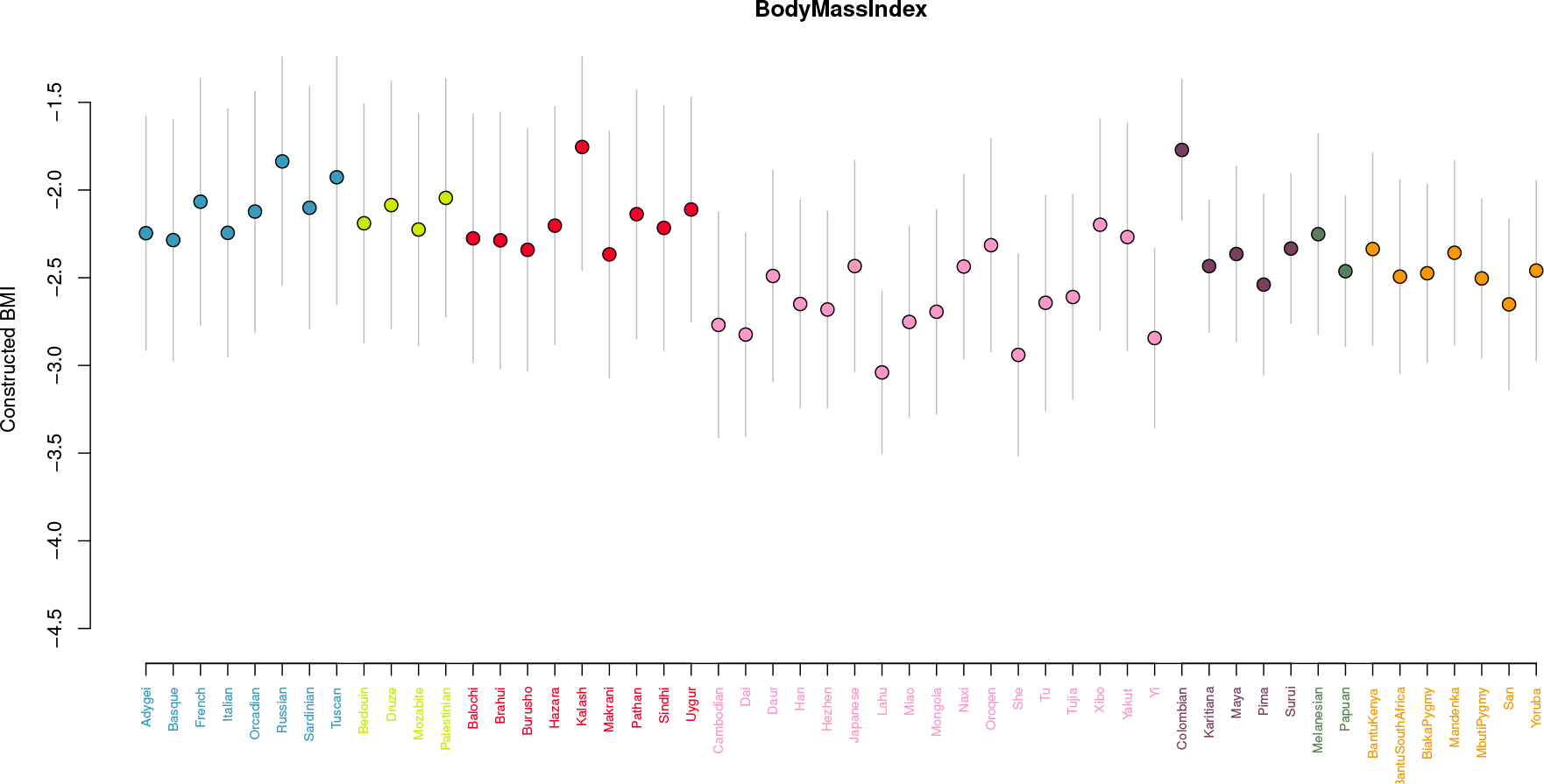
The distribution of genetic BMI score across all 52 HGDP populations. Grey bars represent the 95% confidence interval for the genetic BMI score of an individual randomly chosen from that population under Hardy-Weinberg assumptions

**Figure S17:**
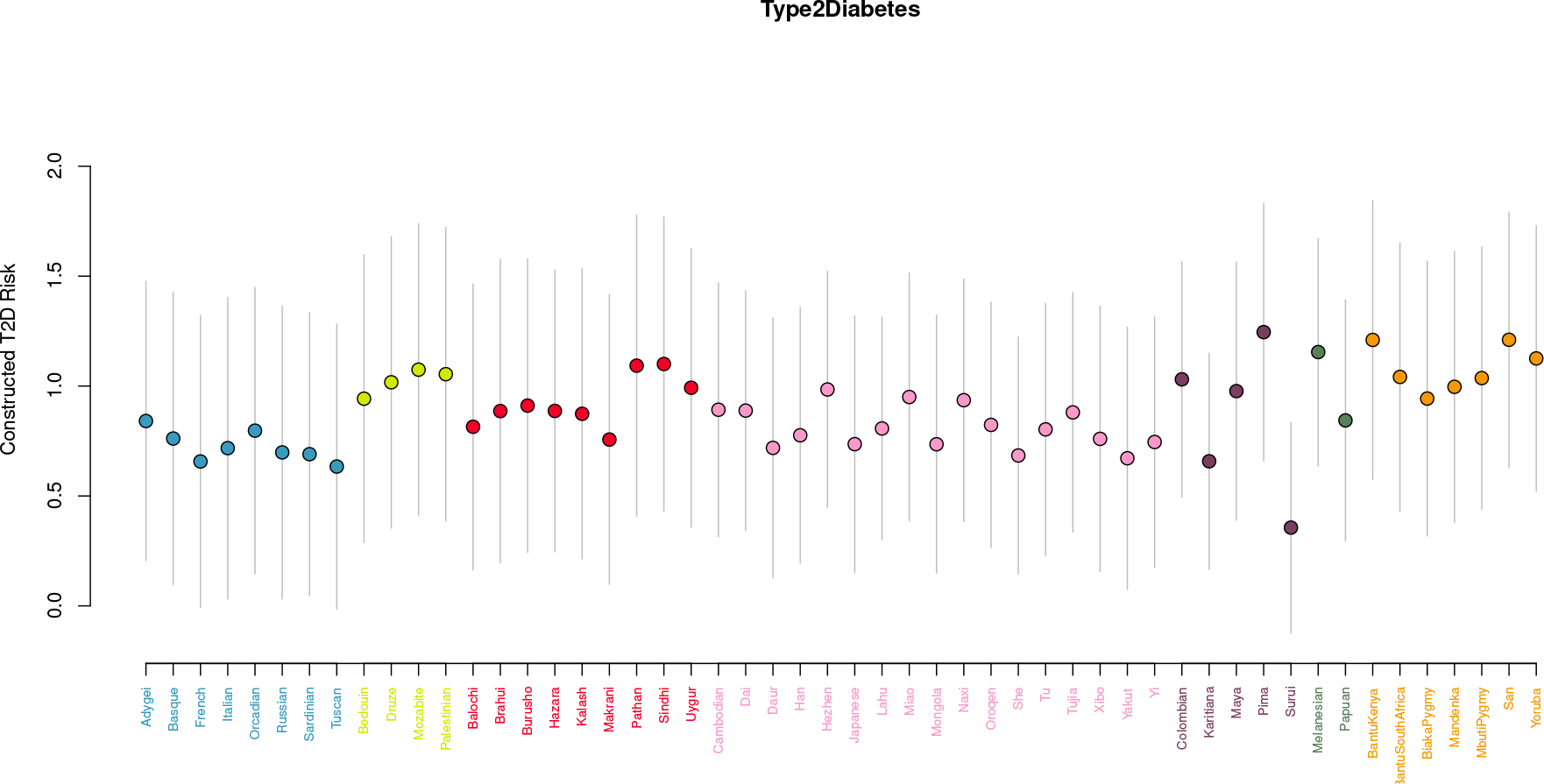
The distribution of genetic T2D risk score across all 52 HGDP populations. Grey bars represent the 95% confidence interval for the genetic T2D risk score of an individual randomly chosen from that population under Hardy-Weinberg assumptions

**Figure S18:**
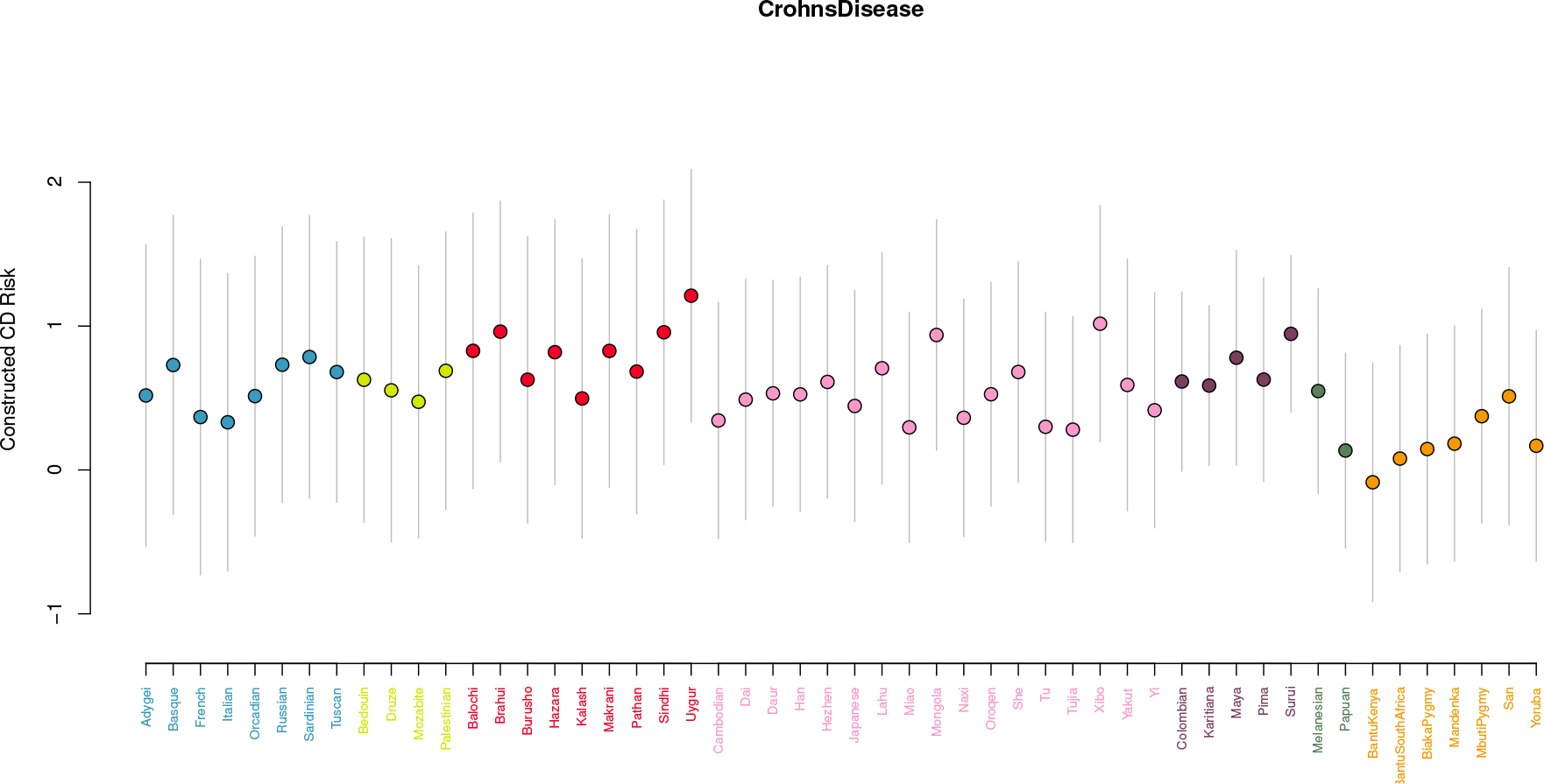
The distribution of genetic CD risk score across all 52 HGDP populations. Grey bars represent the 95% confidence interval for the genetic CD risk score of an individual randomly chosen from that population under Hardy-Weinberg assumptions

**Figure S19:**
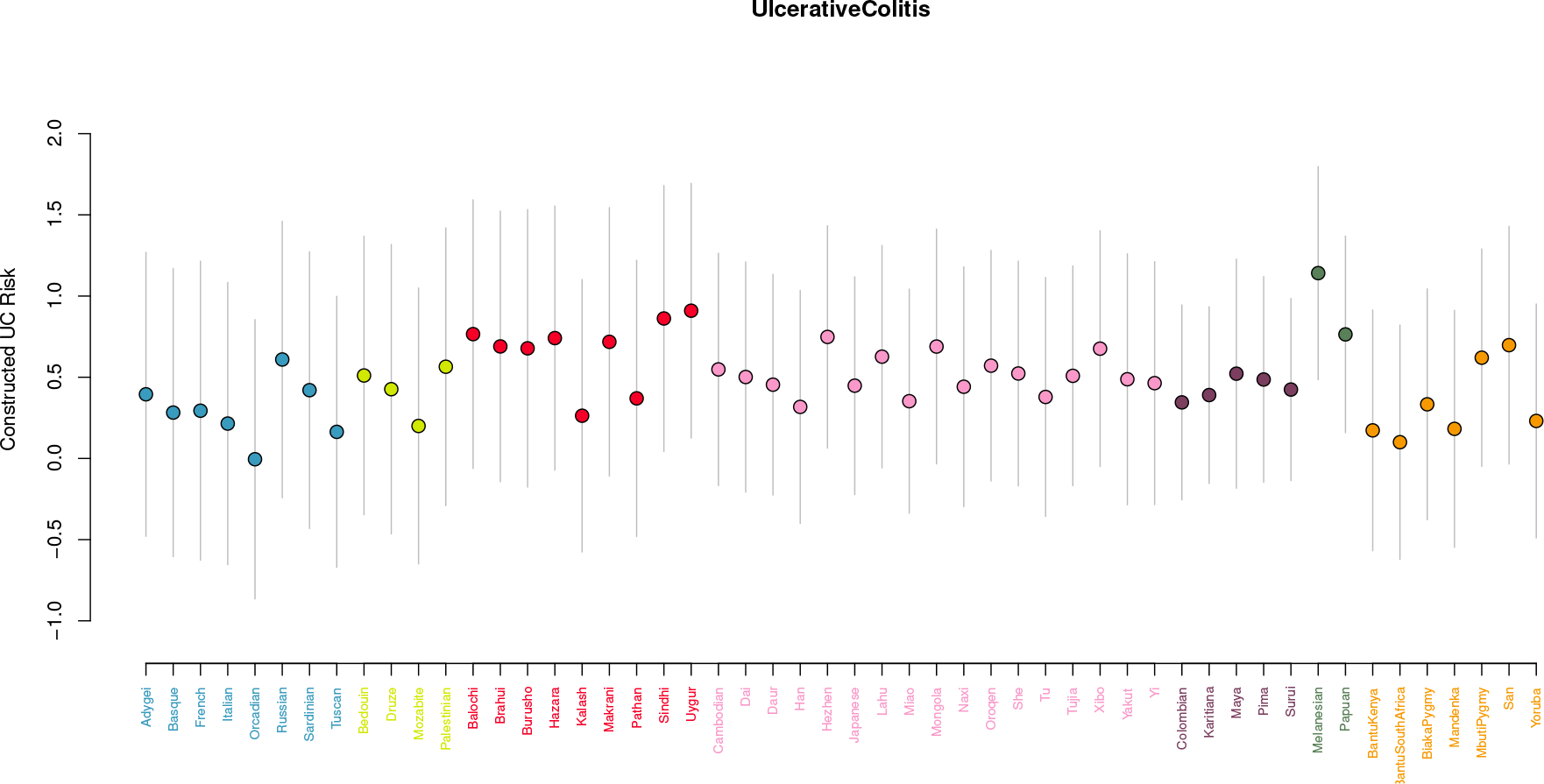
The distribution of genetic UC risk score across all 52 HGDP populations. Grey bars represent the 95% confidence interval for the genetic UC risk score of an individual randomly chosen from that population under Hardy-Weinberg assumptions

**Table S1:** Genetic height scores as compared to true heights for populations with a suitably close match in the dataset of Gustafsson and Lindenfors (2004). See Figure S11 for a plot of genetic height score against sex averaged height.

| Population | Genetic Height Score | Male Height (cm) | Female Height (cm) | Sex Averaged Height (cm) |
| --- | --- | --- | --- | --- |
| BantuKenya | -0.95 | 167.30 | 156.00 | 161.65 |
| BantuSouthAfrica | -0.87 | 166.10 | 156.00 | 161.05 |
| Basque | -0.58 | 170.00 | 157.30 | 163.65 |
| BiakaPygmy | -0.95 | 152.70 | 145.00 | 148.85 |
| Druze | -0.63 | 165.60 | 152.20 | 158.90 |
| French | -0.31 | 169.60 | 160.40 | 165.00 |
| Han | -0.79 | 167.10 | 156.00 | 161.55 |
| Italian | -0.57 | 174.00 | 162.00 | 168.00 |
| Maya | -0.61 | 156.80 | 142.80 | 149.80 |
| MbutiPygmy | -1.08 | 144.19 | 137.35 | 140.77 |
| Melanesian | -0.60 | 162.10 | 150.40 | 156.25 |
| Mongola | -0.72 | 164.83 | 151.33 | 158.08 |
| Orcadian | -0.38 | 173.90 | 160.90 | 167.40 |
| Papuan | -0.59 | 155.97 | 149.41 | 152.69 |
| Pima | -0.84 | 171.00 | 157.00 | 164.00 |
| Russian | -0.46 | 171.80 | 159.80 | 165.80 |
| San | -0.98 | 157.70 | 146.60 | 152.15 |
| Yoruba | -0.85 | 167.50 | 155.00 | 161.25 |

**Table S2:**
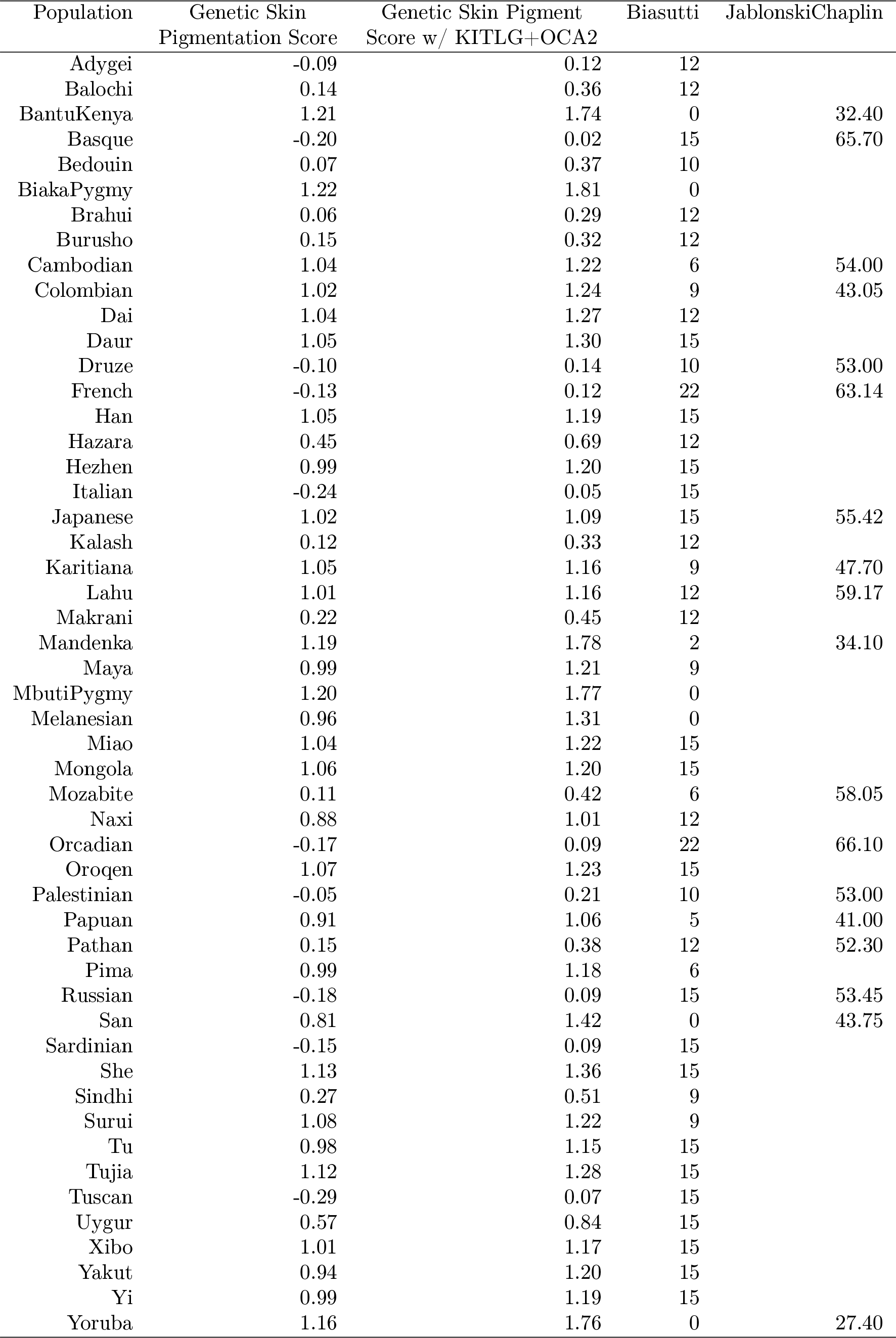
Genetic skin pigmentation score as compared to values from Biasutti (Parra *et al.*, 2004; Lao *et al.*, 2007) and Jablonski and Chaplin (2000). We also calculate a genetic skin pigmentation score including previously reported associations at KITLG and OCA2 for comparisson. See also Figures S12 and S13.

**Table S3:**
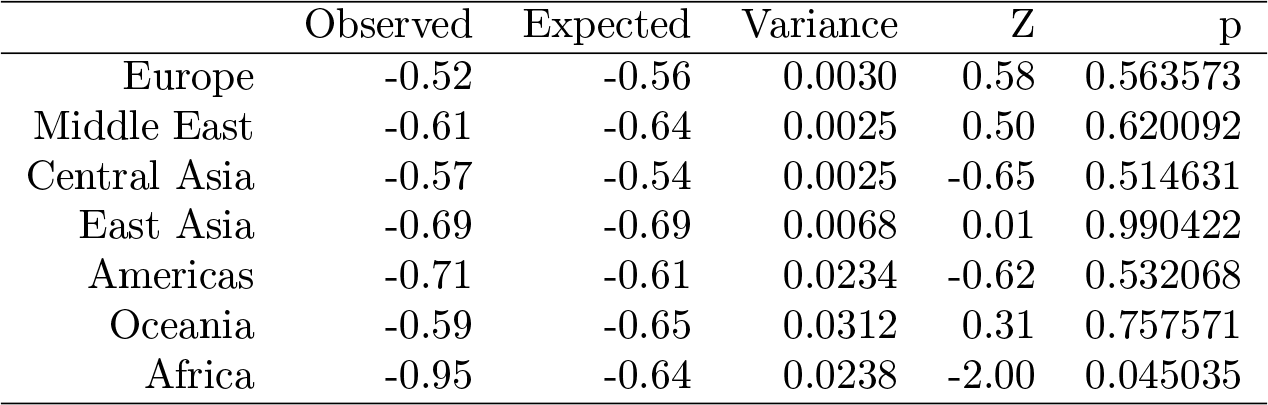
Conditional analysis at the regional level for the height dataset

**Table S4:**
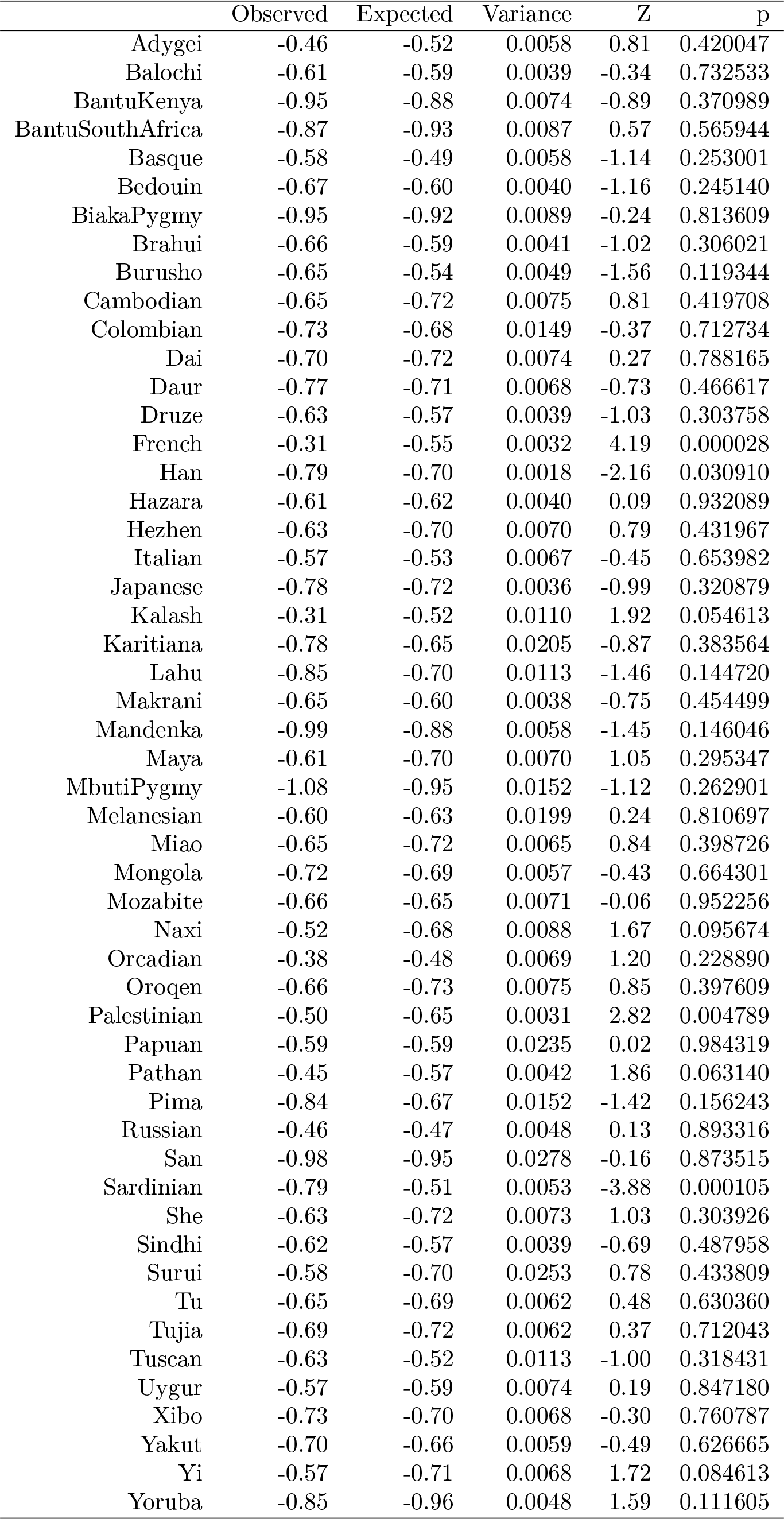
Conditional analysis at the individual population level for the height dataset

**Table S5:**
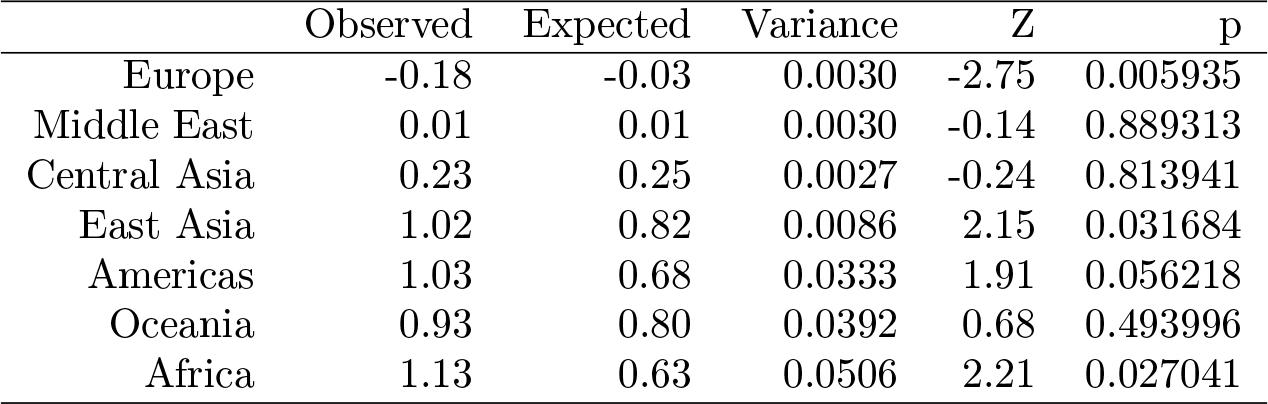
Conditional analysis at the regional level for the skin pigmentation dataset

**Table S6:**
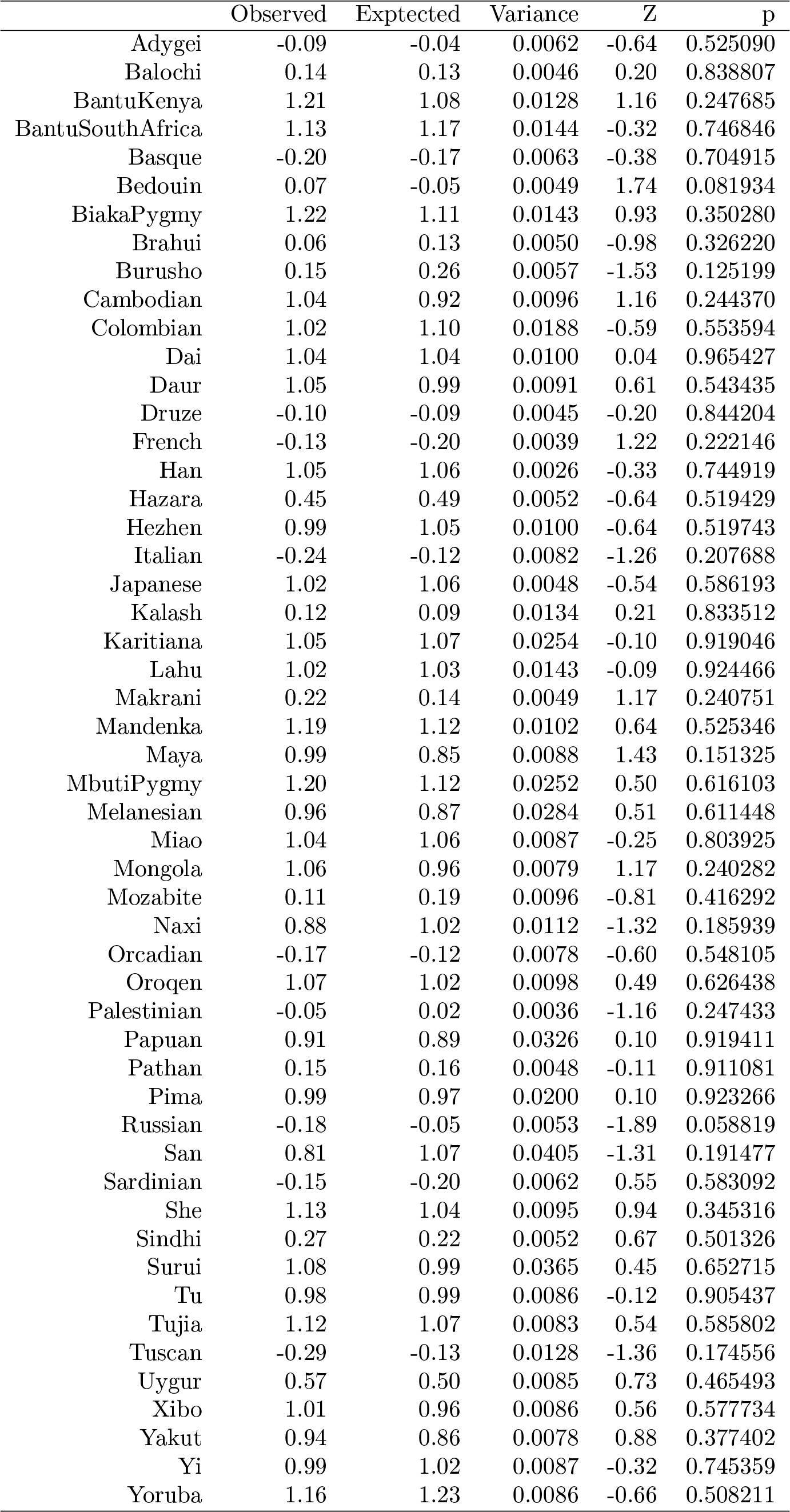
Conditional analysis at the individual population level for the skin pigmentation dataset

**Table S7:**
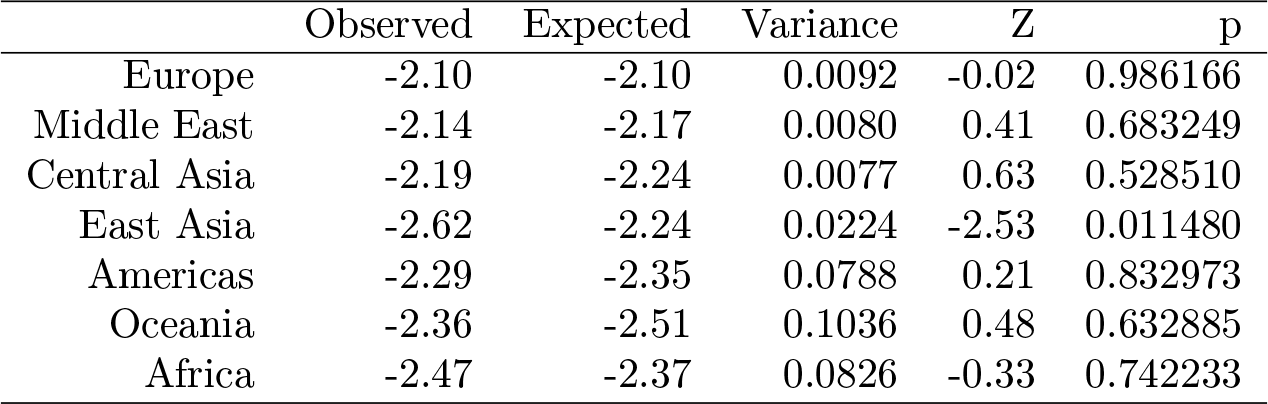
Condtional analysis at the regional level for the BMI dataset

**Table S8:**
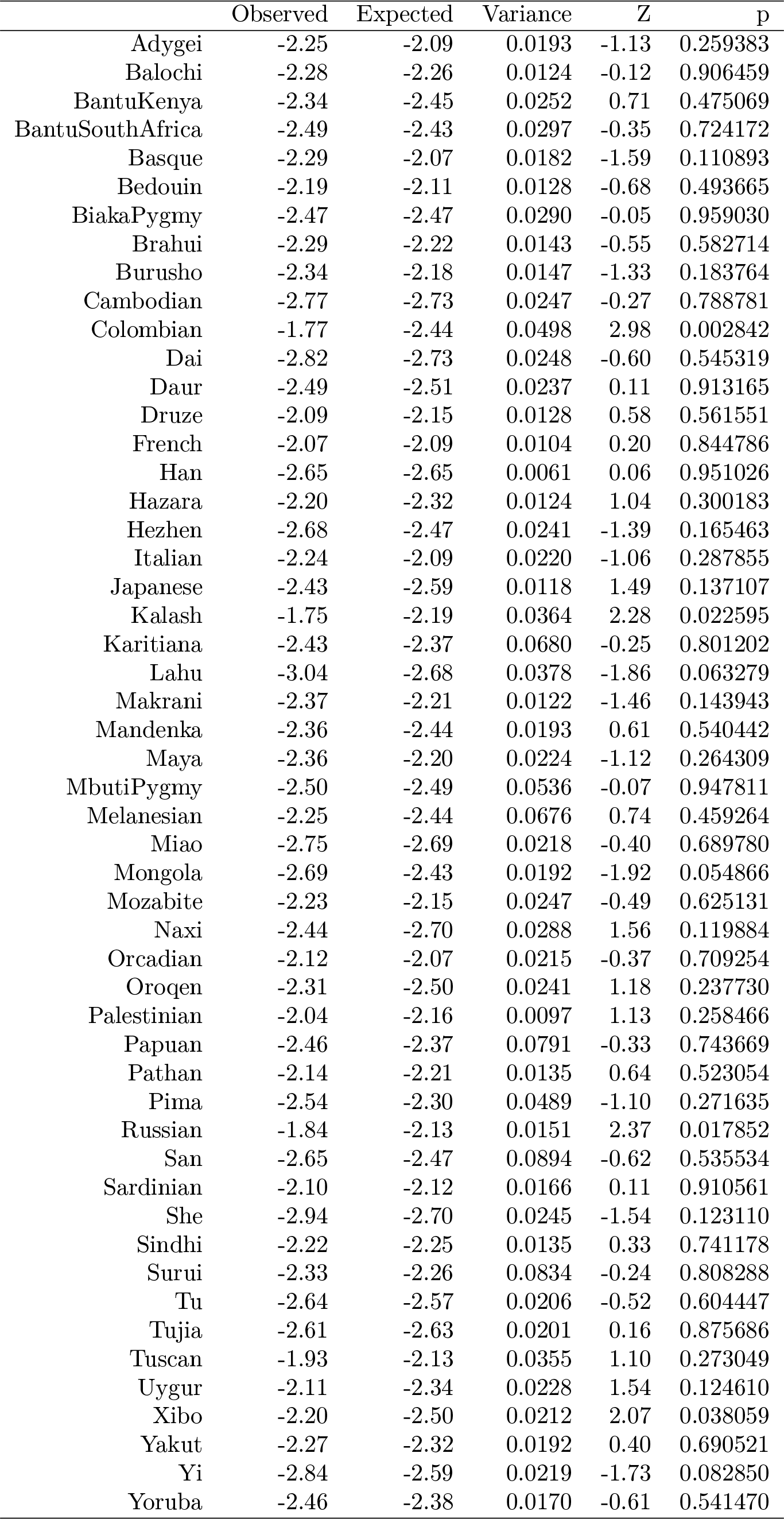
Conditional analysis at the individual population level for the BMI dataset

**Table S9:**
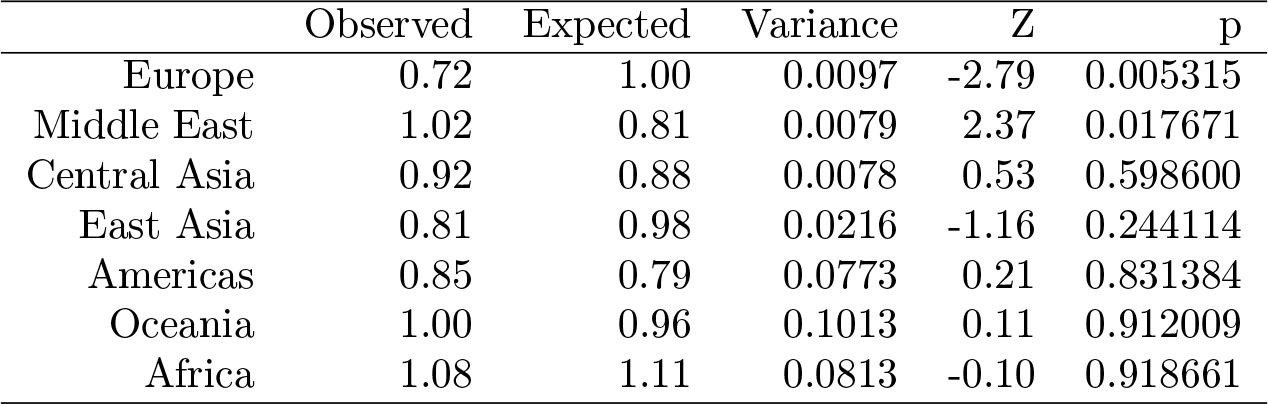
Conditional analysis at the regional level for the T2D dataset.

**Table S10:**
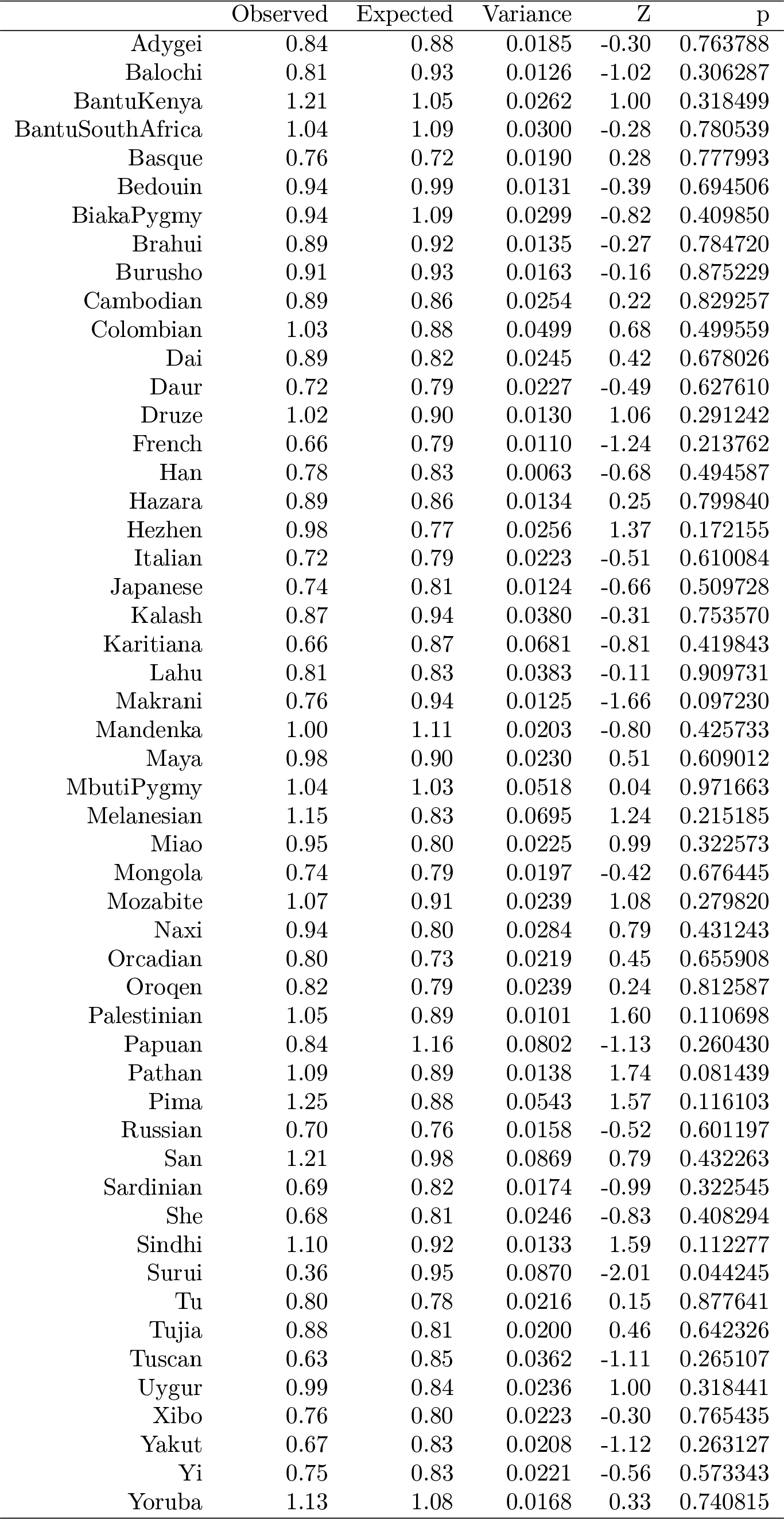
Conditional analysis at the individual population level for the T2D dataset.

**Table S11:**
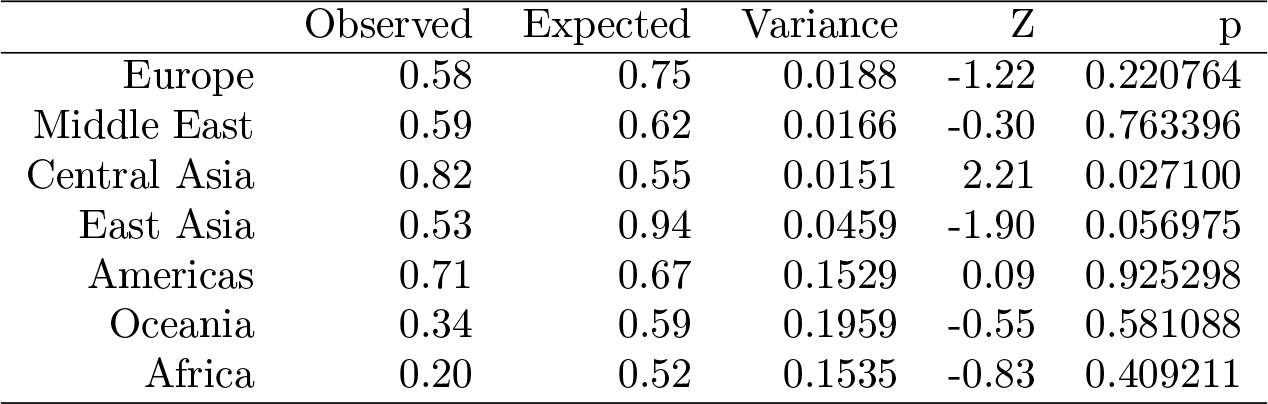
Conditional analysis at the regional level for the CD dataset.

**Table S12:**
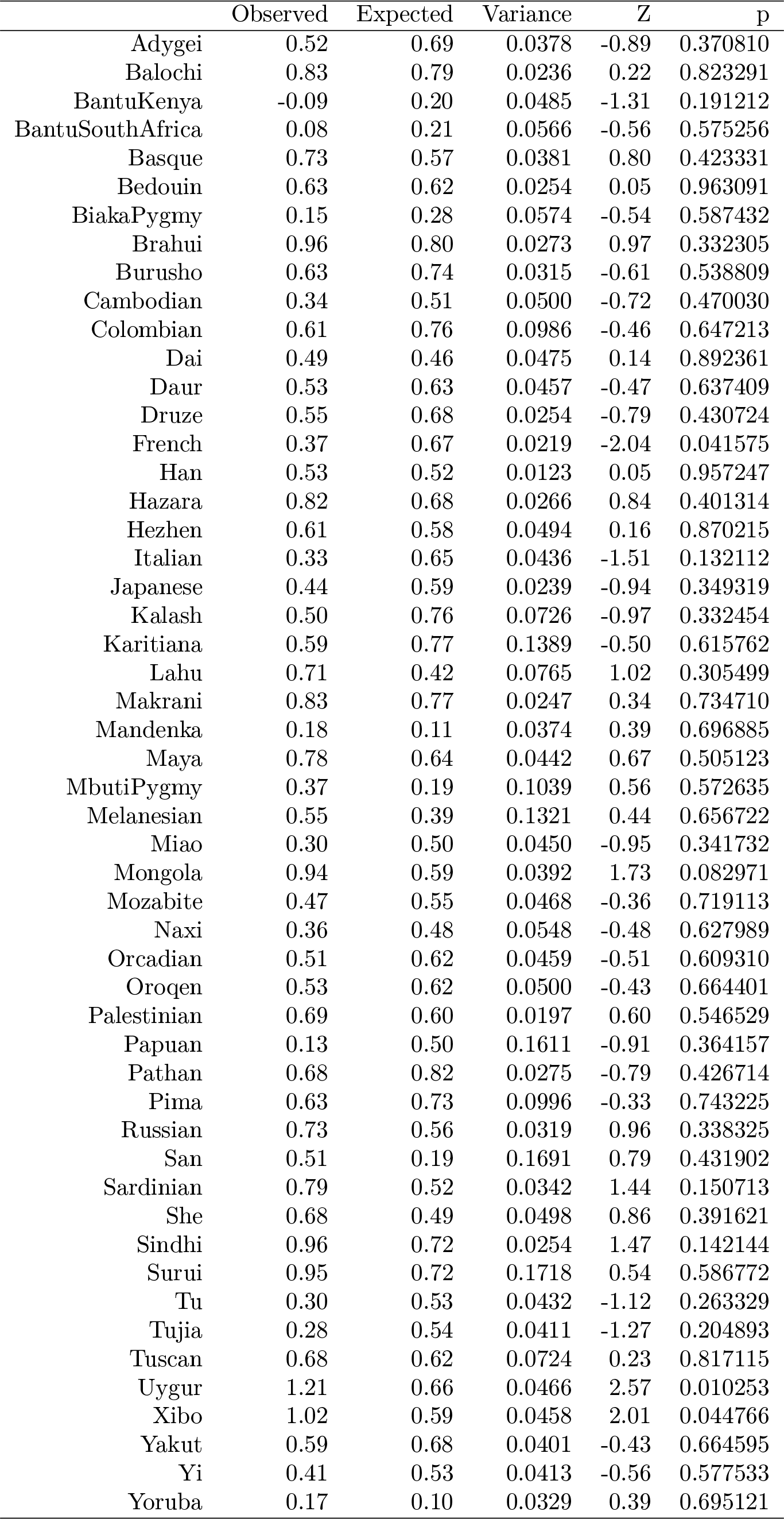
Conditional analysis at the individual population level for the CD dataset.

**Table S13:**
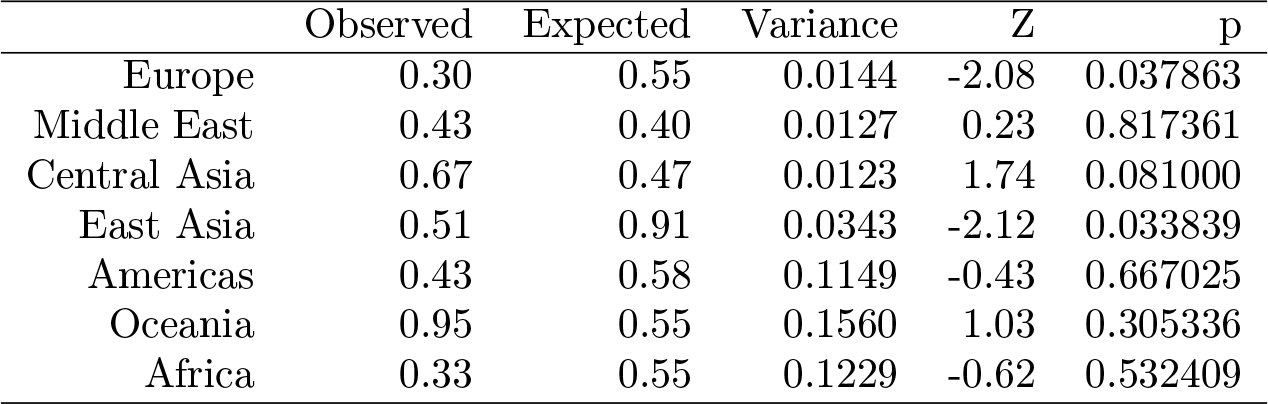
Conditional analysis at the regional level for the UC dataset.

**Table S14:**
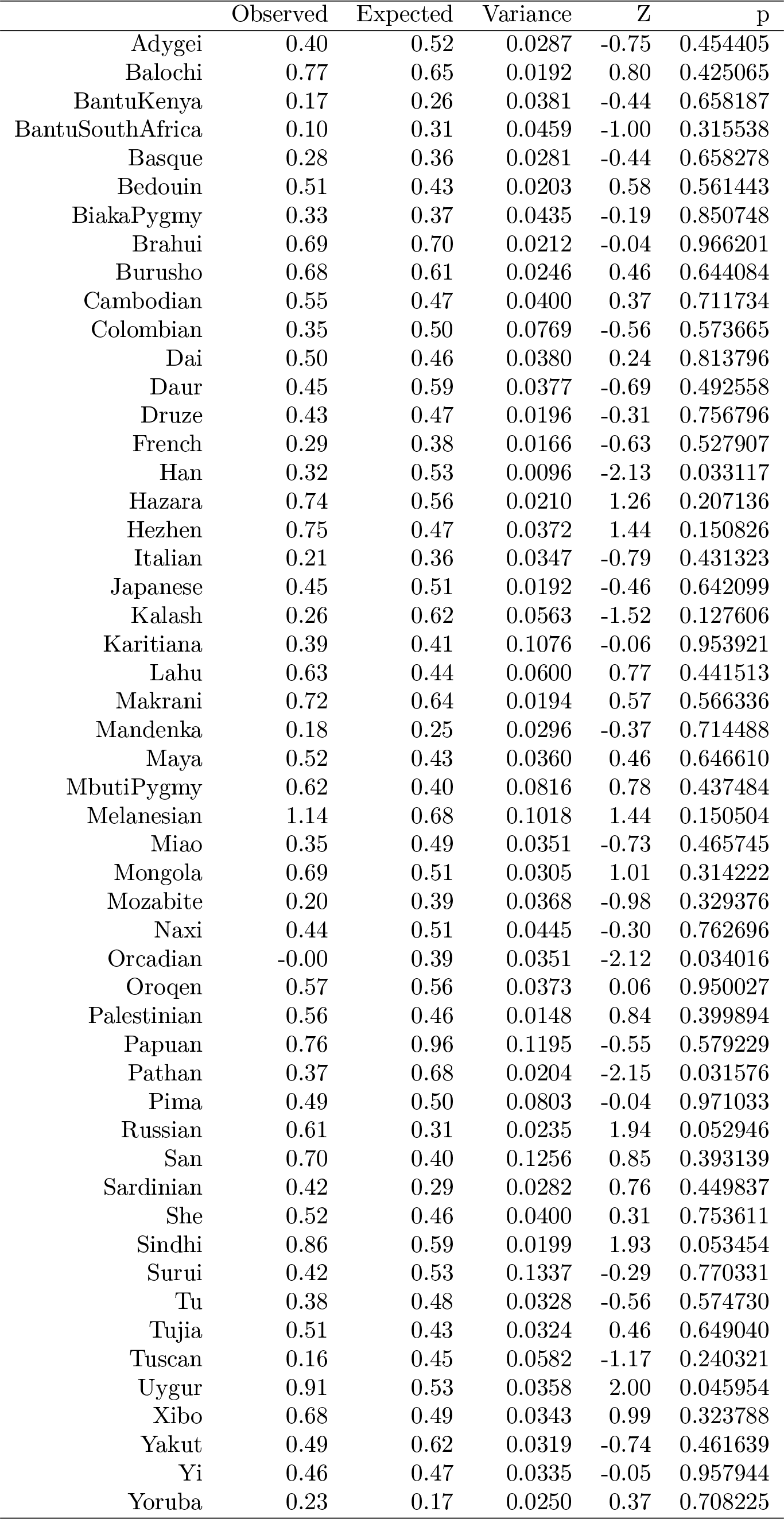
Conditional analysis at the individual population level for the UC dataset

**Table S15:** Corresponding *β* statistics for all analyses presented in 2.

| Phenotype | SUMPC1 | SUMPC2 | WINPC1 | WINPC2 | Latitude |
| --- | --- | --- | --- | --- | --- |
| Height | -0.22 (0.11) | 0.003 (0.93) | -0.12 (0.37) | **0.38 (0.008)** | 0.12 (0.34) |
| Skin Pigmentation | **0.31(0.025)** | 0.07 (0.62) | 0.27 (0.061) | -0.11 (0.38) | **-0.36 (0.009)** |
| Body Mass Index | -0.22 (0.15) | 0.04 (0.74) | -0.17 (0.29) | 0.24 (0.11) | 0.21 (0.21) |
| Type 2 Diabetes | 0.10 (0.36) | 0.10 (0.42) | 0.14 (0.25) | -0.07 (0.56) | -0.20 (0.093) |
| Crohn's Disease | 0.25 (0.066) | **-0.30 (0.033)** | 0.01 (0.86) | -0.28 (0.053) | 0.08 (0.61) |
| Ulcerative Colitis | 0.17 (0.22) | **-0.29 (0.046)** | 0.06 (0.61) | -0.21 (0.15) | -0.11(0.40) |

**Table S16:**
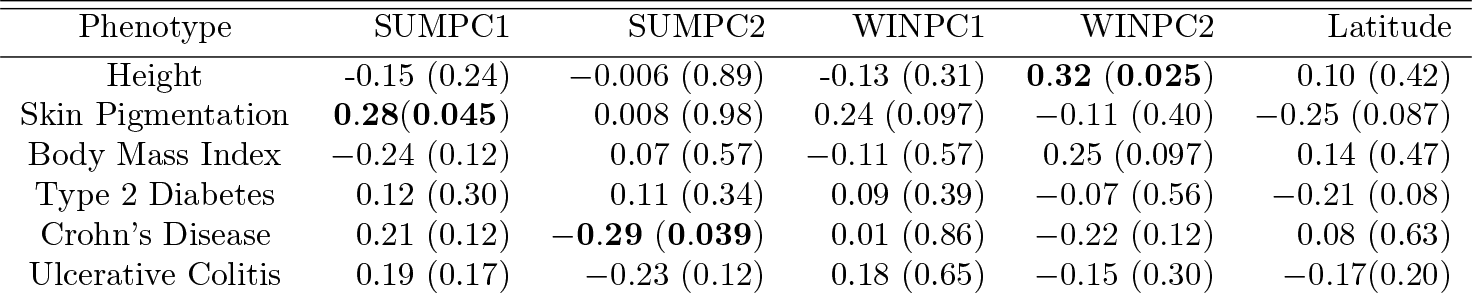
Corresponding *ρ* statistics for all analyses presented in 2.

| Phenotype | SUMPC1 | SUMPC2 | WINPC1 | WINPC2 | Latitude |
| --- | --- | --- | --- | --- | --- |
| Height | -0.15 (0.24) | -0.006 (0.89) | -0.13 (0.31) | **0.32 (0.025)** | 0.10 (0.42) |
| Skin Pigmentation | **0.28(0.045)** | 0.008 (0.98) | 0.24 (0.097) | -0.11 (0.40) | -0.25 (0.087) |
| Body Mass Index | -0.24 (0.12) | 0.07 (0.57) | -0.11 (0.57) | 0.25 (0.097) | 0.14 (0.47) |
| Type 2 Diabetes | 0.12 (0.30) | 0.11 (0.34) | 0.09 (0.39) | -0.07 (0.56) | -0.21 (0.08) |
| Crohn's Disease | 0.21 (0.12) | **-0.29 (0.039)** | 0.01 (0.86) | -0.22 (0.12) | 0.08 (0.63) |
| Ulcerative Colitis | 0.19 (0.17) | -0.23 (0.12) | 0.18 (0.65) | -0.15 (0.30) | -0.17(0.20) |

